# Insulin resistance alters the coupling between cerebral blood flow and glucose metabolism in younger and older adults: Implications for neurovascular coupling

**DOI:** 10.1101/2024.02.07.579389

**Authors:** H.A. Deery, E. Liang, R. Di Paolo, K. Voigt, G. Murray, M.N. Siddiqui, G.F. Egan, C. Moran, S.D. Jamadar

**Author notes:** **Corresponding author address:** Monash Biomedical Imaging, Monash University, 770 Blackburn Rd, Melbourne, 3800, Australia. **e-mail address:**.

## Abstract

Rising rates of insulin resistance and an ageing population are set to exact an increasing toll on individuals and society. Here we examine the contribution of insulin resistance and age to the coupling of cerebral blood flow and glucose metabolism; a critical process in the supply of energy for the brain. Thirty-four younger (20-42 years) and 41 older (66-86 years) healthy adults underwent a simultaneous resting state MR/PET scan, including arterial spin labelling. Rates of cerebral blood flow and glucose metabolism were derived using a functional atlas of 100 brain regions. Older adults had lower cerebral blood flow than younger adults in 95 regions, reducing to 36 regions after controlling for cortical atrophy and blood pressure. Younger and older insulin sensitive adults showed small, negative correlations between relatively high rates of regional cerebral blood flow and glucose metabolism. This pattern was inverted in insulin resistant older adults, who showed hypoperfusion and hypometabolism across the cortex, and a positive coupling. In insulin resistant younger adults, coupling showed inversion to positive correlations, although not to the extent seen in older adults. Our findings suggest that the normal course of ageing and insulin resistance alter the rates and coupling of cerebral blood flow and metabolism. They underscore the criticality of insulin sensitivity to brain health across the adult lifespan.

## Introduction

One of society’s most pressing challenges is dealing with the health, social and economic costs of non-communicable or “lifestyle diseases” [1, 2]. Historically, conditions such as obesity, insulin resistance and type 2 diabetes, have been associated with older age and economic development, but they are increasingly prevalent in younger adults and low- and middle-income countries [3]. People with these conditions are at increased risk for a range of other health issues, including cognitive decline [4], making their prevention and management priorities for governments and health authorities around the world [2].

Due to the limited ability of the brain to store energy, it is susceptible to factors that influence real-time energy supply. One such factor is insulin. Insulin mediates the uptake of glucose into cells and supports glucose homeostasis. Some people are more resistant to the action of insulin than others. Lifestyle conditions and associated behaviours, such as low physical activity and poor diet, are associated with increased resistance to insulin’s action [5]. Clinical and sub-clinical levels of insulin resistance are associated with changes to the brain [6]. For example, insulin resistance is a signature of Alzheimer’s disease and type 2 diabetes and is associated with deleterious changes to the brain’s structure, vasculature and metabolism (for reviews see [4, 7]). Increased grey matter atrophy and cognitive decline have been found with higher levels of insulin resistance, even in the absence of comorbidities [8, 9]. Elevated glycaemia in otherwise healthy individuals in their 20s to 40s can cause grey matter atrophy, reduced white matter integrity and impaired cognition [10-12].

In the human brain, a tight coupling is required between neuronal activity and cerebral blood flow (CBF) for the delivery of oxygenated blood, glucose and other nutrients to neurons [13]. While differences in the absolute rates of CBF and glucose metabolism have been well studied in ageing and disease, less is known about their coupling. For example, older adults have lower absolute CBF and CMR_GLC_ than younger adults [14-16], particularly in the frontal and temporal regions [15, 17]. Lower rates of CBF have been associated with increased glycemia, insulin resistance and adiposity in otherwise healthy middle aged and older adults, [18-23]. The links between insulin resistance and reduced cerebral metabolic rates of glucose (CMR_GLC_) are also well established (see [6], for review). In recent work, we demonstrated that older adults had lower CMR_GLC_ than younger adults, but insulin resistance significantly attenuated this association in younger adults only [24].

Insulin appears to influence CMR_GLC_, and CBF coupling (see [25] for a review). For example, insulin modulates brain perfusion via vasoactive effects [26, 27] and controls coupling of brain glucose and blood flow via astrocytic receptors [28, 29]. In terms of age differences in CBF and CMR_GCL_ coupling, Bentourkia et al. reported that older people had lower CBF and CMR_GLC_ than younger people, but the coupling between CBF and CMR_GLC_ was similar in both groups [30]. In contrast, Henriksen et al. found coupling between CBF and CMR_GLC_ at the regional but not the whole brain level in older adults [31].

In the current study we examined the rates and coupling of cerebral blood flow and glucose metabolism in healthy older and younger adults and explored the contribution of insulin resistance and age to this coupling. We hypothesised that older age and higher insulin resistance are associated with lower CBF, and the effects of older age and insulin resistance are mediated by cortical thickness and blood pressure. We hypothesised that the coupling of CBF and CMR_GLC_ is stronger in younger than older adults. We further hypothesised that CBF and CMR_GLC_ coupling is strongest in those with the least insulin resistance and weakest in those with the most insulin resistance. Finally, we hypothesised that there are spatial differences in the coupling for older versus younger adults and for levels of insulin resistance.

## Results

### Demographic Factors

There was no statistically significant difference in sex, years of education or body mass index between the older and younger adult groups (Table 1). However, older adults had lower whole brain cortical thickness and higher systolic blood pressure (p < .001) and resting heart rate (p < .05) compared to younger adults. Although participants did not have a previous diagnosis of hypertension upon study entry, nine younger and 26 older participants would meet the definition for hypertension from the measurements taken, i.e., a systolic blood pressure of 140 mmHg or higher, a diastolic blood pressure 90 mmHg or higher, or both [32].

**Table 1.** Demographics comparison of younger and older groups and four sub-groups based on age category and HOMA-IR median split. Continuous variables are mean (standard deviation); categorical variables are %.

|  | Age Category: Younger vs Older |  |  | 4 Groups: Age Category and HOMA-IR Median Split |  |  |  |  |
| --- | --- | --- | --- | --- | --- | --- | --- | --- |
|  | Younger<br>(N = 34) | Older<br>(N = 41) | P<sup>1</sup> | Younger Insulin<br>Sensitive<br>(n = 14) | Younger Insulin<br>Resistant<br>(n = 20) | Older Insulin<br>Sensitive<br>(n = 25) | Older Insulin<br>Resistant<br>(n = 16) | P<sup>1</sup> |
| Age (years) | 28.2 (6.3) | 75.7 (5.8) | < .001 | 29.6 (6.1) | 27.2 (6.3) | 75.3 (6) | 76.3 (5.5) | < .001 |
| Sex - Number of Females (%) | 19 (56%) | 18 (44%) | .302 | 9 (64%) | 10 (50%) | 14 (56%) | 4 (25%) | 0.139 |
| Years of Education | 18.2 (2.9) | 16.9 (3.7) | .125 | 19.2 (2.8) | 17.4 (2.7) | 16.1 (2.9) | 17.8 (4.5) | 0.088 |
| Insulin (mU/L) | 4.8 (3.2) | 4.1 (2.8) | .280 | 2.2 (1.0) | 6.6 (2.9) | 2.3 (0.7) | 6.7 (2.6) | < .001 |
| Blood glucose (mmol/L) | 4.7 (0.4) | 5.2 (0.6) | < .001 | 4.6 (0.4) | 4.8 (0.3) | 4.8 (0.4) | 5.6 (0.4) | < .001 |
| HOMA-IR | 1.0 (0.7) | 1.0 (0.8) | .821 | 0.4 (0.2) | 1.4 (0.6) | 0.5 (0.1) | 1.7 (0.7) | < .001 |
| HOMA-IR2 | 0.61 (0.41) | 0.54 (0.38) | .431 | 0.2 (0.1) | 0.8 (0.3) | 0.3 (0.1) | 0.9 (0.3) | < .001 |
| Systolic Blood Pressure (mmHg) | 120.1 (17.8) | 148.3 (24.6) | < .001 | 113.3 (16.3) | 124.8 (17.7) | 151.0 (28.0) | 144.0 (18.0) | < .001 |
| Diastolic Blood Pressure (mmHg) | 79.6 (13.7) | 83.3 (12.1) | .218 | 73.9 (10.2) | 83.6 (14.6) | 86.8 (12.5) | 77.8 (9.2) | 0.010 |
| Resting Heart Rate (BPM) | 81.8 (17.2) | 73.8 (13) | .026 | 75.2 (16.8) | 86.3 (16.2) | 75.7 (13.0) | 70.8 (12.6) | 0.015 |
| Body Mass Index (BMI) | 24.2 (4.8) | 25.7 (3.5) | .127 | 22.7 (3.1) | 25.2 (5.5) | 24.7 (3.6) | 27.2 (2.7) | 0.029 |
| Cortical Thickness (mm) | 2.52 (0.08) | 2.36 (0.11) | < .001 | 2.5 (0.07) | 2.5 (0.07) | 2.3 (0.09) | 2.3 (0.13) | < .001 |

<sup>1</sup>P-values are based on ANOVA for continuous and Chi-square for categorical variables.

Significant differences were found between the four groups based on age category and insulin resistance levels on all demographics except sex and years of education (see Table 1). For the insulin sensitive younger and older groups, the mean HOMA-IR was 0.4 and 0.5, respectively; whereas for insulin resistant younger and older groups it was 1.4 and 1.7, respectively. A clinical diagnosis of insulin resistance is not usually made from HOMA-IR, however, thresholds have been proposed in the literature ranging from 1.8 in to 2.5 [33]. Four and five participants in the younger and older insulin resistant groups would meet a threshold of 1.8 HOMA-IR, respectively; and three and four participants a HOMA-IR threshold of 2.5 (see Table S2). A higher percentage of older adults than younger adults in both the high and low HOMA-IR groups would also meet the threshold for hypertension.

Because of the known impact of blood pressure and the other demographic factors on the brain, we ran additional GLMs with them as covariates to test the coupling of CBF with CMR_GLC_ (see Supplement, Tables S4 and S5, and Figures S2 and S3).

### The Effect of Age, Cortical Thickness and Blood Pressure on Regional CBF

Older age was associated with lower CBF in 95 regions (p-FDR < .05; Figure 1A and Table S1). Older adults had lower CBF than younger adults in 85 regions (p-FDR < .05) when cortical thickness was added to the models (Figure 1B). When cortical thickness and systolic and diastolic blood pressure were included, older adults had lower CBF in 36 regions (Figure 1C). The largest age differences in CBF were in the bilateral temporal poles (.282 to .317), temporal cortices (.246), ventral, orbital, lateral, medial and dorsal prefrontal cortices (.203 to .327), superior parietal lobule (.210), and insula (.201). Adding cortical thickness had a minimal impact on both the magnitude of the effect sizes and the regions showing the largest age differences in CBF (Table S3). However, the effect sizes reduced by approximately 40% to 60% in those same regions when systolic and diastolic blood pressure were also added to the models.

**Figure 1.**
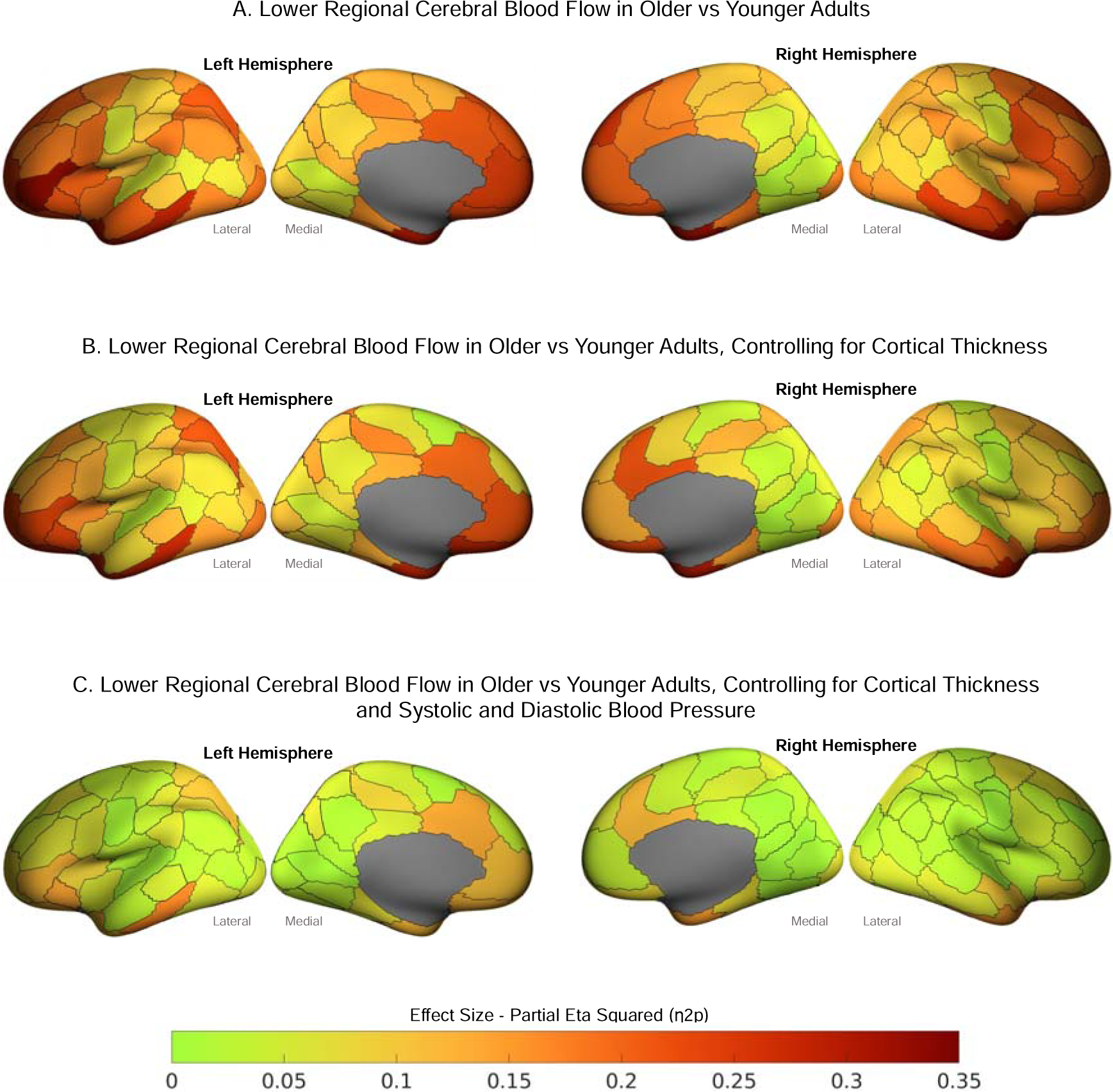
Effect sizes (η2p) from general linear models showing lower regional cerebral blood flow in older vs younger adults. Cerebral blood flow is significantly lower in older than younger adults in (A) 95 regions, (B) 85 regions when cortical thickness was added as a covariate, and (C) 36 regions when cortical thickness and systolic and diastolic blood pressure were included as covariates. See Table S1 for details of GLMs.

### The Effect of Age and Insulin Resistance on Regional CBF

A significant difference was found in CBF in 40 regions between the four groups based on age category and insulin resistance levels (see Figure 2 and Table S2). Post-hoc contrast showed that older insulin resistant adults had lower CBF than younger insulin sensitive adults. This pattern was found in regions within all networks, except the somatomotor network. The effect size was the largest in the posterior cingulate (.265 to .185), precuneus (.156), temporal cortex (.114), and lateral prefrontal cortex (.119 to .186) in the control network; the post central (.112 to .114) and superior parietal lobule (.171) in the dorsal attention network; the medial and dorsal prefrontal cortices (.110 to .209) and ventral prefrontal cortices (.132 to .186) in the default network; the insula (.115), parietal medial (.152) and medial posterior prefrontal cortices (.174 to .207) in the salience ventral attention; and the orbital frontal (.134 to .167) and temporal poles (.189 to .212) in the limbic network.

**Figure 2.**
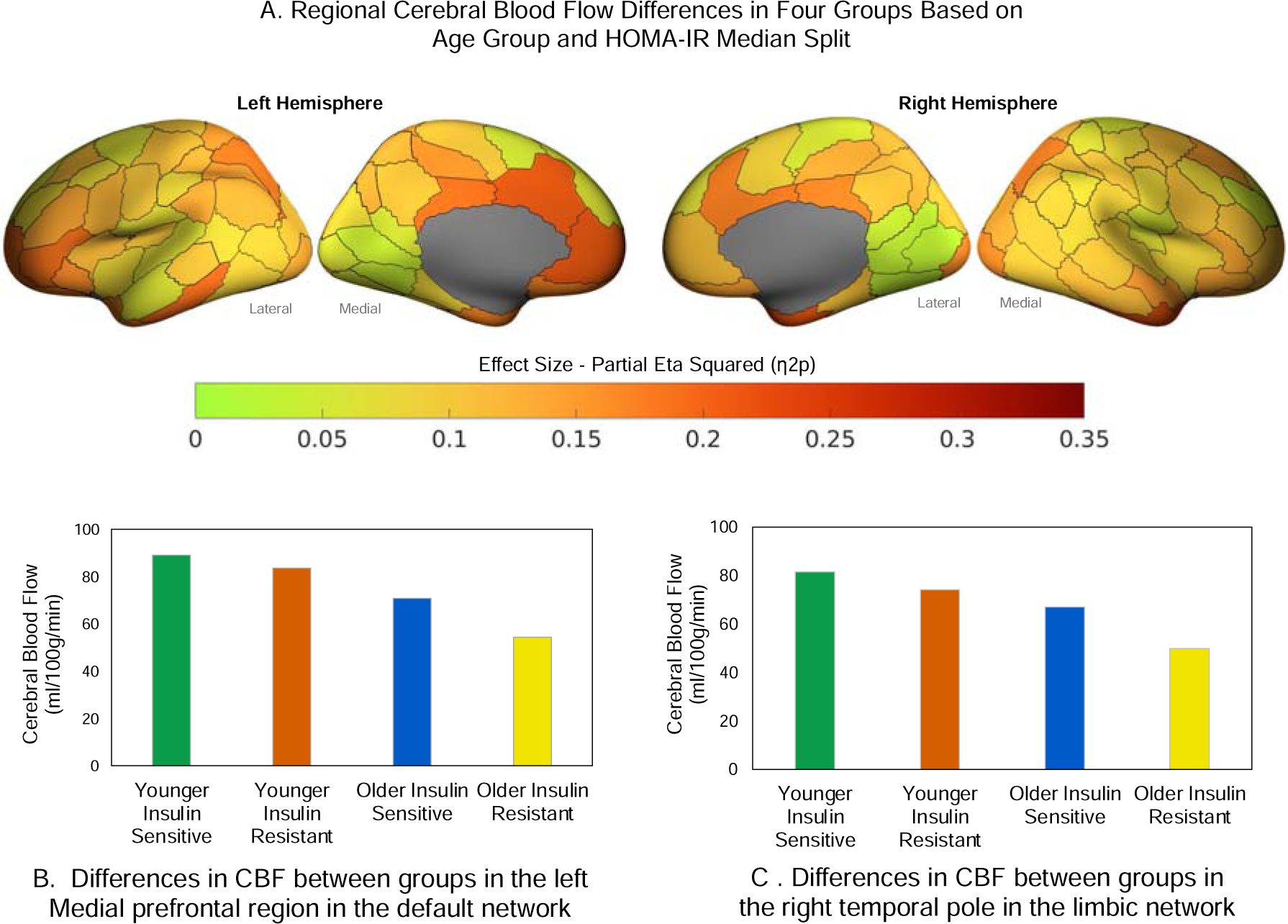
(A) Effect sizes (η2p) from general linear models of regional cerebral blood flow differences among four groups based on age group and HOMA-IR median split, controlling for cortical thickness and blood pressure. Example CBF group differences in the (B) left medial prefrontal cortex in the default network, and (C) right temporal pole in the limbic network. See Table S3 for details of GLMs.

When the four groups were derived from age group and a HOMA-IR2 median split (younger low/high HOMA-IR2, older low/high HOMA-IR2), we found minimal to no differences in results to those based on HOMA-IR (see Supplement Table S3). This was expected as HOMA-IR2 models increases in the insulin secretion curve for plasma glucose concentrations above 10 mmol/L [34]; a threshold that than none of the participants reached. Hence, the remainder of the results are based on analyses of HOMA-IR.

### The Effect of Age and HOMA-IR on the Coupling of CBF and CMR_GLC_

When differences in coupling were examined for the four groups based on age category and insulin resistance levels, we found widespread between-group differences at both the network (Figure 3) and regional level (Figure 4 and Figure S1). Older insulin resistant adults had strong positive correlations between CBF and CMR_GLC_ in all networks. The highest correlations were in the somatomotor and salience ventral attention networks (Figure 3), followed by the dorsal attention, default and control network. For younger insulin resistant adults, the correlations between CBF and CMR_GLC_ were mostly small-to-moderate positive in all networks and much lower than for the older insulin resistant adults. In contrast, both younger and older insulin sensitive adults showed small-to-moderate negative correlations between CBF and CMR_GLC_ across the networks.

**Figure 3.**
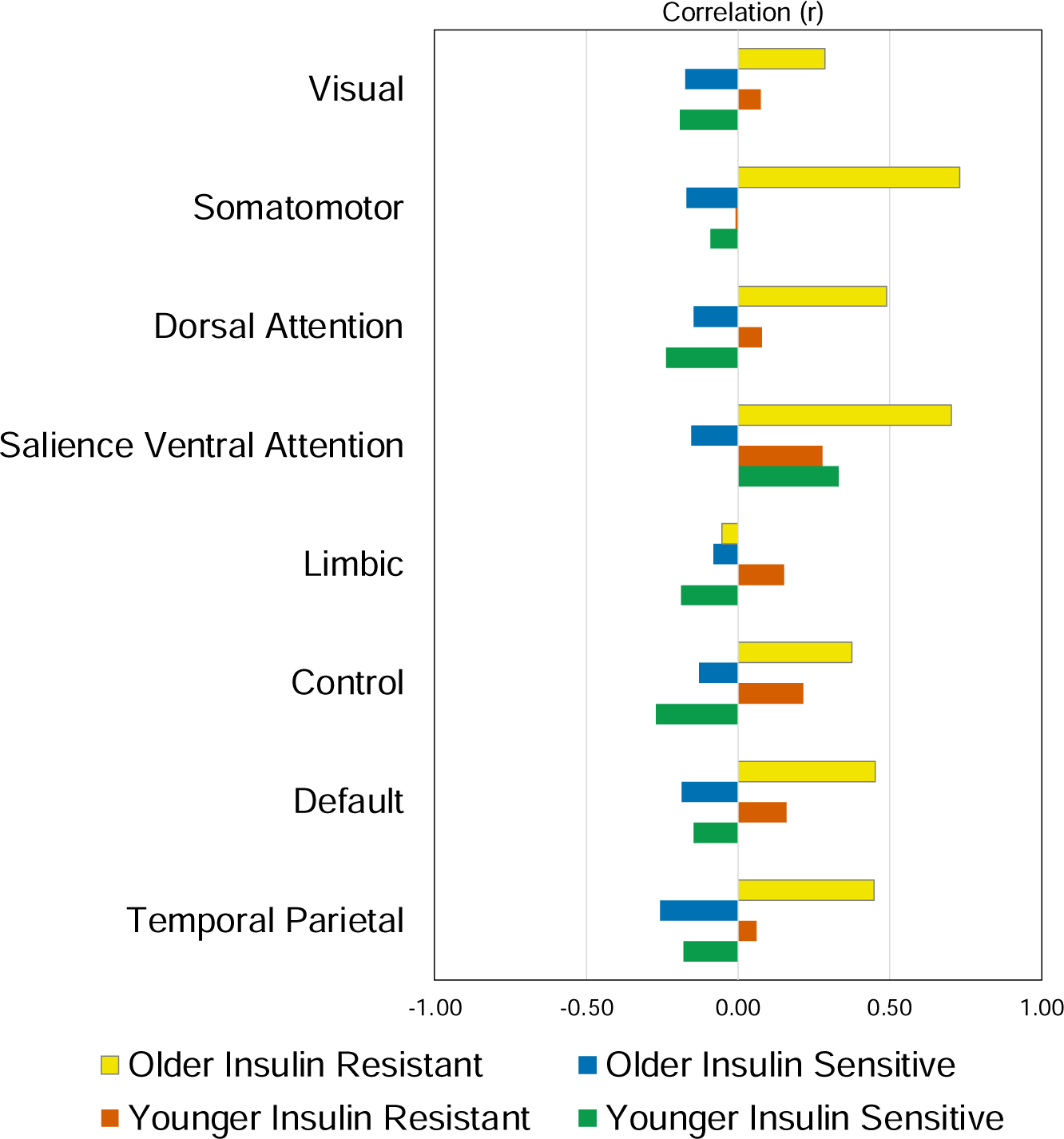
Partial correlation of cerebral blood flow (CBF) and glucose metabolism (CMR_GLC_) controlling for cortical thickness and systolic and diastolic blood pressure in the Schaefer networks for four groups: younger insulin sensitive; younger insulin resistant; older insulin sensitive; and older insulin resistant.

**Figure 4.**
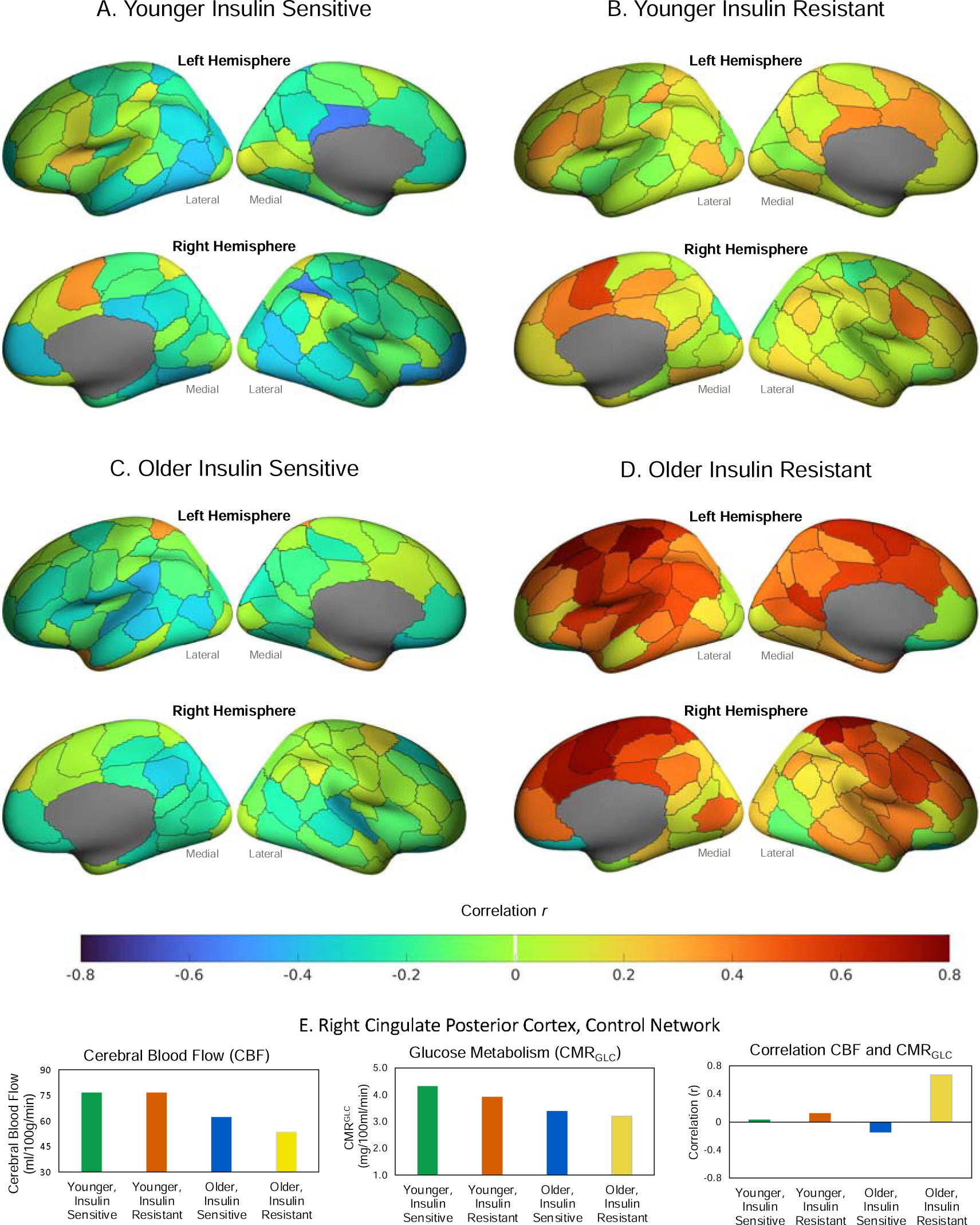
Partial correlation of regional cerebral blood flow (CBF) and glucose metabolism (CMR_GLC_) controlling for cortical thickness and systolic and diastolic blood pressure for four groups: Younger insulin sensitive (A), younger insulin resistant (B), older insulin sensitive and (C) older insulin resistant (D). Example results for the four groups in the right cingulate posterior cortex for absolute CBF and CMR_GLC_ and the CBF-CMR_GLC_ correlation (E).

The pattern of CBF and CMR_GLC_ coupling described above at the network level was reflected across many of the regions (Figure 4 and Figure S1). Younger and older insulin sensitive adults showed primarily negative correlations, with 79 and 73 total negative correlations, and 50 and 38 correlations below -.2, respectively. The pattern was inverted in older insulin resistant adults, with positive correlations between CBF and CMR_GLC_ in 94 regions (79 region had an r > .2). Older insulin resistant adults also showed the lowest absolute rates of CBF and CMR_GLC_ among all groups (see Figure 4E for an example). Younger insulin resistant adults showed smaller positive correlations in 82 regions (25 with r > .2).

The Pearson and Spearman correlations assessing the strength and rank of the CBF and CMR_GLC_ coupling across the 100 regions between the younger insulin sensitive and older insulin resistant groups were both significant (r = .31 and r = .27, p < .01). The DICE coefficient was 0.42 when comparing the top 50% of regional correlations and 0.24 for the top 25% of absolute regional correlations between younger insulin sensitive and older insulin resistant groups.

The Pearson correlations approached significance for the young insulin sensitive and resistant groups (r = .12, p < .060). The DICE coefficient was .48 and .54 comparing the top 50% of regional correlations between the younger insulin sensitive and younger insulin resistant and older insulin resistant groups, respectively. For the top 25% of regions, the DICE coefficients were .16 and .28 for comparison of the younger insulin sensitive group with the younger insulin resistant and older insulin sensitive groups, respectively.

### The Effect of Age and HOMA-IR on the Coupling of CBF and fMRI Graph Metrics

The GLM was significant for the coupling of CBF and local efficiency (Table 2). Significant differences were found between the four groups based on age category and insulin resistance levels (.182), and diastolic blood pressure (.064). One sample T-tests showed that the correlation between CBF and local efficiency was significantly different to zero for all groups (p< .001), except the older insulin resistant adults. Post-hoc contrasts for the group effect from the GLM showed that younger insulin sensitive adults had significantly higher negative correlations between CBF and local efficiency than both older adult groups (Figure 5), that is, higher cerebral blood flow was more strongly associated with lower local network efficiency among insulin sensitive younger adults than insulin sensitive older adults. For older insulin resistant adults, the correlation was both significantly different to younger insulin sensitive adults and weakly positive.

**Figure 5.**
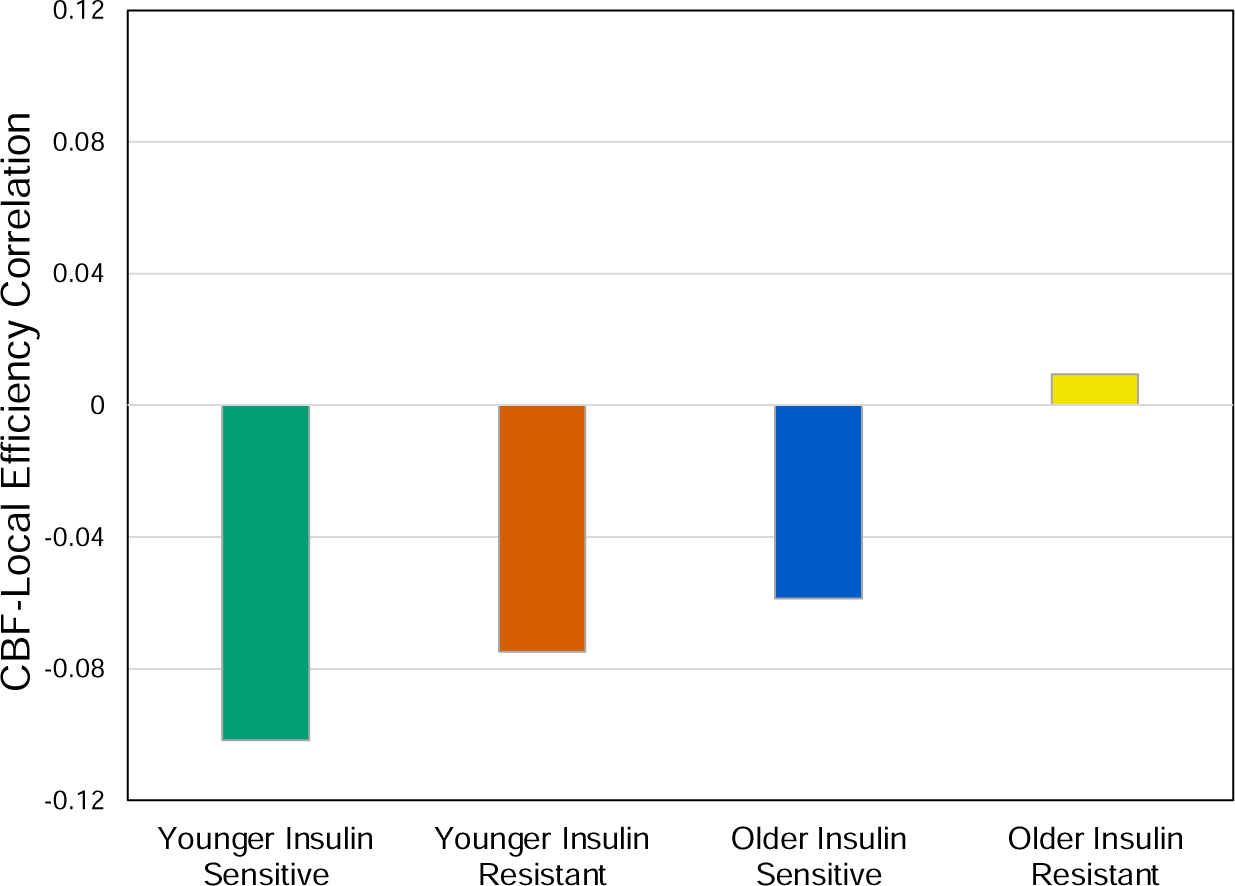
Significant CBF and local efficiency correlation for four groups based on age group and HOMA-IR median slit. Post hoc contrasts are significant for younger insulin sensitive vs older insulin sensitive and older insulin resistant groups (see Table 2). One sample T-tests also show that the correlation for younger insulin sensitive, younger insulin resistant and older insulin sensitive adults are significantly different to zero (all p < .05).

**Table 2.** General linear models of coupling of CBF and fMRI graph metrics (individual correlations across the 100 regions). The group effect for local efficiency is shown in Figure 5.

|  | Overall |  |  | 4 Groups: Age Category and HOMA<br>Median Split |  |  |  |  |  | Systolic BP |  |  | Diastolic BP |  |  | Cortical Thickness |  |  |
| --- | --- | --- | --- | --- | --- | --- | --- | --- | --- | --- | --- | --- | --- | --- | --- | --- | --- | --- |
| | F | p | $\eta^2_p$ | F | p | $\eta^2_p$ | YIS: vs<br>YIR | YIS: vs<br>OIS | YIS: vs<br>OIR | F | p | $\eta^2_p$ | F | p | $\eta^2_p$ | F | p | $\eta^2_p$ |
| Correlation CBF and Global Efficiency | 1.9 | 0.091 | 0.150 | 0.1 | 0.982 | 0.003 |  |  |  | 0.4 | 0.537 | 0.006 | 0.3 | 0.613 | 0.004 | 6.5 | 0.013 | 0.090 |
| Correlation CBF and Local Efficiency | 4.3 | 0.001 | 0.285 | 4.8 | 0.004 | 0.182 | 0.161 | 0.011 | 0.000 | 0.2 | 0.645 | 0.003 | 4.5 | 0.039 | 0.064 | 3.9 | 0.053 | 0.056 |
| Correlation CBF and Betwenness Centrality | 1.7 | 0.135 | 0.136 | 1.1 | 0.359 | 0.048 |  |  |  | 0.0 | 0.998 | 0.000 | 1.7 | 0.197 | 0.025 | 0.2 | 0.659 | 0.003 |

YIS = younger insulin sensitive; YIR = younger insulin resistant; OIS = older insulin sensitive; OIR = older insulin resistant

## Discussion

Consistent with our first hypothesis, we found lower cerebral blood flow in older than younger adults in 95% of brain regions. The percentage of regions showing significant age differences reduced to 85% when lower cortical thickness among the older adults was taken into account. These results confirm research suggesting that the older adult brain has lower blood flow on a per tissue basis, particularly in the frontal and temporal regions [14, 16, 17], and highlight the importance of adjusting for cortical atrophy when examining age differences in cerebral blood flow. After also adjusting for blood pressure differences, the percentage of regions showing lower age-related cerebral blood reduced to 36%. These results are consistent with previous research [35-40] suggesting that elevated blood pressure in ageing is an important contributor to cerebral hypoperfusion. We discuss the possible mechanisms driving these results below.

The results partly support our second hypothesis that higher insulin resistance is associated with lower CBF. Older insulin resistant adults had lower CBF than younger insulin sensitive adults in 40 regions, but CBF was not significantly lower in younger insulin resistant adults. Older insulin resistant adults also had lower absolute CBF than younger insulin resistant adults. Taken together, these results indicate that insulin resistance attenuates age-related hypoperfusion in older adults. Our results add to a growing body of research showing that insulin modulates brain perfusion [26, 27, 41] and indicate that increased insulin resistance is associated with lower cerebral blood flow in older adults without Type 2 diabetes or clinical levels of insulin resistance. The fact that we found age group differences after adjusting for blood pressure differences suggests that insulin resistance is also exerting an influence on regional cerebral blood flow in ageing that is independent of blood pressure.

We found support for our third hypothesis of altered cerebral blood flow and glucose metabolism coupling in older and insulin resistant adults. Older people with insulin resistance had a very different pattern of coupling between CBF and CMR_GLC_ than the insulin sensitive groups, suggesting a failure of the coupling. Older insulin resistant adults showed a pattern of moderate (r > .2), positive correlations in 79% of regions. We recently reported the relationship between CMR_GLC_, age and insulin resistance in the same sample used in the current study [24]. Older adults had lower absolute regional CMR_GLC_ than younger adults, and insulin resistance had a small, non-significant negative effect on CMR_GLC_ in older adults. For the older insulin resistant adults, the positive coupling of CBF and CMR_GLC_ in the current study appears to be an outcome of normal age related hypoperfusion and hypometabolism, as well as an additional insulin resistant-driven reduction in blood flow. Together these results suggest that the positive coupling in older insulin resistant adults is a response to an “energy crisis” of low absolute levels of blood flow and rates of glucose metabolism. Such an “energy crisis” is possibly an early pathological event in age-related neurodegenerative disease [42].

The regional coupling in younger and older insulin sensitive adults showed mostly small-to-moderate negative correlations. These negative correlations may allow for the maintenance of metabolic homeostasis within a relatively tight physiological range. In other words, to maintain the supply of blood carrying O_2_ and glucose, blood flow and glucose metabolic rates increase in compensation for a decrease in the other and vice versa. For example, if cerebral rates of glucose metabolism drop, blood flow is increased to ensure the supply of additional blood carrying metabolic substrates.

In the same sample as the current study, we recently showed a significant attenuation of regional CMR_GLC_ in younger insulin resistant adults [24]. Coupling of CBF and CMR_GLC_ in the insulin resistant younger adults appears to show early signs of changes, although not to the extent seen in older insulin resistant adults (only 25% regions showed an inversion of positive correlations > .2, compared to 79% in older adults). These results raise the possibility that for younger insulin resistant adults, progression of their insulin resistance may result in a phenotype resembling older insulin resistant adults, even in mid-life. In other words, insulin resistance may accelerate “normal” age-related reductions in CMR_GLC_ and its coupling with cerebral blood flow. Additional research across the spectrum of insulin resistance and the adult lifespan could test this hypothesis and further characterise the effects.

There were low topological similarities among the groups in terms of the regions showing the strongest coupling of cerebral blood flow and metabolism. The highest 25% of correlations between CBF and CMR_GLC_ among the insulin resistant older adults ranged from 0.62 to 0.88. Fifteen of the 25 regions (60%) were in default, control and salience ventral attention networks, including regions in the prefrontal and medial frontal cortices, the insula and superior parietal lobule. Seven regions (28%) were in the somatomotor cortex. These results suggest that the positive coupling in response to cerebral hypoperfusion and hypometabolism in older insulin resistant adults is prioritised to the prefrontal and somatomotor cortices to support sensory and motor processing and executive functions, such as decision making, attention and cognitive control [43]. They also raise the possibility that reduced supply of metabolic substrates is contributing to age-related impairments widely reported in executive function [44].

We found that insulin resistance affects the strength and direction of coupling between cerebral blood flow and local network efficiency. In insulin sensitive individuals, higher cerebral blood flow was associated with lower local network efficiency, particularly in younger adults. However, the correlation was reduced in strength for older insulin sensitive adults and weakly positive for older insulin resistant adults. Separate areas of research have shown that older age is associated with an increase in insulin resistance [45, 46] and a loss of local network efficiency [47]. This study is the first to examine these effects within the same cohort and the results indicate that age-related increases in insulin resistance likely impact the efficiency of local network communication via cerebral blood flow.

Although we did not directly assess the mechanisms driving CBF alterations in ageing, changes to the physiology of the cerebrovascular in older adults have been well described. For example, age related cerebrovascular damage causes a loss of elasticity and thickening of the arterial walls, impaired cerebrovascular reactivity, damage to the blood brain barrier and ischemia [48-50]. Changes to larger arteries progresses to distal and downstream damage of the microvasculature and the neurovascular unit, which in turn leads to reduced efficiency in neural processing [50]. These changes can also impact cognition due to reduced oxygen and glucose and an impaired ability of the microvasculature to respond to neuronal metabolic demands [51].

Our results are consistent with research showing that elevated blood pressure reduces CBF and alters neurovascular coupling [52, 53]. Narrowing of major cerebral arteries in the early stages of hypertension may protect downstream vessels from elevated pressure but lead to eventual stenosis of the arteries and decreased downstream blood supply [54]. Hypertension also alters autoregulation of blood flow [55]. We found that elevated systolic blood pressure was more strongly associated with lower CBF than elevated diastolic blood pressure, whereas diastolic blood pressure was more important in CBF and local network efficiency coupling. This later result is consistent with that seen in cortical thickness. In Gutteridge et al., we found that long-term variability in diastolic, and not systolic, blood pressure was an important influence on cortical thickness in older age [56]. Of note, in the current study we obtained a single measure of blood pressure at a single timepoint, whereas in Gutteridge et al. variability in blood pressure over three years was examined. On balance, the evidence indicates that both diastolic and systolic blood pressure are important mediators of cerebral blood flow and brain health in older age [57, 58] and suggest that lifestyle and antihypertensive medications that reduce blood pressure may support cognitive health among ageing adults at risk for hypertension (see [59], for discussion of previous trials).

Disrupted neurovascular coupling is seen in diabetes and other diseases in which insulin resistance and cardiometabolic dysfunction play a role, such as dementia (see [16, 17, 60] for reviews). Mechanistically, insulin resistance and Type 2 diabetes have been shown to affect the cells types (e.g., neurons, astrocytes, microglia), pathways and blood flow underpinning neurovascular coupling [50, 61, 62]. Our findings suggest that these effects are not confined to adults who meet clinical thresholds for insulin resistance. Rather, the coupling of blood flow with both glucose metabolism and local efficiency is affected in insulin resistant adults who have not progressed to a clinical diagnosis. Maintaining insulin sensitivity and cerebrovascular health through a healthy lifestyle could support cognitive ageing. A focus on exercise, sleep, stress and nutrition could lead to improvements in cerebrovascular and metabolic health and cognitive outcomes in ageing [50].

The current study used a cross-sectional design, limiting conclusions about the causal relationship between age, insulin resistance and cerebral blood flow. It is also possible that the differences we found between groups reflects unmeasured differences in the cohorts rather than age-related changes. Research measuring changes in CBF and CMR_GLC_ longitudinally could help confirm the causal pathways and the magnitude of yearly or decade-long reductions in CBF and their impact on coupling and cognition. Such research may also help explain the time course and relative impact of mediating factors, such as blood pressure. Research also suggests that CBF alterations across the lifespan may follow a non-linear trajectory. CBF undergoes a rapid drop in the second decade, remains stable until approximately the fifth decade, before gradually declining in older age [63]. Due to the absence of “middle aged” adults in the current study, non-linear relationships could not be tested but could be a topic for future research.

Although our results are informative in terms of coupling between CBF and CMR_GLC_ at the regional level and over relatively long timescales, additional research assessing these factors at other spatial and temporal resolutions is needed. In particular, investigating the relationship between neurovascular coupling and insulin resistance at higher spatial and temporal resolutions could provide important insights into brain health in ageing and disease (see [64], for a review of methods). Our age and insulin resistance groups also had relatively small sample sizes and our the results require replication.

In conclusion, older adults show hypoperfusion and hypometabolism across the cortex. Cerebral blood flow is further reduced in ageing with peripheral insulin resistance and the presence of hypertension. For older insulin resistant adults, the coupling of cerebral blood flow and metabolism is strongly positive, particularly in the prefrontal and somatomotor cortices. This positive coupling possibly reflects a response to an under supply of metabolic substrates and an effort to support the brain’s high energy demands. Rates of glucose metabolism and blood flow are higher in younger adults. However, insulin resistance in younger adults attenuates cerebral glucose metabolism. Coupling in younger insulin resistant adults shows early signs of inversion to positive correlations, although not yet to the extent seen in older insulin resistant adults. “Normal” age-related changes and/or progression in insulin resistance are likely to further alter blood flow and coupling and place younger insulin resistant individuals at risk for cognitive decline. Our findings underscore the criticality of insulin sensitivity to brain health across the adult lifespan and the importance of maintaining cerebrovascular and metabolic health to support cognitive function.

## Method

Full details of the methods are provided in the Supplement.

### Participants

Participants were recruited from the general community via local advertising. The final sample included 75 individuals, 34 younger (mean 28.2; SD 6.3; range 20-42 years) and 41 older (mean 75.7; SD 5.8; range 66-86 years) adults (see Table 1). Exclusion criteria included a known diagnosis or history of hypertension, diabetes, neurological or psychiatric illness.

### Demographic Variables

Prior to the scan, participants completed an online demographic and lifestyle questionnaire including age, sex, education, height and weight.

### Simultaneous MR-PET Data Acquisitions

Participants underwent a 90-minute simultaneous MR-PET scan in a Siemens (Erlangen) Biograph 3-Tesla molecular MR scanner (for scan parameters see Supplementary Information). At the beginning of the scan, half of the 260 MBq FDG tracer was administered via the left forearm as a bolus, the remaining 130 MBq of the FDG tracer dose was infused at a rate of 36ml/hour over 50 minutes [65, 66]. Non-functional MRI scans were acquired during the first 12 minutes, including a T1 3DMPRAGE and T2 FLAIR. Thirteen minutes into the scan, list-mode PET and T2* EPI BOLD-EPI sequences were initiated. A 40-minute resting-state scan was undertaken in naturalistic viewing conditions. At 53 minutes, a 5-delay pseudo-continuous arterial spin labelling (pCASL) scan was undertaken [67]. Plasma radioactivity levels were measured every 10 minutes throughout the scan.

### MRI/PET Processing

Full details of the pre-processing steps and calculations of MRI and PET measures are provided in the Supplement. Briefly, single−subject whole−brain CBF maps were calculated from perfusions weighted images in BASIL. Processing included motion correction, distortion correction with field map, and partial volume correction. The resulting CBF images were aligned to the anatomical T1w images, normalised to MNI152 space and parcellated using the Schaefer functional atlas of 100 regions and seven networks [68]. For the structural T1 images, the brain was extracted in Freesurfer, manually checked and registered to MNI152 space. Regional cortical thickness was extracted from the Freesurfer reconstruction statistics.

For the BOLD-fMRI data, T2* images were brain extracted (FSL BET), unwarped and motion corrected (FSL MCFLIRT), temporally detrended, normalised to MNI space and smoothed at 8mm FWHM [69]. The images were checked for excessive head motion (see Supplement for details). The BOLD timeseries data was loaded to CONN [70] to generate graph metrics from the Schaefer 100 regions, with thresholding of the matrices at the highest 40% of edges. Global efficiency, local efficiency and betweenness centrality were used.

The list-mode PET data for each subject was binned into 344 3D sinogram frames of 16s intervals and corrected for attenuation (see Supplement for details). The reconstructed DICOM slices were converted to NIFTI format, temporally concatenated to a 4D NIFTI volume, and motion corrected (FSL MCFLIRT; [69]). The images were corrected for partial volume effects using the modified Müller-Gartner method. Calculation of regional CMR_GLC_ was undertaken in PMOD 4.4 (see Supplement).

### Insulin resistance

The baseline blood sample was used to collect 2ml of plasma for insulin and glucose measurement, which was undertaken by a commercial laboratory. HOMA-IR was calculated as fasting glucose (mmol/L) x fasting insulin (mU/L) / 22.5 [71]. The constant of 22.5 is a normalising factor for normal fasting plasma insulin and glucose (i.e., 4.5 mmol/L x 5 mU/L = 22.5). Higher HOMA-IR values indicate greater insulin resistance. We also calculated HOMA-IR2 using the calculator at https://www.rdm.ox.ac.uk/.

### Data Analysis

All analyses were run in SPSS version 29.0 (https://www.ibm.com/products/spss-statistics).

#### Demographics

For the demographic variables, independent T-tests for age group differences in continuous and Chi-square tests for categorical variables were undertaken and tested at uncorrected p < 0.5.

Four groups were created based on age category (younger and older) and a HOMA-IR median split (low and high HOMA-IR, or insulin sensitive and insulin resistant): younger insulin sensitive, younger insulin resistant, older insulin sensitive and older insulin resistant. Group differences in the demographics variables were tested using general linear models (GLMs).

#### The Effect of Age, Cortical Thickness and Blood Pressure on Regional CBF

Three series of GLMs was run to assess whether regional CBF was lower in older versus younger adults and whether the effect of age group was mediated by differences in cortical thickness and blood pressure. In the first series of GLMs, the association between regional CBF and age group was tested. In the second series, cortical thickness was included in the models as a covariate; in the third series, cortical thickness and systolic and diastolic blood pressure were also added.

#### The Effect of Age and HOMA-IR on Regional CBF

Differences in regional CBF were compared among the four groups based on age category and HOMA-IR median split in a series of GLMs. Regional cortical thickness and systolic and diastolic blood pressure were included as covariates. For regions showing significant group differences, post-hoc contrasts were run comparing the younger insulin sensitive group to the other three groups. A similar series of analyses were run for four groups based on age category and HOMA-IR2 median split.

Each series of GLMs was FDR-corrected at p < .05 for the overall model [72].

#### Coupling of Regional and Network CBF and CMR_GLC_

Differences in CBF and CMR_GLC_ coupling were compared among the four groups based on age category and levels of insulin resistance. A separate series of partial correlations was calculated at the seven network and 100 region levels for each of the four groups, controlling for cortical thickness at the same level as well as systolic and diastolic blood pressure. At the region level, Pearson correlations, Spearman correlations and DICE coefficients were calculated comparing the strength, rank and spatial similarity of the coupling for the younger insulin sensitive group to each of the other three groups.

#### Effect of Age and HOMA-IR on the Coupling of CBF and Functional Brain Network Properties

For each participant, a Pearson’s correlation was computed between CBF and global and local efficiency and betweenness centrality across the 100 regions. The correlations were considered a measure of coupling of CBF and functional network properties and used as the dependent variables in separate GLMs. Differences between the four groups based on age category and insulin resistance levels were tested, together with whole brain cortical thickness and systolic and diastolic blood pressure as covariates.

## Data Availability Statement

The datasets used and/or analyzed during the current study available from the corresponding author on reasonable request.

## Author Contributions

S.D.J. and G.F.E., conceived the project. H.D., S.D.J., C.M., and G.F.E. designed and developed the manuscript. H.D. and E.L. analysed the data. H.D. wrote the manuscript, S.D.J., G.F.E. and C.M. reviewed/edited the manuscript. G.M., N.M.S., K.V., and H.D. collected the data. All authors read and approved the final manuscript.

## Other Information

The authors declare no conflicts of interest.

