## Supplementary material for "Insulin resistance alters the coupling between cerebral blood flow and glucose metabolism in younger and older adults: Implications for neurovascular coupling": Supplementary Materials_Deery et al.pdf

**~ Supplementary Information ~**

**Supplementary Methods**

### *1. Ethical Considerations*

The study protocol was reviewed and approved by the Monash University Human Research Ethics Committee in accordance with Australian Code for the Responsible Conduct of Research (2007) and the Australian National Statement on Ethical Conduct in Human Research (2007). Administration of ionizing radiation was approved by the Monash Health Principal Medical Physicist, following the Australian Radiation Protection and Nuclear Safety Agency Code of Practice (2005). For participants older than 18 years, the annual radiation exposure limit of 5 mSv applies. The effective dose in this study was 4.9 mSv.

### *2. Recruitment*

Ninety participants were recruited from the general community via local advertising. An initial screening interview ensured that participants had the capacity to provide informed consent, did not have a history of hypertension, a diagnosis of diabetes, neurological or psychiatric illness, and were not taking psychoactive medication that could affect cognitive function or metabolism. Participants were also screened for claustrophobia, non-MR compatible implants, and clinical or research PET scan in the past 12 months. Women were screened for current or suspected pregnancy. Participants received a $100 voucher for participating in the study.

Fifteen participants were excluded from further analyses due to blood haemolysis or well counter issues preventing insulin measurement or kinetic modelling (n=7), excessive head motion (n=2) or incomplete PET (n=2) or ASL scans (N = 2). Four participants were excluded due to consistently high or low ASL or $CMR_{GLC}$ values more than 2.5 standard deviations from the mean.

### *3. MR-PET Data Acquisition*

Participants underwent a 90-minute simultaneous MR-PET scan in a Siemens (Erlangen) Biograph 3-Tesla molecular MR scanner. Participants were directed to consume a high-protein/low-sugar diet for the 24 hours prior to the scan. They were also instructed to fast for six hours and to drink 2–6 glasses of water. Prior to FDG infusion, participants were cannulated in the vein in each forearm and a 10ml baseline blood sample taken. At the beginning of the scan, half of the 260 MBq FDG tracer was administered via the left forearm as a bolus, providing a strong PET signal from the beginning of the scan. The remaining 130 MBq of the FDG tracer dose was infused at a rate of 36ml/hour over 50 minutes, minimising the amount of signal decay over the course of the data acquisition. We have previously demonstrated that this protocol provides a good balance between a fast increase in signal-to-noise ratio at the start of the scan, and maintenance of signal-to-noise ratio over the duration of the scan [1].

Participants were positioned supine in the scanner bore with their head in a 32-channel radiofrequency head coil and were instructed to lie as still as possible. The scan sequence was as follows. Non-functional MRI scans were acquired during the first 12 minutes, including a T1 3DMPRAGE (TA = 3.49 min, TR = 1640ms, TE = 234ms, flip angle = 8°, field of view = 256 × 256 mm2, voxel size = 1.0 × 1.0 × 1.0 mm3, 176 slices, sagittal acquisition) and T2 FLAIR (TA = 5.52 min, TR = 5,000ms, TE = 396ms, field of view = 250 × 250 mm2, voxel size = .5 × .5 × 1 mm3, 160 slices) to image the anatomical grey and white matter structures, respectively. Thirteen minutes into the scan, list-mode PET (voxel size = 2.3 x 2.3 x 5.0mm3) and T2* EPI BOLD-fMRI (TA = 40 minutes; TR = 1000ms, TE = 39ms, FOV = 210 mm2, 2.4 × 2.4 × 2.4 mm3 voxels, 64 slices, ascending axial acquisition) sequences were initiated. A 40-minute resting-state scan was undertaken in naturalistic viewing conditions watching a movie of a drone flying over the Hawaii Islands. At 53 minutes, a 5-delay pseudo-continuous arterial spin labelling (pCASL) scan was undertaken. Scan parameters were TR = 4,220 ms; TE = 45.46 ms; FOV = 240 mm; slice thickness = 3 mm; voxel size 2.5 x 2.5 x 3.0 mm. PLDs were 0.5, 1, 1.5, 2, and 2.5 s, duration of the labelling pulse was 1.51s. At 58 minutes, diffusion-weighted imaging (DWI) was acquired with 71 directions to index white matter connectivity. DWI results are not reported here.

Plasma radioactivity levels were measured throughout the duration of the scan. Beginning at 10-minutes post infusion onset, 5ml blood samples were taken from the right forearm using a vacutainer at 10-minute intervals for a total of nine samples. The blood sample were immediately placed in a Heraeus Megafuge 16 centrifuge (ThermoFisher Scientific, Osterode, Germany) and spun at 2,000 rpm (RCF ~ 515g) for 5 minutes. 1,000-μL plasma was pipetted, transferred to a counting tube, and placed in a well counter for four minutes. The count start time, total number of counts, and counts per minute were recorded for each sample.

#### *4. MRI Pre-Processing, Cerebral Blood Flow, Cortical Thickness and Graph Theory Metrics*

To quantify the ASL signal, the BASIL toolkit of the Oxford Centre for Functional MRI of the BRAIN (FMRIB)'s software library (FSL) was used (https://fsl.fmrib.ox.ac.uk/fsl/fslwiki/BASIL). A calibration map M0 of proton density weighted image was acquired for each participant. Single−subject whole−brain CBF maps were calculated from perfusions weighted images (direct subtraction of label and control volumes) in BASIL. Processing included motion correction, distortion correction with field map and partial volume correction. The model included a macro vascular component, adaptive spatial regularisation of perfusion, and incorporating T1 uncertainty. The arterial transit time was set at 1.3s, T1/T1b at 1.3/1.66s, and inversion efficiency at 0.85. The resulting CBF images in native space were aligned to the anatomical T1w images and normalised to MNI152 space with Advanced Normalization Tools (ANTs).

For the structural T1 images, the brain was extracted in Freesurfer; and the quality of the pial/white matter surface was manually checked and corrected. Corrected Freesurfer surfaces were registered to MNI152 space using ANTs. Cortical thickness for the Schaefer 100 regions were obtained from the Freesurfer reconstruction statistics for each participant. Freesurfer calculates cortical thickness as the closest distance from the grey and white matter boundary to the grey matter and cerebrospinal fluid boundary at each vertex [2, 3].

For the BOLD-fMRI data, T2* images were brain extracted (FSL BET), unwarped and motion corrected with six rotation and translation parameters (FSL MCFLIRT), temporally detrended, normalised to MNI space and smoothed at 8mm FWHM (Jenkinson et al., 2002). Framewise displacement was calculated for each participant to check for excessive head motion (mean and % of frames > .3mm). The pre-processed BOLD timeseries data was loaded to the CONN toolbox [4] and denoised by regression of white matter and CSF confounds. The timeseries data was bandpass frequency filtered between 0.01 Hz and 0.1 Hz. Regions of interest were generated using the Schaefer 100 parcellations [5]. Graph metrics were derived after thresholding the resulting matrices at the highest 40% of edges to describe the topological properties of the entire network [6]. Global efficiency, local efficiency and betweenness centrality were used, as follows:

**Global Efficiency** at a node is defined as the average of the shortest inverse-distances between the node and all other nodes in the graph. Across the entire graph, global efficiency represents a measure of global integration.

**Local Efficiency** at each node is defined as the average of shortest inverse-distances between the nodes within the neighbouring sub-graph (all nodes neighbouring that node and all existing edges among them). Network local efficiency is a measure of local integration of a network.

**Betweenness Centrality** is defined as the proportion of times that a node is part of a shortest-path between any two pairs of nodes within a graph. It represents an alternative measure of node centrality within a graph.

### *5. PET Image Reconstruction, Pre-Processing and $CMR_{GLC}$ calculations*

The list-mode PET data for each subject was binned into 344 3D sinogram frames of 16s intervals. Attenuation was corrected via the pseudo-CT method for hybrid PET-MR scanners [7] Ordinary Poisson-Ordered Subset Expectation Maximization algorithm (3 iterations, 21 subsets) with point spread function correction was used to reconstruct 3D volumes from the sinogram frames. The reconstructed DICOM slices were converted to NIFTI format with size 344 × 344 × 127 (voxel size: 2.09 × 2.09 × 2.03 mm3) for each volume. All 3D volumes were temporally concatenated to form a single 4D NIFTI volume. After concatenation, the PET volumes were motion corrected using FSL MCFLIRT [8], with the mean PET image used to mask the 4D data. PET images were corrected for partial volume effects using the modified Müller-Gartner method.

Calculation of $CMR_{GLC}$ was undertaken in PMOD 4.4 (http://www.pmod.com) using the FDG time activity curves for the Schaefer 100 parcellation. The FDG in the plasma samples was decay-corrected for the time between sampling and counting and used as the input function to the Patlak models. A lumped constant of 0.89 was used, and equilibrium (t) set at 10 mins, the time corresponding to the peak of the bolus and onset of a stable signal [1]. The fractional blood space (vB) was set at 0.05 [9]. Participant's plasma glucose (mmol) was entered in the model from their baseline blood sample.

#### *6. Correction for Partial Volume Effects*

PET images were corrected for partial volume effects using the modified Müller-Gartner method implemented in PetSurf (https://surfer.nmr.mgh.harvard.edu/fswiki/PetSurfer). The method corrects for white matter spill in and grey matter spill out of the PET signal [10, 11]. The equation subtracts from the grey matter voxel signal the white matter signal (convoluted by the point spread function) and divides by the grey matter signal. This division can introduce over-correction at the grey matter boundary, and hence a grey matter binary mask is recommended, with the threshold level needing to be chosen. A grey matter threshold of 20-30% is recommended in ageing because atrophy can influence results [10]. For our analyses, we chose a 25% grey matter threshold and surface-based spatial smoothing [10, 11]. We used a Gaussian kernel with a full width at half maximum of 12 mm to increase the signal-to-noise ratio. Subcortical structures were partial volume corrected and spatially smoothed in volume space and merged with the cortical data.

### Supplementary Results

Table S1. General liner models of the association of regional cerebral blood flow with age category (model 1), age category and cortical thickness (model 2) and age category, cortical thickness and blood pressure (model 3). The age category effect sizes from each model are plotted on the brain surface in Figure 1 in main document.

| | Model 1: Age Category | | | Model 2: Age Category and Cortical Thickness | | | | | | Model 3: Age Category, Cortical Thickness, Systolic and Diastolic Blood Pressure | | | | | |
|---|---|---|---|---|---|---|---|---|---|---|---|---|---|---|---|
| | Age Category | | | Overall Model | | | Age Category | | | Overall Model | | | Age Category | | |
| | F | p-FDR | $\eta^2_p$ | F | p | $\eta^2_p$ | F | p | $\eta^2_p$ | F | p | $\eta^2_p$ | F | p | $\eta^2_p$ |
| Visual Central: Extra Striate Cortex 1 | 7.3 | 0.011 | 0.093 | 3.6 | 0.032 | 0.093 | 6.4 | 0.013 | 0.083 | 2.9 | 0.030 | 0.142 | 2.2 | 0.139 | 0.031 |
| Visual Central: Extra Striate Cortex 2 | 16.1 | 0.000 | 0.183 | 8.1 | 0.001 | 0.186 | 14.8 | 0.000 | 0.173 | 7.3 | 0.000 | 0.297 | 3.8 | 0.056 | 0.052 |
| Visual Central: Striate Cortex 1 | 7.8 | 0.009 | 0.097 | 5.0 | 0.009 | 0.123 | 8.5 | 0.005 | 0.107 | 4.3 | 0.004 | 0.199 | 1.5 | 0.232 | 0.021 |
| Visual Central: Extra Striate Cortex 3 | 11.2 | 0.002 | 0.135 | 5.6 | 0.006 | 0.136 | 8.8 | 0.004 | 0.111 | 4.2 | 0.004 | 0.195 | 2.1 | 0.152 | 0.030 |
| Visual Peripheral: Extra Striate Inferior 1 | 4.7 | 0.036 | 0.062 | 2.5 | 0.088 | 0.066 | 4.9 | 0.030 | 0.064 | 1.5 | 0.216 | 0.079 | 1.7 | 0.197 | 0.024 |
| Visual Peripheral: Striate Cortex Calcarine 1 | 3.3 | 0.078 | 0.043 | 1.7 | 0.198 | 0.045 | 3.2 | 0.078 | 0.043 | 1.4 | 0.243 | 0.075 | 0.5 | 0.504 | 0.006 |
| Visual Peripheral: Extra Striate Cortex Sup 1 | 7.7 | 0.009 | 0.097 | 3.8 | 0.026 | 0.097 | 5.9 | 0.017 | 0.077 | 2.5 | 0.050 | 0.127 | 1.2 | 0.274 | 0.017 |
| Somatomotor A: 1 | 7.1 | 0.012 | 0.090 | 3.7 | 0.030 | 0.094 | 4.8 | 0.032 | 0.063 | 2.2 | 0.077 | 0.113 | 2.1 | 0.151 | 0.030 |
| Somatomotor A: 2 | 10.1 | 0.004 | 0.123 | 5.1 | 0.009 | 0.125 | 5.2 | 0.025 | 0.068 | 2.6 | 0.046 | 0.129 | 3.3 | 0.073 | 0.046 |
| Somatomotor B: Auditory 1 | 4.3 | 0.046 | 0.056 | 2.1 | 0.125 | 0.057 | 2.3 | 0.138 | 0.031 | 1.5 | 0.202 | 0.082 | 0.5 | 0.497 | 0.007 |
| Somatomotor B: S2 1 | 9.0 | 0.005 | 0.112 | 4.5 | 0.015 | 0.112 | 6.2 | 0.015 | 0.080 | 3.3 | 0.016 | 0.160 | 1.1 | 0.288 | 0.016 |
| Somatomotor B: S2 2 | 8.2 | 0.008 | 0.102 | 4.3 | 0.018 | 0.107 | 7.1 | 0.009 | 0.091 | 2.6 | 0.042 | 0.132 | 2.3 | 0.133 | 0.032 |
| Somatomotor B: Central 1 | 4.5 | 0.040 | 0.059 | 2.2 | 0.113 | 0.059 | 3.7 | 0.059 | 0.049 | 1.8 | 0.131 | 0.096 | 0.6 | 0.445 | 0.008 |
| Dorsal Attention A: Temporal Occipital 1 | 14.5 | 0.001 | 0.168 | 7.2 | 0.001 | 0.169 | 12.1 | 0.001 | 0.146 | 5.2 | 0.001 | 0.231 | 4.3 | 0.042 | 0.058 |
| Dorsal Attention A: Parietal Occipital 1 | 5.6 | 0.023 | 0.073 | 3.3 | 0.045 | 0.084 | 6.4 | 0.014 | 0.083 | 3.3 | 0.016 | 0.160 | 1.3 | 0.258 | 0.018 |
| Dorsal Attention A: Superior Parietal Lobule 1 | 19.1 | 0.000 | 0.210 | 10.5 | 0.000 | 0.228 | 19.2 | 0.000 | 0.213 | 6.6 | 0.000 | 0.276 | 7.9 | 0.006 | 0.103 |
| Dorsal Attention B: Post Central 1 | 8.5 | 0.007 | 0.105 | 4.2 | 0.019 | 0.106 | 6.1 | 0.016 | 0.079 | 2.3 | 0.065 | 0.119 | 2.5 | 0.122 | 0.034 |
| Dorsal Attention B: Post Central 2 | 7.5 | 0.010 | 0.095 | 4.6 | 0.013 | 0.115 | 9.1 | 0.004 | 0.113 | 2.8 | 0.030 | 0.142 | 4.7 | 0.033 | 0.064 |
| Dorsal Attention B: Post Central 3 | 13.0 | 0.001 | 0.153 | 7.9 | 0.001 | 0.182 | 15.8 | 0.000 | 0.182 | 4.5 | 0.003 | 0.207 | 6.8 | 0.011 | 0.089 |
| Dorsal Attention B: Frontal Eye Fields 1 | 14.3 | 0.001 | 0.166 | 8.2 | 0.001 | 0.188 | 4.7 | 0.034 | 0.062 | 4.4 | 0.003 | 0.203 | 2.4 | 0.130 | 0.033 |
| Salience Ventral Attention A: Parietal Operculum 1 | 15.3 | 0.001 | 0.175 | 7.7 | 0.001 | 0.178 | 8.6 | 0.005 | 0.108 | 4.8 | 0.002 | 0.217 | 4.4 | 0.040 | 0.060 |
| Salience Ventral Attention A: Insula: 1 | 16.5 | 0.000 | 0.186 | 9.2 | 0.000 | 0.206 | 8.0 | 0.006 | 0.102 | 4.8 | 0.002 | 0.217 | 4.1 | 0.047 | 0.056 |
| Salience Ventral Attention A: Insula: 2 | 18.1 | 0.000 | 0.201 | 9.2 | 0.000 | 0.205 | 13.4 | 0.000 | 0.159 | 5.2 | 0.001 | 0.233 | 6.4 | 0.014 | 0.085 |
| Salience Ventral Attention A: Parietal Medial 1 | 14.2 | 0.001 | 0.165 | 8.0 | 0.001 | 0.184 | 14.3 | 0.000 | 0.168 | 4.8 | 0.002 | 0.219 | 6.5 | 0.013 | 0.086 |
| Salience Ventral Attention A: Frontal Medial 1 | 10.3 | 0.004 | 0.125 | 6.2 | 0.003 | 0.148 | 1.6 | 0.212 | 0.022 | 3.1 | 0.020 | 0.154 | 1.0 | 0.315 | 0.015 |
| Salience Ventral Attention B: Lateral Prefrontal Cortex 1 | 14.9 | 0.001 | 0.172 | 7.4 | 0.001 | 0.172 | 10.2 | 0.002 | 0.126 | 4.8 | 0.002 | 0.218 | 3.6 | 0.062 | 0.050 |
| Salience Ventral Attention B: Medial Posterior Prefrontal 1 | 19.0 | 0.000 | 0.209 | 10.6 | 0.000 | 0.230 | 17.4 | 0.000 | 0.196 | 5.3 | 0.001 | 0.233 | 10.5 | 0.002 | 0.132 |
| LimbiC: B: Orbital Frontal Cortex 1 | 21.4 | 0.000 | 0.229 | 13.9 | 0.000 | 0.281 | 27.1 | 0.000 | 0.276 | 8.7 | 0.000 | 0.336 | 9.3 | 0.003 | 0.119 |
| LimbiC: A: Temporal Pole 1 | 30.0 | 0.000 | 0.294 | 17.8 | 0.000 | 0.334 | 26.6 | 0.000 | 0.273 | 11.5 | 0.000 | 0.399 | 9.1 | 0.004 | 0.117 |
| LimbiC: A: Temporal Pole 2 | 28.2 | 0.000 | 0.282 | 14.2 | 0.000 | 0.285 | 26.1 | 0.000 | 0.269 | 8.8 | 0.000 | 0.337 | 13.6 | 0.000 | 0.165 |
| Control A: Intraparietal Sulcus 1 | 16.6 | 0.000 | 0.188 | 8.4 | 0.001 | 0.191 | 13.1 | 0.001 | 0.155 | 5.2 | 0.001 | 0.232 | 5.8 | 0.019 | 0.077 |
| Control A: Lateral Prefrontal Cortex 1 | 15.2 | 0.001 | 0.175 | 7.8 | 0.001 | 0.180 | 12.3 | 0.001 | 0.148 | 5.0 | 0.001 | 0.223 | 4.7 | 0.034 | 0.063 |
| Control A: Lateral Prefrontal Cortex 2 | 16.4 | 0.000 | 0.186 | 8.2 | 0.001 | 0.188 | 10.4 | 0.002 | 0.128 | 5.5 | 0.001 | 0.240 | 4.5 | 0.037 | 0.061 |
| Control B: Lateral Prefrontal Cortex 1 | 24.2 | 0.000 | 0.252 | 12.1 | 0.000 | 0.254 | 22.9 | 0.000 | 0.244 | 9.4 | 0.000 | 0.353 | 6.0 | 0.017 | 0.080 |
| Control C: Precuneus 1 | 10.2 | 0.004 | 0.124 | 5.9 | 0.004 | 0.143 | 11.6 | 0.001 | 0.140 | 3.6 | 0.010 | 0.173 | 4.7 | 0.033 | 0.064 |
| Control C: Precuneus 2 | 7.3 | 0.011 | 0.092 | 4.1 | 0.022 | 0.103 | 7.9 | 0.006 | 0.100 | 2.2 | 0.074 | 0.115 | 3.6 | 0.062 | 0.050 |
| Control C: Cingulate Posterior 1 | 11.7 | 0.002 | 0.139 | 6.3 | 0.003 | 0.152 | 10.8 | 0.002 | 0.132 | 4.0 | 0.005 | 0.190 | 3.8 | 0.054 | 0.053 |
| Default A: Dorsal Prefrontal Cortex 1 | 18.3 | 0.000 | 0.203 | 9.0 | 0.000 | 0.203 | 12.3 | 0.001 | 0.148 | 5.2 | 0.001 | 0.231 | 5.4 | 0.023 | 0.073 |
| Default A: Precuneus Posterior Cingulate Cortex1 | 7.0 | 0.012 | 0.088 | 3.4 | 0.037 | 0.088 | 4.6 | 0.036 | 0.060 | 2.3 | 0.066 | 0.118 | 1.2 | 0.269 | 0.018 |
| Default A: Medial Prefrontal Cortex 1 | 22.6 | 0.000 | 0.239 | 11.5 | 0.000 | 0.245 | 19.7 | 0.000 | 0.217 | 6.2 | 0.000 | 0.263 | 8.4 | 0.005 | 0.108 |
| Default B: Temp 1 | 18.1 | 0.000 | 0.201 | 10.1 | 0.000 | 0.221 | 6.1 | 0.016 | 0.079 | 6.0 | 0.000 | 0.259 | 1.9 | 0.172 | 0.027 |
| Default B: Temp 2 | 8.5 | 0.007 | 0.106 | 5.0 | 0.009 | 0.124 | 9.5 | 0.003 | 0.118 | 3.2 | 0.019 | 0.155 | 3.6 | 0.061 | 0.050 |
| Default B: Inferior Parietal Lobule 1 | 13.7 | 0.001 | 0.160 | 6.8 | 0.002 | 0.161 | 5.6 | 0.020 | 0.074 | 5.1 | 0.001 | 0.228 | 1.9 | 0.172 | 0.027 |
| Default B: Dorsal Prefrontal Cortex 1 | 19.3 | 0.000 | 0.212 | 9.9 | 0.000 | 0.219 | 5.5 | 0.022 | 0.072 | 5.4 | 0.001 | 0.240 | 2.6 | 0.110 | 0.037 |
| Default B: Lateral Prefrontal Cortex 1 | 10.4 | 0.003 | 0.126 | 5.3 | 0.007 | 0.129 | 8.4 | 0.005 | 0.106 | 3.5 | 0.011 | 0.169 | 3.0 | 0.086 | 0.042 |
| Default B: Ventral Prefrontal Cortex 1 | 19.8 | 0.000 | 0.216 | 11.1 | 0.000 | 0.239 | 21.4 | 0.000 | 0.232 | 5.6 | 0.001 | 0.246 | 11.1 | 0.001 | 0.138 |
| Default B: Ventral Prefrontal Cortex 2 | 34.9 | 0.000 | 0.327 | 17.2 | 0.000 | 0.327 | 21.6 | 0.000 | 0.234 | 10.0 | 0.000 | 0.367 | 9.3 | 0.003 | 0.118 |
| Default C: Retro Superior Parietal Lobuleenial 1 | 10.1 | 0.004 | 0.123 | 5.5 | 0.006 | 0.134 | 6.4 | 0.014 | 0.083 | 4.3 | 0.003 | 0.201 | 1.1 | 0.295 | 0.016 |
| Default C: Parahippocampal Cortex 1 | 13.3 | 0.001 | 0.156 | 7.0 | 0.002 | 0.165 | 10.3 | 0.002 | 0.126 | 3.6 | 0.010 | 0.174 | 5.2 | 0.026 | 0.070 |
| Temporal Parietal 1 | 4.9 | 0.034 | 0.064 | 2.5 | 0.087 | 0.066 | 4.3 | 0.043 | 0.057 | 1.8 | 0.147 | 0.093 | 1.5 | 0.224 | 0.021 |

| | Model 1: Age Category | | | Model 2: Age Category and Cortical Thickness | | | | | | Model 3: Age Category, Cortical Thickness, Systolic and Diastolic Blood Pressure | | | | | |
|---|---|---|---|---|---|---|---|---|---|---|---|---|---|---|---|
| | Age Category | | | Overall Model | | | Age Category | | | Overall Model | | | Age Category | | |
| Visual Central: Extra Striate Cortex 1 | 3.1 | 0.084 | 0.041 | 1.5 | 0.223 | 0.041 | 2.3 | 0.134 | 0.031 | 0.9 | 0.465 | 0.050 | 0.7 | 0.422 | 0.009 |
| Visual Central: Extra Striate Cortex 2 | 14.4 | 0.001 | 0.166 | 7.2 | 0.001 | 0.168 | 13.0 | 0.001 | 0.155 | 4.6 | 0.002 | 0.211 | 4.8 | 0.032 | 0.065 |
| Visual Central: Extra Striate Cortex 3 | 7.2 | 0.011 | 0.091 | 3.7 | 0.028 | 0.095 | 6.8 | 0.011 | 0.088 | 3.5 | 0.012 | 0.168 | 1.5 | 0.220 | 0.022 |
| Visual Peripheral: Striate Cortex Calcarine 1 | 2.7 | 0.107 | 0.036 | 1.4 | 0.244 | 0.039 | 2.4 | 0.128 | 0.032 | 1.1 | 0.345 | 0.062 | 0.3 | 0.578 | 0.004 |
| Visual Peripheral: Extra Striate Inferior 1 | 1.5 | 0.226 | 0.020 | 0.8 | 0.473 | 0.021 | 0.8 | 0.367 | 0.011 | 0.8 | 0.506 | 0.046 | 0.0 | 0.918 | 0.000 |
| Visual Peripheral: Extra Striate Superior 1 | 4.1 | 0.050 | 0.054 | 2.2 | 0.114 | 0.059 | 4.4 | 0.040 | 0.058 | 1.6 | 0.176 | 0.086 | 1.3 | 0.261 | 0.018 |
| Somatomotor A: 1 | 4.2 | 0.048 | 0.055 | 3.7 | 0.029 | 0.095 | 1.2 | 0.272 | 0.017 | 2.2 | 0.075 | 0.114 | 0.1 | 0.700 | 0.002 |
| Somatomotor A: 2 | 6.4 | 0.016 | 0.082 | 3.9 | 0.026 | 0.098 | 4.1 | 0.047 | 0.055 | 2.3 | 0.072 | 0.116 | 1.6 | 0.213 | 0.022 |
| Somatomotor A: 3 | 10.7 | 0.003 | 0.129 | 5.4 | 0.007 | 0.132 | 7.4 | 0.008 | 0.095 | 3.2 | 0.017 | 0.158 | 2.4 | 0.126 | 0.034 |
| Somatomotor A: 4 | 8.1 | 0.008 | 0.101 | 4.7 | 0.012 | 0.117 | 2.8 | 0.098 | 0.038 | 2.6 | 0.042 | 0.132 | 1.4 | 0.246 | 0.019 |
| Somatomotor B: Auditory 1 | 7.5 | 0.010 | 0.094 | 3.9 | 0.024 | 0.100 | 2.5 | 0.118 | 0.034 | 2.9 | 0.029 | 0.143 | 0.2 | 0.642 | 0.003 |
| Somatomotor B: S2 1 | 9.1 | 0.005 | 0.113 | 4.5 | 0.014 | 0.113 | 7.3 | 0.009 | 0.093 | 3.0 | 0.024 | 0.149 | 1.4 | 0.247 | 0.019 |
| Somatomotor B: S2 2 | 11.5 | 0.002 | 0.138 | 7.1 | 0.002 | 0.167 | 3.9 | 0.052 | 0.052 | 4.7 | 0.002 | 0.214 | 0.3 | 0.560 | 0.005 |
| Somatomotor B: Central 1 | 4.0 | 0.052 | 0.053 | 2.3 | 0.105 | 0.061 | 1.7 | 0.200 | 0.023 | 1.9 | 0.121 | 0.099 | 0.1 | 0.715 | 0.002 |
| Dorsal Attention A: Temporal Occipital 1 | 12.8 | 0.001 | 0.151 | 6.3 | 0.003 | 0.151 | 10.8 | 0.002 | 0.132 | 4.7 | 0.002 | 0.215 | 4.1 | 0.047 | 0.056 |
| Dorsal Attention A: Parietal Occipital 1 | 7.6 | 0.010 | 0.096 | 3.8 | 0.028 | 0.096 | 5.2 | 0.026 | 0.068 | 2.8 | 0.032 | 0.140 | 1.5 | 0.218 | 0.022 |
| Dorsal Attention A: Superior Parietal Lobule 1 | 10.1 | 0.004 | 0.123 | 5.7 | 0.005 | 0.138 | 11.1 | 0.001 | 0.135 | 3.7 | 0.009 | 0.176 | 4.3 | 0.042 | 0.059 |
| Dorsal Attention B: Post Central 1 | 6.0 | 0.020 | 0.077 | 3.4 | 0.040 | 0.087 | 5.6 | 0.020 | 0.074 | 2.9 | 0.028 | 0.144 | 1.9 | 0.169 | 0.027 |
| Dorsal Attention B: Post Central 2 | 9.7 | 0.004 | 0.119 | 5.4 | 0.007 | 0.131 | 10.4 | 0.002 | 0.128 | 3.2 | 0.019 | 0.155 | 4.1 | 0.047 | 0.056 |
| Dorsal Attention B: Frontal Eye Fields 1 | 16.4 | 0.000 | 0.186 | 8.5 | 0.001 | 0.192 | 8.7 | 0.004 | 0.109 | 4.3 | 0.004 | 0.200 | 5.3 | 0.024 | 0.072 |
| Salience Ventral Attention A: Parietal Operculum 1 | 11.4 | 0.002 | 0.137 | 6.2 | 0.003 | 0.149 | 6.0 | 0.017 | 0.078 | 4.2 | 0.004 | 0.197 | 1.1 | 0.296 | 0.016 |
| Salience Ventral Attention A: Insula: 1 | 14.3 | 0.001 | 0.165 | 7.2 | 0.001 | 0.169 | 6.3 | 0.014 | 0.081 | 4.4 | 0.003 | 0.202 | 1.7 | 0.195 | 0.024 |
| Salience Ventral Attention A: Parietal Medial 1 | 8.8 | 0.006 | 0.109 | 5.3 | 0.007 | 0.130 | 10.2 | 0.002 | 0.126 | 3.3 | 0.015 | 0.162 | 3.8 | 0.056 | 0.052 |
| Salience Ventral Attention A: Frontal Medial 1 | 14.3 | 0.001 | 0.165 | 7.1 | 0.002 | 0.166 | 6.5 | 0.013 | 0.084 | 4.1 | 0.005 | 0.193 | 2.5 | 0.118 | 0.035 |
| Salience Ventral Attention B: Inferior Parietal Lobule 1 | 13.8 | 0.001 | 0.161 | 7.9 | 0.001 | 0.182 | 7.1 | 0.010 | 0.091 | 5.3 | 0.001 | 0.236 | 1.9 | 0.178 | 0.026 |
| Salience Ventral Attention B: Lateral Prefrontal Cortex 1 | 12.9 | 0.001 | 0.152 | 8.2 | 0.001 | 0.187 | 4.7 | 0.033 | 0.062 | 5.7 | 0.000 | 0.250 | 0.6 | 0.446 | 0.008 |
| Salience Ventral Attention B: Medial Posterior Prefrontal 1 | 15.6 | 0.001 | 0.178 | 10.2 | 0.000 | 0.223 | 19.8 | 0.000 | 0.218 | 5.3 | 0.001 | 0.236 | 8.9 | 0.004 | 0.114 |
| LimbiC: B: Orbital Frontal Cortex 1 | 20.0 | 0.000 | 0.217 | 10.9 | 0.000 | 0.236 | 20.7 | 0.000 | 0.226 | 6.9 | 0.000 | 0.284 | 5.7 | 0.019 | 0.077 |
| LimbiC: A: Temporal Pole 1 | 33.4 | 0.000 | 0.317 | 17.1 | 0.000 | 0.325 | 29.1 | 0.000 | 0.291 | 10.8 | 0.000 | 0.386 | 11.5 | 0.001 | 0.143 |
| Control A: Intraparietal Sulcus 1 | 10.5 | 0.003 | 0.127 | 5.2 | 0.008 | 0.127 | 7.1 | 0.010 | 0.090 | 4.4 | 0.003 | 0.202 | 1.5 | 0.232 | 0.021 |
| Control A: Lateral Prefrontal Cortex 1 | 16.2 | 0.000 | 0.183 | 8.3 | 0.001 | 0.189 | 8.7 | 0.004 | 0.109 | 5.7 | 0.001 | 0.248 | 2.9 | 0.094 | 0.040 |
| Control A: Lateral Prefrontal Cortex 2 | 20.8 | 0.000 | 0.224 | 11.9 | 0.000 | 0.252 | 7.3 | 0.009 | 0.093 | 6.6 | 0.000 | 0.276 | 3.4 | 0.070 | 0.047 |
| Control B: Temporal 1 | 23.4 | 0.000 | 0.246 | 12.1 | 0.000 | 0.254 | 15.7 | 0.000 | 0.181 | 7.9 | 0.000 | 0.314 | 4.5 | 0.037 | 0.061 |
| Control B: inferior parietal lobule 1 | 9.1 | 0.005 | 0.112 | 4.5 | 0.015 | 0.112 | 6.5 | 0.013 | 0.084 | 3.3 | 0.015 | 0.162 | 2.1 | 0.157 | 0.029 |
| Control B: Lateral Prefrontal Cortexd 1 | 15.6 | 0.001 | 0.178 | 8.1 | 0.001 | 0.185 | 6.6 | 0.012 | 0.085 | 4.6 | 0.002 | 0.212 | 2.2 | 0.139 | 0.031 |
| Control B: Lateral Prefrontal Cortexv 1 | 18.4 | 0.000 | 0.203 | 9.1 | 0.000 | 0.203 | 15.7 | 0.000 | 0.182 | 8.4 | 0.000 | 0.326 | 2.0 | 0.159 | 0.029 |
| Control C: Cingulate Posterior 1 | 9.9 | 0.004 | 0.121 | 4.9 | 0.010 | 0.122 | 6.4 | 0.013 | 0.083 | 3.1 | 0.022 | 0.150 | 1.9 | 0.169 | 0.027 |
| Control C: Precuneus 1 | 5.9 | 0.021 | 0.076 | 3.0 | 0.057 | 0.078 | 4.2 | 0.045 | 0.055 | 2.1 | 0.096 | 0.107 | 1.3 | 0.261 | 0.018 |
| Default A: Inferior Parietal Lobule 1 | 9.2 | 0.005 | 0.114 | 4.8 | 0.012 | 0.118 | 3.3 | 0.072 | 0.045 | 3.1 | 0.022 | 0.151 | 1.1 | 0.294 | 0.016 |
| Default A: Dorsal Prefrontal Cortex 1 | 20.3 | 0.000 | 0.220 | 10.6 | 0.000 | 0.230 | 10.0 | 0.002 | 0.123 | 5.6 | 0.001 | 0.245 | 5.4 | 0.023 | 0.073 |
| Default A: Precuneus Posterior Cingulate Cortex 1 | 3.8 | 0.056 | 0.051 | 2.1 | 0.136 | 0.055 | 2.2 | 0.142 | 0.030 | 1.6 | 0.194 | 0.083 | 0.4 | 0.543 | 0.005 |
| Default A: Medial Prefrontal Cortex 1 | 17.8 | 0.000 | 0.199 | 9.4 | 0.000 | 0.209 | 10.3 | 0.002 | 0.127 | 5.8 | 0.000 | 0.250 | 2.5 | 0.115 | 0.036 |
| Default B: Dorsal Prefrontal Cortex 1 | 24.2 | 0.000 | 0.251 | 12.6 | 0.000 | 0.262 | 8.8 | 0.004 | 0.110 | 7.0 | 0.000 | 0.288 | 4.2 | 0.045 | 0.057 |
| Default B: Ventral Prefrontal Cortex 1 | 21.4 | 0.000 | 0.229 | 10.6 | 0.000 | 0.229 | 16.1 | 0.000 | 0.185 | 6.7 | 0.000 | 0.278 | 5.7 | 0.020 | 0.077 |
| Default B: Ventral Prefrontal Cortex 2 | 19.9 | 0.000 | 0.216 | 10.7 | 0.000 | 0.232 | 8.6 | 0.004 | 0.108 | 6.5 | 0.000 | 0.273 | 2.6 | 0.111 | 0.036 |
| Default C: Retro Superior Parietal Lobuleenial 1 | 5.7 | 0.022 | 0.073 | 2.9 | 0.061 | 0.076 | 3.9 | 0.052 | 0.052 | 2.5 | 0.052 | 0.126 | 0.5 | 0.469 | 0.008 |
| Default C: Parahippocampal Cortex 1 | 15.7 | 0.001 | 0.179 | 8.1 | 0.001 | 0.185 | 11.8 | 0.001 | 0.143 | 4.1 | 0.005 | 0.191 | 5.5 | 0.022 | 0.074 |
| Temporal Parietal 1 | 22.1 | 0.000 | 0.235 | 10.9 | 0.000 | 0.236 | 16.6 | 0.000 | 0.190 | 5.9 | 0.000 | 0.255 | 7.4 | 0.008 | 0.097 |
| Temporal Parietal 2 | 12.5 | 0.001 | 0.148 | 6.3 | 0.003 | 0.150 | 8.1 | 0.006 | 0.102 | 4.2 | 0.004 | 0.195 | 1.8 | 0.183 | 0.026 |
| Temporal Parietal 3 | 5.7 | 0.023 | 0.074 | 3.0 | 0.058 | 0.077 | 4.8 | 0.032 | 0.063 | 2.4 | 0.059 | 0.121 | 1.3 | 0.253 | 0.019 |

Table S2. General linear models of the association between regional cerebral blood flow among the four groups based on age category and insulin resistance levels, with blood pressure and cortical thickness as covariates. Post-hoc contrasts in the 40 regions with significant group difference are shown, comparing the young insulin sensitive group to the other three groups. The group effect sizes are plotted on the brain surface in Figure 2 in main document.

| | Overall | | | 4 Groups Based on Age Group and HOMA-IR Median Split | | | | | | Systolic BP | | | Diastolic BP | | | Cortical Thickness | | |
|---|---|---|---|---|---|---|---|---|---|---|---|---|---|---|---|---|---|---|
| | F | p-FDR | $\eta^2_p$ | F | p | $\eta^2_p$ | YIS vs YIR | YIS vs OIS | YIS vs OIR | F | p | $\eta^2_p$ | F | p | $\eta^2_p$ | F | p | $\eta^2_p$ |
| Visual Central: Extra Striate Cortex 1 | 2.2 | 0.060 | 0.165 | 1.4 | 0.264 | 0.057 | | | | 0.4 | 0.543 | 0.006 | 1.4 | 0.237 | 0.021 | 0.0 | 0.916 | 0.000 |
| Visual Central: Extra Striate Cortex 2 | 6.1 | 0.000 | 0.352 | 3.2 | 0.028 | 0.126 | 0.361 | 0.781 | 0.040 | 2.4 | 0.126 | 0.035 | 2.6 | 0.113 | 0.037 | 0.2 | 0.658 | 0.003 |
| Visual Central: Striate Cortex 1 | 3.3 | 0.011 | 0.230 | 1.4 | 0.259 | 0.058 | | | | 2.1 | 0.151 | 0.031 | 0.8 | 0.385 | 0.011 | 2.0 | 0.161 | 0.029 |
| Visual Central: Extra Striate Cortex 3 | 3.9 | 0.005 | 0.256 | 2.6 | 0.061 | 0.104 | | | | 2.4 | 0.126 | 0.035 | 0.2 | 0.660 | 0.003 | 0.0 | 0.828 | 0.001 |
| Visual Peripheral: Extra Striate Inferior 1 | 1.1 | 0.363 | 0.092 | 0.9 | 0.459 | 0.038 | | | | 0.5 | 0.477 | 0.008 | 0.0 | 0.824 | 0.001 | 0.2 | 0.646 | 0.003 |
| Visual Peripheral: Striate Cortex Calcarine 1 | 1.3 | 0.267 | 0.106 | 0.9 | 0.429 | 0.040 | | | | 1.2 | 0.284 | 0.017 | 0.1 | 0.747 | 0.002 | 0.1 | 0.725 | 0.002 |
| Visual Peripheral: Extra Striate Cortex Sup 1 | 2.5 | 0.038 | 0.183 | 2.0 | 0.128 | 0.081 | | | | 1.5 | 0.224 | 0.022 | 0.0 | 0.951 | 0.000 | 0.4 | 0.536 | 0.006 |
| Somatomotor A: 1 | 2.5 | 0.038 | 0.184 | 2.7 | 0.055 | 0.106 | | | | 0.1 | 0.789 | 0.001 | 0.8 | 0.384 | 0.011 | 0.7 | 0.397 | 0.011 |
| Somatomotor A: 2 | 2.4 | 0.044 | 0.177 | 2.4 | 0.073 | 0.098 | | | | 0.0 | 0.879 | 0.000 | 0.2 | 0.658 | 0.003 | 0.2 | 0.664 | 0.003 |
| Somatomotor B: Auditory 1 | 1.8 | 0.123 | 0.137 | 1.6 | 0.197 | 0.067 | | | | 1.3 | 0.264 | 0.019 | 0.0 | 0.911 | 0.000 | 0.0 | 0.915 | 0.000 |
| Somatomotor B: S2 1 | 3.4 | 0.010 | 0.235 | 2.6 | 0.061 | 0.104 | | | | 2.2 | 0.143 | 0.032 | 0.1 | 0.769 | 0.001 | 0.0 | 0.927 | 0.000 |
| Somatomotor B: S2 2 | 2.2 | 0.058 | 0.166 | 1.7 | 0.178 | 0.070 | | | | 1.2 | 0.282 | 0.017 | 0.0 | 0.939 | 0.000 | 0.5 | 0.489 | 0.007 |
| Somatomotor B: Central 1 | 2.5 | 0.040 | 0.181 | 2.5 | 0.065 | 0.102 | | | | 1.1 | 0.297 | 0.016 | 0.3 | 0.608 | 0.004 | 0.1 | 0.715 | 0.002 |
| Dorsal Attention A: Temporal Occipital 1 | 3.8 | 0.006 | 0.255 | 2.1 | 0.104 | 0.087 | | | | 0.6 | 0.456 | 0.008 | 2.0 | 0.166 | 0.028 | 0.1 | 0.787 | 0.001 |
| Dorsal Attention A: Parietal Occipital 1 | 3.2 | 0.013 | 0.222 | 2.3 | 0.089 | 0.092 | | | | 2.6 | 0.109 | 0.038 | 0.4 | 0.538 | 0.006 | 1.1 | 0.292 | 0.017 |
| Dorsal Attention A: Superior Parietal Lobule 1 | 5.5 | 0.001 | 0.330 | 4.6 | 0.005 | 0.171 | 0.629 | 0.067 | 0.001 | 1.9 | 0.169 | 0.028 | 0.2 | 0.656 | 0.003 | 1.9 | 0.173 | 0.028 |
| Dorsal Attention B: Post Central 1 | 2.1 | 0.068 | 0.159 | 1.9 | 0.137 | 0.079 | | | | 0.8 | 0.383 | 0.011 | 0.0 | 0.827 | 0.001 | 0.0 | 0.925 | 0.000 |
| Dorsal Attention B: Post Central 2 | 2.7 | 0.030 | 0.193 | 3.0 | 0.035 | 0.120 | 0.614 | 0.169 | 0.006 | 0.4 | 0.547 | 0.005 | 0.6 | 0.455 | 0.008 | 2.2 | 0.143 | 0.032 |
| Dorsal Attention B: Post Central 3 | 3.7 | 0.006 | 0.251 | 3.6 | 0.017 | 0.140 | 0.388 | 0.050 | 0.002 | 1.5 | 0.227 | 0.022 | 0.0 | 0.850 | 0.001 | 3.7 | 0.059 | 0.052 |
| Dorsal Attention B: Frontal Eye Fields 1 | 3.3 | 0.012 | 0.227 | 1.5 | 0.227 | 0.062 | | | | 0.1 | 0.737 | 0.002 | 0.5 | 0.471 | 0.008 | 1.6 | 0.215 | 0.023 |
| Salience Ventral Attention A: Parietal Operculum 1 | 4.3 | 0.003 | 0.276 | 3.4 | 0.024 | 0.131 | 0.893 | 0.306 | 0.007 | 0.5 | 0.494 | 0.007 | 1.3 | 0.263 | 0.019 | 0.1 | 0.763 | 0.001 |
| Salience Ventral Attention A: Insula: 1 | 3.5 | 0.009 | 0.236 | 1.9 | 0.135 | 0.079 | | | | 0.2 | 0.646 | 0.003 | 0.1 | 0.704 | 0.002 | 1.8 | 0.183 | 0.026 |
| Salience Ventral Attention A: Insula: 2 | 3.9 | 0.005 | 0.259 | 2.9 | 0.041 | 0.115 | 0.790 | 0.089 | 0.008 | 2.0 | 0.158 | 0.029 | 0.2 | 0.658 | 0.003 | 0.6 | 0.444 | 0.009 |
| Salience Ventral Attention A: Parietal Medial 1 | 4.3 | 0.003 | 0.276 | 4.0 | 0.011 | 0.152 | 0.969 | 0.227 | 0.004 | 0.5 | 0.470 | 0.008 | 1.1 | 0.297 | 0.016 | 0.5 | 0.469 | 0.008 |
| Salience Ventral Attention A: Frontal Medial 1 | 2.6 | 0.035 | 0.187 | 1.3 | 0.295 | 0.053 | | | | 0.0 | 0.936 | 0.000 | 0.5 | 0.461 | 0.008 | 0.8 | 0.372 | 0.012 |
| Salience Ventral Attention B: Lateral Prefrontal Cortex 1 | 3.6 | 0.007 | 0.245 | 2.0 | 0.118 | 0.083 | | | | 2.9 | 0.091 | 0.042 | 0.1 | 0.796 | 0.001 | 0.1 | 0.713 | 0.002 |
| Salience Ventral Attention B: Medial Posterior Prefrontal 1 | 4.8 | 0.002 | 0.299 | 5.8 | 0.001 | 0.207 | 0.965 | 0.036 | 0.000 | 0.0 | 0.943 | 0.000 | 0.6 | 0.449 | 0.009 | 2.0 | 0.165 | 0.029 |
| Limbic: B: Orbital Frontal Cortex 1 | 6.6 | 0.000 | 0.372 | 4.5 | 0.006 | 0.167 | 0.392 | 0.029 | 0.001 | 3.0 | 0.088 | 0.043 | 0.0 | 0.936 | 0.000 | 4.4 | 0.040 | 0.062 |
| Limbic: A: Temporal Pole 1 | 8.6 | 0.000 | 0.435 | 4.6 | 0.006 | 0.169 | 0.434 | 0.031 | 0.001 | 2.2 | 0.140 | 0.032 | 0.6 | 0.444 | 0.009 | 5.1 | 0.028 | 0.070 |
| Limbic: A: Temporal Pole 2 | 6.2 | 0.000 | 0.356 | 5.2 | 0.003 | 0.189 | 0.776 | 0.029 | 0.002 | 0.2 | 0.673 | 0.003 | 2.6 | 0.115 | 0.037 | 0.2 | 0.641 | 0.003 |
| Control A: Intraparietal Sulcus 1 | 4.4 | 0.003 | 0.285 | 3.7 | 0.017 | 0.141 | 0.835 | 0.218 | 0.004 | 0.6 | 0.453 | 0.008 | 1.1 | 0.297 | 0.016 | 0.0 | 0.993 | 0.000 |
| Control A: Lateral Prefrontal Cortex 1 | 4.4 | 0.003 | 0.282 | 3.5 | 0.021 | 0.135 | 0.633 | 0.160 | 0.004 | 3.0 | 0.088 | 0.043 | 0.1 | 0.759 | 0.001 | 1.2 | 0.286 | 0.017 |
| Control A: Lateral Prefrontal Cortex 2 | 4.5 | 0.003 | 0.287 | 3.0 | 0.036 | 0.119 | 0.983 | 0.254 | 0.012 | 3.2 | 0.080 | 0.045 | 0.0 | 0.972 | 0.000 | 0.7 | 0.406 | 0.010 |
| Control B: Lateral Prefrontal Cortex 1 | 8.4 | 0.000 | 0.428 | 5.1 | 0.003 | 0.186 | 0.256 | 0.085 | 0.000 | 8.3 | 0.005 | 0.111 | 0.4 | 0.551 | 0.005 | 1.5 | 0.228 | 0.022 |
| Control C: Precuneus 1 | 3.2 | 0.012 | 0.225 | 3.1 | 0.031 | 0.123 | 0.929 | 0.316 | 0.010 | 0.8 | 0.373 | 0.012 | 0.4 | 0.533 | 0.006 | 1.1 | 0.308 | 0.016 |
| Control C: Precuneus 2 | 2.5 | 0.041 | 0.180 | 3.0 | 0.035 | 0.119 | 0.390 | 0.156 | 0.005 | 0.5 | 0.480 | 0.007 | 0.0 | 0.869 | 0.000 | 2.0 | 0.167 | 0.028 |
| Control C: Cingulate Posterior 1 | 4.9 | 0.002 | 0.303 | 5.1 | 0.003 | 0.185 | 0.912 | 0.369 | 0.002 | 1.3 | 0.257 | 0.019 | 0.6 | 0.460 | 0.008 | 1.3 | 0.264 | 0.019 |
| Default A: Dorsal Prefrontal Cortex 1 | 3.9 | 0.005 | 0.261 | 2.7 | 0.050 | 0.109 | 0.735 | 0.265 | 0.022 | 1.7 | 0.202 | 0.024 | 0.0 | 0.996 | 0.000 | 0.0 | 0.890 | 0.000 |
| Default A: Precuneus Posterior Cingulate Cortex1 | 2.7 | 0.029 | 0.194 | 2.5 | 0.064 | 0.102 | | | | 1.1 | 0.294 | 0.016 | 0.2 | 0.650 | 0.003 | 0.1 | 0.820 | 0.001 |
| Default A: Medial Prefrontal Cortex 1 | 5.9 | 0.001 | 0.346 | 5.9 | 0.001 | 0.209 | 0.635 | 0.081 | 0.000 | 1.1 | 0.289 | 0.017 | 0.0 | 0.939 | 0.000 | 0.3 | 0.618 | 0.004 |
| Default B: Temp 1 | 4.5 | 0.003 | 0.286 | 1.5 | 0.230 | 0.062 | | | | 1.4 | 0.246 | 0.020 | 0.1 | 0.731 | 0.002 | 2.0 | 0.164 | 0.029 |
| Default B: Temp 2 | 2.8 | 0.024 | 0.201 | 2.5 | 0.063 | 0.102 | | | | 1.1 | 0.291 | 0.017 | 0.1 | 0.802 | 0.001 | 1.5 | 0.232 | 0.021 |
| Default B: Inferior Parietal Lobule 1 | 4.9 | 0.002 | 0.306 | 3.2 | 0.029 | 0.125 | 0.895 | 0.493 | 0.013 | 2.2 | 0.143 | 0.032 | 0.7 | 0.408 | 0.010 | 0.1 | 0.811 | 0.001 |
| Default B: Dorsal Prefrontal Cortex 1 | 3.9 | 0.005 | 0.259 | 1.4 | 0.238 | 0.061 | | | | 1.6 | 0.205 | 0.024 | 0.1 | 0.706 | 0.002 | 0.2 | 0.622 | 0.004 |
| Default B: Lateral Prefrontal Cortex 1 | 2.8 | 0.024 | 0.201 | 1.9 | 0.138 | 0.078 | | | | 2.1 | 0.151 | 0.031 | 0.0 | 0.925 | 0.000 | 0.8 | 0.376 | 0.012 |
| Default B: Ventral Prefrontal Cortex 1 | 4.0 | 0.005 | 0.264 | 4.2 | 0.008 | 0.159 | 0.807 | 0.050 | 0.004 | 0.3 | 0.559 | 0.005 | 0.0 | 0.831 | 0.001 | 1.2 | 0.279 | 0.017 |
| Default B: Ventral Prefrontal Cortex 2 | 7.9 | 0.000 | 0.415 | 5.1 | 0.003 | 0.186 | 0.622 | 0.030 | 0.000 | 3.2 | 0.079 | 0.045 | 0.0 | 0.833 | 0.001 | 0.1 | 0.798 | 0.001 |
| Default C: Retro Superior Parietal Lobuleenial 1 | 3.4 | 0.010 | 0.232 | 1.3 | 0.284 | 0.055 | | | | 1.7 | 0.203 | 0.024 | 0.8 | 0.364 | 0.012 | 1.3 | 0.262 | 0.019 |
| Default C: Parahippocampal Cortex 1 | 2.5 | 0.037 | 0.184 | 2.0 | 0.124 | 0.082 | | | | 0.1 | 0.791 | 0.001 | 0.4 | 0.550 | 0.005 | 1.0 | 0.314 | 0.015 |
| Temporal Parietal 1 | 1.9 | 0.103 | 0.144 | 1.9 | 0.144 | 0.077 | | | | 1.2 | 0.282 | 0.017 | 0.1 | 0.791 | 0.001 | 0.9 | 0.354 | 0.013 |

| | Overall | | | 4 Groups Based on Age Group and HOMA-IR Median Split | | | | | | Systolic BP | | | Diastolic BP | | | Cortical Thickness | | |
|---|---|---|---|---|---|---|---|---|---|---|---|---|---|---|---|---|---|---|
| | F | p-FDR | $\eta^2_p$ | F | p | $\eta^2_p$ | YIS vs YIR | YIS vs OIS | YIS vs OIR | F | p | $\eta^2_p$ | F | p | $\eta^2_p$ | F | p | $\eta^2_p$ |
| Visual Central: Extra Striate Cortex 1 | 0.8 | 0.544 | 0.070 | 0.7 | 0.558 | 0.030 | | | | 0.3 | 0.566 | 0.005 | 0.0 | 0.845 | 0.001 | 0.0 | 0.959 | 0.000 |
| Visual Central: Extra Striate Cortex 2 | 4.5 | 0.003 | 0.289 | 4.2 | 0.009 | 0.158 | 0.458 | 0.663 | 0.013 | 0.7 | 0.422 | 0.010 | 1.7 | 0.198 | 0.025 | 0.2 | 0.679 | 0.003 |
| Visual Central: Extra Striate Cortex 3 | 3.9 | 0.005 | 0.260 | 3.3 | 0.024 | 0.130 | 0.873 | 0.974 | 0.027 | 3.1 | 0.081 | 0.045 | 0.2 | 0.642 | 0.003 | 0.7 | 0.391 | 0.011 |
| Visual Peripheral: Striate Cortex Calcarine 1 | 1.0 | 0.412 | 0.086 | 0.7 | 0.567 | 0.030 | | | | 0.8 | 0.380 | 0.012 | 0.1 | 0.769 | 0.001 | 0.1 | 0.768 | 0.001 |
| Visual Peripheral: Extra Striate Inferior 1 | 0.9 | 0.476 | 0.078 | 0.8 | 0.518 | 0.033 | | | | 0.9 | 0.350 | 0.013 | 0.1 | 0.752 | 0.002 | 0.1 | 0.819 | 0.001 |
| Visual Peripheral: Extra Striate Superior 1 | 2.1 | 0.067 | 0.160 | 2.4 | 0.074 | 0.098 | | | | 0.6 | 0.447 | 0.009 | 0.6 | 0.425 | 0.010 | 0.3 | 0.569 | 0.005 |
| Somatomotor A: 1 | 2.9 | 0.021 | 0.206 | 2.6 | 0.057 | 0.106 | | | | 0.3 | 0.598 | 0.004 | 0.8 | 0.386 | 0.011 | 4.5 | 0.037 | 0.064 |
| Somatomotor A: 2 | 2.2 | 0.057 | 0.167 | 1.9 | 0.134 | 0.079 | | | | 0.1 | 0.781 | 0.001 | 0.8 | 0.377 | 0.012 | 1.8 | 0.188 | 0.026 |
| Somatomotor A: 3 | 2.5 | 0.036 | 0.186 | 1.6 | 0.203 | 0.066 | | | | 0.9 | 0.358 | 0.013 | 0.1 | 0.767 | 0.001 | 0.2 | 0.690 | 0.002 |
| Somatomotor A: 4 | 2.2 | 0.062 | 0.164 | 1.3 | 0.280 | 0.055 | | | | 0.1 | 0.822 | 0.001 | 0.7 | 0.416 | 0.010 | 1.4 | 0.248 | 0.020 |
| Somatomotor B: Auditory 1 | 3.1 | 0.015 | 0.218 | 2.2 | 0.094 | 0.090 | | | | 2.5 | 0.118 | 0.036 | 0.0 | 0.935 | 0.000 | 0.7 | 0.415 | 0.010 |
| Somatomotor B: S2 1 | 2.6 | 0.032 | 0.190 | 1.6 | 0.196 | 0.067 | | | | 2.0 | 0.159 | 0.029 | 0.0 | 0.888 | 0.000 | 0.0 | 0.968 | 0.000 |
| Somatomotor B: S2 2 | 3.8 | 0.006 | 0.253 | 1.3 | 0.290 | 0.054 | | | | 3.3 | 0.074 | 0.047 | 0.3 | 0.608 | 0.004 | 3.3 | 0.074 | 0.047 |
| Somatomotor B: Central 1 | 2.3 | 0.056 | 0.168 | 1.9 | 0.136 | 0.079 | | | | 1.5 | 0.224 | 0.022 | 0.0 | 0.864 | 0.000 | 0.5 | 0.486 | 0.007 |
| Dorsal Attention A: Temporal Occipital 1 | 3.6 | 0.007 | 0.246 | 2.3 | 0.085 | 0.094 | | | | 0.9 | 0.344 | 0.013 | 1.4 | 0.247 | 0.020 | 0.1 | 0.753 | 0.001 |
| Dorsal Attention A: Parietal Occipital 1 | 2.9 | 0.020 | 0.207 | 2.4 | 0.072 | 0.098 | | | | 1.0 | 0.321 | 0.015 | 0.8 | 0.372 | 0.012 | 0.0 | 0.895 | 0.000 |
| Dorsal Attention A: Superior Parietal Lobule 1 | 4.2 | 0.004 | 0.274 | 4.6 | 0.006 | 0.170 | 0.630 | 0.316 | 0.002 | 0.6 | 0.449 | 0.009 | 1.0 | 0.314 | 0.015 | 0.8 | 0.360 | 0.013 |
| Dorsal Attention B: Post Central 1 | 2.9 | 0.020 | 0.208 | 2.5 | 0.069 | 0.100 | | | | 2.3 | 0.136 | 0.033 | 0.2 | 0.668 | 0.003 | 1.3 | 0.256 | 0.019 |
| Dorsal Attention B: Post Central 2 | 2.9 | 0.022 | 0.205 | 2.8 | 0.046 | 0.112 | 0.829 | 0.236 | 0.011 | 0.3 | 0.564 | 0.005 | 0.6 | 0.446 | 0.009 | 1.1 | 0.308 | 0.016 |
| Dorsal Attention B: Frontal Eye Fields 1 | 3.4 | 0.010 | 0.232 | 2.7 | 0.052 | 0.108 | | | | 0.1 | 0.806 | 0.001 | 0.4 | 0.550 | 0.005 | 0.3 | 0.601 | 0.004 |
| Salience Ventral Attention A: Parietal Operculum 1 | 3.8 | 0.005 | 0.256 | 2.2 | 0.101 | 0.088 | | | | 2.1 | 0.156 | 0.030 | 0.0 | 0.838 | 0.001 | 1.1 | 0.300 | 0.016 |
| Salience Ventral Attention A: Insula: 1 | 3.8 | 0.006 | 0.252 | 2.1 | 0.109 | 0.086 | | | | 1.7 | 0.197 | 0.025 | 0.0 | 0.832 | 0.001 | 0.5 | 0.479 | 0.008 |
| Salience Ventral Attention A: Parietal Medial 1 | 3.3 | 0.011 | 0.229 | 3.3 | 0.026 | 0.128 | 0.612 | 0.545 | 0.021 | 0.8 | 0.369 | 0.012 | 0.7 | 0.408 | 0.010 | 1.4 | 0.249 | 0.020 |
| Salience Ventral Attention A: Frontal Medial 1 | 3.5 | 0.009 | 0.238 | 2.2 | 0.097 | 0.089 | | | | 0.6 | 0.460 | 0.008 | 0.6 | 0.439 | 0.009 | 0.0 | 0.942 | 0.000 |
| Salience Ventral Attention B: Inferior Parietal Lobule 1 | 4.2 | 0.004 | 0.273 | 1.8 | 0.159 | 0.074 | | | | 1.4 | 0.236 | 0.021 | 0.4 | 0.510 | 0.007 | 1.2 | 0.274 | 0.018 |
| Salience Ventral Attention B: Lateral Prefrontal Cortex 1 | 4.3 | 0.003 | 0.279 | 1.1 | 0.354 | 0.047 | | | | 2.8 | 0.097 | 0.041 | 0.1 | 0.787 | 0.001 | 2.0 | 0.162 | 0.029 |
| Salience Ventral Attention B: Medial Posterior Prefrontal 1 | 4.5 | 0.003 | 0.288 | 4.7 | 0.005 | 0.174 | 0.903 | 0.088 | 0.002 | 0.6 | 0.433 | 0.009 | 0.1 | 0.716 | 0.002 | 3.1 | 0.081 | 0.045 |
| Limbic: B: Orbital Frontal Cortex 1 | 5.5 | 0.001 | 0.329 | 3.5 | 0.021 | 0.134 | 0.306 | 0.074 | 0.002 | 3.6 | 0.063 | 0.051 | 0.2 | 0.636 | 0.003 | 0.8 | 0.378 | 0.012 |
| Limbic: A: Temporal Pole 1 | 8.6 | 0.000 | 0.435 | 6.0 | 0.001 | 0.212 | 0.134 | 0.005 | 0.000 | 1.8 | 0.188 | 0.026 | 0.5 | 0.474 | 0.008 | 1.3 | 0.260 | 0.019 |
| Control A: Intraparietal Sulcus 1 | 4.1 | 0.004 | 0.270 | 2.6 | 0.059 | 0.105 | | | | 1.8 | 0.185 | 0.026 | 1.2 | 0.275 | 0.018 | 0.0 | 0.903 | 0.000 |
| Control A: Lateral Prefrontal Cortex 1 | 4.8 | 0.002 | 0.298 | 2.6 | 0.058 | 0.105 | | | | 3.2 | 0.076 | 0.046 | 0.0 | 0.991 | 0.000 | 0.0 | 0.939 | 0.000 |
| Control A: Lateral Prefrontal Cortex 2 | 5.3 | 0.001 | 0.320 | 2.6 | 0.059 | 0.104 | | | | 0.5 | 0.463 | 0.008 | 0.5 | 0.501 | 0.007 | 2.1 | 0.149 | 0.031 |
| Control B: Temporal 1 | 6.1 | 0.001 | 0.353 | 2.9 | 0.043 | 0.114 | 0.404 | 0.106 | 0.005 | 3.1 | 0.083 | 0.044 | 0.0 | 0.901 | 0.000 | 1.0 | 0.325 | 0.014 |
| Control B: inferior parietal lobule 1 | 3.3 | 0.011 | 0.230 | 2.7 | 0.053 | 0.108 | | | | 0.8 | 0.378 | 0.012 | 1.0 | 0.320 | 0.015 | 0.0 | 0.827 | 0.001 |
| Control B: Lateral Prefrontal Cortexd 1 | 3.7 | 0.006 | 0.251 | 1.9 | 0.130 | 0.080 | | | | 0.9 | 0.334 | 0.014 | 0.2 | 0.641 | 0.003 | 0.6 | 0.446 | 0.009 |
| Control B: Lateral Prefrontal Cortexv 1 | 7.0 | 0.000 | 0.384 | 2.8 | 0.045 | 0.112 | 0.316 | 0.281 | 0.008 | 9.0 | 0.004 | 0.119 | 0.3 | 0.613 | 0.004 | 0.0 | 0.884 | 0.000 |
| Control C: Cingulate Posterior 1 | 4.1 | 0.004 | 0.270 | 4.4 | 0.007 | 0.165 | 0.796 | 0.991 | 0.013 | 0.7 | 0.414 | 0.010 | 1.0 | 0.333 | 0.014 | 0.3 | 0.595 | 0.004 |
| Control C: Precuneus 1 | 2.7 | 0.028 | 0.195 | 2.9 | 0.040 | 0.116 | 0.909 | 0.916 | 0.031 | 0.2 | 0.634 | 0.003 | 1.4 | 0.247 | 0.020 | 0.0 | 0.911 | 0.000 |
| Default A: Inferior Parietal Lobule 1 | 3.1 | 0.016 | 0.215 | 2.2 | 0.093 | 0.091 | | | | 1.2 | 0.286 | 0.017 | 0.3 | 0.608 | 0.004 | 0.0 | 0.901 | 0.000 |
| Default A: Dorsal Prefrontal Cortex 1 | 4.7 | 0.002 | 0.297 | 3.5 | 0.019 | 0.137 | 0.696 | 0.122 | 0.003 | 0.3 | 0.570 | 0.005 | 0.3 | 0.568 | 0.005 | 0.3 | 0.609 | 0.004 |
| Default A: Precuneus Posterior Cingulate Cortex 1 | 2.4 | 0.046 | 0.175 | 2.6 | 0.058 | 0.105 | | | | 0.5 | 0.481 | 0.007 | 0.9 | 0.346 | 0.013 | 0.1 | 0.799 | 0.001 |
| Default A: Medial Prefrontal Cortex 1 | 5.0 | 0.002 | 0.309 | 2.8 | 0.048 | 0.111 | 0.773 | 0.414 | 0.013 | 3.2 | 0.077 | 0.046 | 0.4 | 0.555 | 0.005 | 0.6 | 0.433 | 0.009 |
| Default B: Dorsal Prefrontal Cortex 1 | 4.7 | 0.002 | 0.295 | 1.6 | 0.201 | 0.066 | | | | 1.7 | 0.198 | 0.025 | 0.0 | 0.904 | 0.000 | 0.4 | 0.512 | 0.006 |
| Default B: Ventral Prefrontal Cortex 1 | 5.3 | 0.001 | 0.322 | 3.4 | 0.023 | 0.132 | 0.919 | 0.200 | 0.005 | 2.9 | 0.096 | 0.041 | 0.0 | 0.954 | 0.000 | 0.0 | 0.973 | 0.000 |
| Default B: Ventral Prefrontal Cortex 2 | 5.9 | 0.001 | 0.345 | 3.4 | 0.022 | 0.132 | 0.626 | 0.361 | 0.007 | 2.5 | 0.121 | 0.036 | 0.0 | 0.929 | 0.000 | 0.7 | 0.407 | 0.010 |
| Default C: Retro Superior Parietal Lobuleenial 1 | 2.0 | 0.077 | 0.155 | 0.9 | 0.424 | 0.041 | | | | 1.0 | 0.327 | 0.014 | 0.7 | 0.396 | 0.011 | 0.4 | 0.533 | 0.006 |
| Default C: Parahippocampal Cortex 1 | 3.2 | 0.013 | 0.221 | 2.7 | 0.052 | 0.108 | | | | 0.3 | 0.564 | 0.005 | 0.1 | 0.803 | 0.001 | 0.7 | 0.412 | 0.010 |
| Temporal Parietal 1 | 4.6 | 0.002 | 0.293 | 3.7 | 0.015 | 0.143 | 0.981 | 0.120 | 0.004 | 0.7 | 0.417 | 0.010 | 0.3 | 0.617 | 0.004 | 0.0 | 0.974 | 0.000 |
| Temporal Parietal 2 | 3.9 | 0.005 | 0.257 | 2.5 | 0.067 | 0.101 | | | | 2.7 | 0.106 | 0.039 | 0.0 | 0.980 | 0.000 | 0.0 | 0.962 | 0.000 |
| Temporal Parietal 3 | 2.4 | 0.041 | 0.179 | 2.0 | 0.117 | 0.084 | | | | 1.3 | 0.249 | 0.020 | 0.4 | 0.550 | 0.005 | 0.4 | 0.508 | 0.007 |

YIS = younger insulin sensitive; YIR = younger insulin resistant; OIS = older insulin sensitive; OIR = older insulin resistant

Table S3. General linear models of the association between regional cerebral blood flow among the four sub-groups based on age category and HOMA-IR2 levels, with blood pressure and cortical thickness as covariates. Post-hoc contrasts in the 42 regions with significant group difference are shown, comparing the young insulin sensitive group to the other three groups. The location of the 40 regions is plotted on the brain surface in Figure 2 in main document.

**Left Hemisphere**

| | Overall | | | 4 Groups: Age Cat and HOMAIR2 Median Split | | | Post-Hoc Contrasts | | | Systolic BP | | | Diastolic BP | | | Cortical Thickness | | |
|---|---|---|---|---|---|---|---|---|---|---|---|---|---|---|---|---|---|---|
| | F | p-FDR | $\eta^2_p$ | F | p | $\eta^2_p$ | YIS vs YIR | YIS vs OIS | YIS vs OIR | F | p | $\eta^2_p$ | F | p | $\eta^2_p$ | F | p | $\eta^2_p$ |
| Visual Central: Extra Striate Cortex 1 | 1.9 | 0.101 | 0.145 | 0.8 | 0.500 | 0.000 | | | | 0.3 | 0.567 | 0.005 | 1.1 | 0.307 | 0.016 | 0.0 | 0.945 | 0.000 |
| Visual Central: Extra Striate Cortex 2 | 5.8 | 0.001 | 0.343 | 2.9 | 0.043 | 0.000 | 0.649 | 0.553 | 0.027 | 2.3 | 0.135 | 0.033 | 2.2 | 0.141 | 0.032 | 0.3 | 0.585 | 0.004 |
| Visual Central: Striate Cortex 1 | 3.0 | 0.017 | 0.213 | 0.9 | 0.458 | 0.000 | | | | 2.2 | 0.146 | 0.031 | 0.3 | 0.576 | 0.005 | 1.8 | 0.188 | 0.026 |
| Visual Central: Extra Striate Cortex 3 | 4.0 | 0.005 | 0.265 | 2.9 | 0.043 | 0.000 | 0.819 | 0.605 | 0.017 | 2.5 | 0.122 | 0.035 | 0.2 | 0.692 | 0.002 | 0.0 | 0.934 | 0.000 |
| Visual Peripheral: Extra Striate Inferior 1 | 1.0 | 0.411 | 0.085 | 0.7 | 0.556 | 0.000 | | | | 0.5 | 0.466 | 0.008 | 0.0 | 0.966 | 0.000 | 0.2 | 0.634 | 0.003 |
| Visual Peripheral: Striate Cortex Calcarine 1 | 1.3 | 0.279 | 0.104 | 0.9 | 0.454 | 0.000 | | | | 1.2 | 0.278 | 0.018 | 0.0 | 0.952 | 0.000 | 0.0 | 0.905 | 0.000 |
| Visual Peripheral: Extra Striate Cortex Sup 1 | 2.8 | 0.025 | 0.200 | 2.5 | 0.069 | 0.000 | | | | 1.6 | 0.204 | 0.024 | 0.0 | 0.849 | 0.001 | 0.4 | 0.520 | 0.006 |
| Somatomotor A: 1 | 2.2 | 0.060 | 0.165 | 2.1 | 0.107 | 0.000 | | | | 0.1 | 0.701 | 0.002 | 0.4 | 0.531 | 0.006 | 0.4 | 0.524 | 0.006 |
| Somatomotor A: 2 | 2.2 | 0.057 | 0.167 | 2.1 | 0.103 | 0.000 | | | | 0.0 | 0.829 | 0.001 | 0.1 | 0.779 | 0.001 | 0.1 | 0.726 | 0.002 |
| Somatomotor B: Auditory 1 | 1.8 | 0.111 | 0.141 | 1.7 | 0.173 | 0.000 | | | | 1.3 | 0.257 | 0.019 | 0.0 | 0.961 | 0.000 | 0.0 | 0.942 | 0.000 |
| Somatomotor B: S2 1 | 3.7 | 0.007 | 0.246 | 3.0 | 0.038 | 0.000 | 0.759 | 0.898 | 0.030 | 2.0 | 0.157 | 0.030 | 0.2 | 0.670 | 0.003 | 0.1 | 0.777 | 0.001 |
| Somatomotor B: S2 2 | 2.5 | 0.038 | 0.184 | 2.2 | 0.093 | 0.000 | | | | 1.1 | 0.301 | 0.016 | 0.1 | 0.805 | 0.001 | 0.7 | 0.403 | 0.010 |
| Somatomotor B: Central 1 | 2.1 | 0.069 | 0.160 | 1.9 | 0.138 | 0.000 | | | | 1.2 | 0.284 | 0.017 | 0.1 | 0.707 | 0.002 | 0.1 | 0.765 | 0.001 |
| Dorsal Attention A: Temporal Occipital 1 | 3.9 | 0.005 | 0.257 | 2.2 | 0.096 | 0.000 | | | | 0.5 | 0.480 | 0.007 | 2.0 | 0.166 | 0.028 | 0.1 | 0.783 | 0.001 |
| Dorsal Attention A: Parietal Occipital 1 | 3.5 | 0.008 | 0.239 | 2.8 | 0.046 | 0.000 | 0.666 | 0.889 | 0.058 | 2.6 | 0.114 | 0.037 | 0.5 | 0.463 | 0.008 | 1.2 | 0.281 | 0.017 |
| Dorsal Attention A: Superior Parietal Lobule 1 | 5.5 | 0.001 | 0.330 | 4.6 | 0.005 | 0.000 | 0.657 | 0.061 | 0.001 | 2.0 | 0.166 | 0.028 | 0.2 | 0.663 | 0.003 | 2.1 | 0.153 | 0.030 |
| Dorsal Attention B: Post Central 1 | 2.0 | 0.087 | 0.151 | 1.7 | 0.183 | 0.000 | | | | 0.8 | 0.366 | 0.012 | 0.1 | 0.777 | 0.001 | 0.0 | 0.993 | 0.000 |
| Dorsal Attention B: Post Central 2 | 2.6 | 0.032 | 0.191 | 3.0 | 0.037 | 0.000 | 0.715 | 0.173 | 0.007 | 0.4 | 0.538 | 0.006 | 0.6 | 0.445 | 0.009 | 2.4 | 0.123 | 0.035 |
| Dorsal Attention B: Post Central 3 | 3.7 | 0.007 | 0.247 | 3.5 | 0.020 | 0.000 | 0.258 | 0.025 | 0.002 | 1.6 | 0.204 | 0.024 | 0.1 | 0.745 | 0.002 | 4.0 | 0.049 | 0.056 |
| Dorsal Attention B: Frontal Eye Fields 1 | 3.1 | 0.016 | 0.215 | 1.1 | 0.346 | 0.000 | | | | 0.1 | 0.728 | 0.002 | 0.4 | 0.541 | 0.006 | 1.6 | 0.204 | 0.024 |
| Salience Ventral Attention A: Parietal Operculum 1 | 4.2 | 0.004 | 0.274 | 3.3 | 0.026 | 0.000 | 0.954 | 0.288 | 0.008 | 0.5 | 0.485 | 0.007 | 1.2 | 0.268 | 0.018 | 0.0 | 0.835 | 0.001 |
| Salience Ventral Attention A: Insula: 1 | 3.5 | 0.008 | 0.239 | 2.0 | 0.120 | 0.000 | | | | 0.2 | 0.656 | 0.003 | 0.2 | 0.628 | 0.004 | 1.6 | 0.212 | 0.023 |
| Salience Ventral Attention A: Insula: 2 | 4.0 | 0.005 | 0.263 | 3.1 | 0.034 | 0.000 | 0.863 | 0.126 | 0.011 | 2.0 | 0.160 | 0.029 | 0.1 | 0.745 | 0.002 | 0.8 | 0.370 | 0.012 |
| Salience Ventral Attention A: Parietal Medial 1 | 4.3 | 0.004 | 0.276 | 4.0 | 0.011 | 0.000 | 0.982 | 0.194 | 0.004 | 0.5 | 0.478 | 0.008 | 1.1 | 0.299 | 0.016 | 0.8 | 0.378 | 0.012 |
| Salience Ventral Attention A: Frontal Medial 1 | 2.3 | 0.051 | 0.172 | 0.8 | 0.478 | 0.000 | | | | 0.0 | 0.921 | 0.000 | 0.3 | 0.562 | 0.005 | 0.8 | 0.366 | 0.012 |
| Salience Ventral Attention B: Lateral Prefrontal Cortex 1 | 3.6 | 0.008 | 0.242 | 1.9 | 0.135 | 0.000 | | | | 2.9 | 0.092 | 0.042 | 0.1 | 0.800 | 0.001 | 0.2 | 0.692 | 0.002 |
| Salience Ventral Attention B: Medial Posterior Prefrontal 1 | 4.7 | 0.002 | 0.297 | 5.7 | 0.001 | 0.000 | 0.894 | 0.027 | 0.000 | 0.0 | 0.979 | 0.000 | 0.6 | 0.437 | 0.009 | 2.6 | 0.110 | 0.038 |
| LimbiC: B: Orbital Frontal Cortex 1 | 6.6 | 0.000 | 0.371 | 4.4 | 0.007 | 0.000 | 0.645 | 0.045 | 0.001 | 3.0 | 0.090 | 0.042 | 0.0 | 0.854 | 0.001 | 5.1 | 0.027 | 0.071 |
| LimbiC: A: Temporal Pole 1 | 8.5 | 0.000 | 0.434 | 4.5 | 0.006 | 0.000 | 0.399 | 0.024 | 0.001 | 2.3 | 0.136 | 0.033 | 0.5 | 0.467 | 0.008 | 4.5 | 0.038 | 0.063 |
| LimbiC: A: Temporal Pole 2 | 6.3 | 0.000 | 0.360 | 5.4 | 0.002 | 0.000 | 0.693 | 0.034 | 0.002 | 0.2 | 0.692 | 0.002 | 2.7 | 0.103 | 0.039 | 0.2 | 0.619 | 0.004 |
| Control A: Intraparietal Sulcus 1 | 4.3 | 0.003 | 0.280 | 3.5 | 0.021 | 0.000 | 0.925 | 0.186 | 0.005 | 0.6 | 0.432 | 0.009 | 1.0 | 0.324 | 0.015 | 0.0 | 0.835 | 0.001 |
| Control A: Lateral Prefrontal Cortex 1 | 4.3 | 0.004 | 0.277 | 3.3 | 0.026 | 0.000 | 0.842 | 0.193 | 0.007 | 3.0 | 0.090 | 0.042 | 0.1 | 0.784 | 0.001 | 1.2 | 0.287 | 0.017 |
| Control A: Lateral Prefrontal Cortex 2 | 4.4 | 0.003 | 0.281 | 2.8 | 0.047 | 0.000 | 0.702 | 0.321 | 0.026 | 3.1 | 0.084 | 0.044 | 0.0 | 0.952 | 0.000 | 0.6 | 0.444 | 0.009 |
| Control B: Lateral Prefrontal Cortex 1 | 8.0 | 0.000 | 0.418 | 4.6 | 0.005 | 0.000 | 0.417 | 0.105 | 0.001 | 8.3 | 0.005 | 0.111 | 0.3 | 0.573 | 0.005 | 1.6 | 0.217 | 0.023 |
| Control C: Precuneus 1 | 3.5 | 0.008 | 0.240 | 3.6 | 0.017 | 0.000 | 0.339 | 0.112 | 0.002 | 0.8 | 0.364 | 0.012 | 0.2 | 0.627 | 0.004 | 1.8 | 0.186 | 0.026 |
| Control C: Precuneus 2 | 2.6 | 0.035 | 0.187 | 3.2 | 0.027 | 0.000 | 0.331 | 0.121 | 0.003 | 0.6 | 0.455 | 0.008 | 0.0 | 0.883 | 0.000 | 2.5 | 0.115 | 0.037 |
| Control C: Cingulate Posterior 1 | 4.9 | 0.002 | 0.306 | 5.2 | 0.003 | 0.000 | 0.840 | 0.270 | 0.001 | 1.2 | 0.268 | 0.018 | 0.5 | 0.467 | 0.008 | 1.8 | 0.180 | 0.027 |
| Default A: Dorsal Prefrontal Cortex 1 | 3.7 | 0.007 | 0.248 | 2.3 | 0.083 | 0.000 | | | | 1.7 | 0.197 | 0.025 | 0.0 | 0.897 | 0.000 | 0.0 | 0.836 | 0.001 |
| Default A: Precuneus Posterior Cingulate Cortex1 | 2.7 | 0.027 | 0.197 | 2.6 | 0.058 | 0.000 | | | | 1.1 | 0.294 | 0.016 | 0.1 | 0.709 | 0.002 | 0.1 | 0.816 | 0.001 |
| Default A: Medial Prefrontal Cortex 1 | 5.7 | 0.001 | 0.337 | 5.5 | 0.002 | 0.000 | 0.866 | 0.100 | 0.001 | 1.1 | 0.292 | 0.017 | 0.0 | 0.921 | 0.000 | 0.3 | 0.616 | 0.004 |
| Default B: Temp 1 | 4.5 | 0.003 | 0.287 | 1.5 | 0.220 | 0.000 | | | | 1.4 | 0.248 | 0.020 | 0.2 | 0.668 | 0.003 | 1.5 | 0.219 | 0.022 |
| Default B: Temp 2 | 3.1 | 0.015 | 0.217 | 3.1 | 0.034 | 0.000 | 0.761 | 0.257 | 0.009 | 1.1 | 0.295 | 0.016 | 0.1 | 0.709 | 0.002 | 1.8 | 0.185 | 0.026 |
| Default B: Inferior Parietal Lobule 1 | 4.7 | 0.002 | 0.296 | 2.8 | 0.045 | 0.000 | 0.977 | 0.486 | 0.019 | 2.2 | 0.144 | 0.032 | 0.7 | 0.423 | 0.010 | 0.1 | 0.786 | 0.001 |
| Default B: Dorsal Prefrontal Cortex 1 | 3.7 | 0.007 | 0.248 | 1.1 | 0.351 | 0.000 | | | | 1.7 | 0.200 | 0.024 | 0.3 | 0.600 | 0.004 | 0.3 | 0.608 | 0.004 |
| Default B: Lateral Prefrontal Cortex 1 | 2.6 | 0.033 | 0.190 | 1.6 | 0.204 | 0.000 | | | | 2.1 | 0.150 | 0.031 | 0.0 | 0.979 | 0.000 | 0.8 | 0.369 | 0.012 |
| Default B: Ventral Prefrontal Cortex 1 | 4.1 | 0.004 | 0.270 | 4.5 | 0.007 | 0.000 | 0.548 | 0.074 | 0.006 | 0.3 | 0.578 | 0.005 | 0.1 | 0.767 | 0.001 | 1.4 | 0.246 | 0.020 |
| Default B: Ventral Prefrontal Cortex 2 | 8.1 | 0.000 | 0.421 | 5.3 | 0.002 | 0.000 | 0.836 | 0.043 | 0.000 | 3.1 | 0.081 | 0.045 | 0.0 | 0.904 | 0.000 | 0.1 | 0.767 | 0.001 |
| Default C: Retro Superior Parietal Lobuleenial 1 | 3.4 | 0.009 | 0.235 | 1.4 | 0.260 | 0.000 | | | | 1.7 | 0.194 | 0.025 | 0.5 | 0.479 | 0.008 | 1.7 | 0.196 | 0.025 |
| Default C: Parahippocampal Cortex 1 | 2.4 | 0.044 | 0.178 | 1.8 | 0.152 | 0.000 | | | | 0.1 | 0.790 | 0.001 | 0.2 | 0.662 | 0.003 | 1.0 | 0.320 | 0.015 |
| Temporal Parietal 1 | 2.3 | 0.052 | 0.171 | 2.7 | 0.056 | 0.000 | | | | 1.1 | 0.297 | 0.016 | 0.2 | 0.671 | 0.003 | 1.0 | 0.323 | 0.015 |

**Right Hemisphere**

| | Overall | | | 4 Groups: Age Cat and HOMAIR2 Median Split | | | Post-Hoc Contrasts | | | Systolic BP | | | Diastolic BP | | | Cortical Thickness | | |
|---|---|---|---|---|---|---|---|---|---|---|---|---|---|---|---|---|---|---|
| | F | p-FDR | $\eta^2_p$ | F | p | $\eta^2_p$ | YIS vs YIR | YIS vs OIS | YIS vs OIR | F | p | $\eta^2_p$ | F | p | $\eta^2_p$ | F | p | $\eta^2_p$ |
| Visual Central: Extra Striate Cortex 1 | 1.0 | 0.450 | 0.080 | 1.0 | 0.420 | 0.000 | | | | 0.3 | 0.557 | 0.005 | 0.0 | 0.998 | 0.000 | 0.0 | 0.888 | 0.000 |
| Visual Central: Extra Striate Cortex 2 | 4.4 | 0.003 | 0.282 | 3.9 | 0.012 | 0.000 | 0.809 | 0.288 | 0.004 | 0.6 | 0.425 | 0.010 | 1.2 | 0.281 | 0.017 | 0.2 | 0.678 | 0.003 |
| Visual Central: Extra Striate Cortex 3 | 3.9 | 0.005 | 0.259 | 3.3 | 0.025 | 0.000 | 0.740 | 0.734 | 0.014 | 3.2 | 0.077 | 0.046 | 0.1 | 0.730 | 0.002 | 1.0 | 0.328 | 0.014 |
| Visual Peripheral: Striate Cortex Calcarine 1 | 1.2 | 0.306 | 0.100 | 1.0 | 0.377 | 0.000 | | | | 0.8 | 0.368 | 0.012 | 0.0 | 0.941 | 0.000 | 0.1 | 0.786 | 0.001 |
| Visual Peripheral: Extra Striate Inferior 1 | 1.0 | 0.410 | 0.086 | 1.0 | 0.412 | 0.000 | | | | 0.9 | 0.337 | 0.014 | 0.0 | 0.948 | 0.000 | 0.1 | 0.702 | 0.002 |
| Visual Peripheral: Extra Striate Superior 1 | 2.0 | 0.087 | 0.150 | 2.1 | 0.105 | 0.000 | | | | 0.6 | 0.439 | 0.009 | 0.3 | 0.586 | 0.004 | 0.2 | 0.622 | 0.004 |
| Somatomotor A: 1 | 2.5 | 0.038 | 0.184 | 2.0 | 0.127 | 0.000 | | | | 0.3 | 0.573 | 0.005 | 0.5 | 0.489 | 0.007 | 4.0 | 0.051 | 0.056 |
| Somatomotor A: 2 | 2.0 | 0.083 | 0.153 | 1.5 | 0.216 | 0.000 | | | | 0.1 | 0.751 | 0.002 | 0.5 | 0.492 | 0.007 | 1.7 | 0.191 | 0.025 |
| Somatomotor A: 3 | 2.4 | 0.047 | 0.176 | 1.3 | 0.283 | 0.000 | | | | 0.9 | 0.343 | 0.013 | 0.0 | 0.849 | 0.001 | 0.1 | 0.737 | 0.002 |
| Somatomotor A: 4 | 2.0 | 0.079 | 0.155 | 1.1 | 0.370 | 0.000 | | | | 0.1 | 0.781 | 0.001 | 0.4 | 0.530 | 0.006 | 1.1 | 0.294 | 0.016 |
| Somatomotor B: Auditory 1 | 3.0 | 0.017 | 0.212 | 2.0 | 0.119 | 0.000 | | | | 2.5 | 0.118 | 0.036 | 0.0 | 0.879 | 0.000 | 0.5 | 0.482 | 0.007 |
| Somatomotor B: S2 1 | 2.6 | 0.034 | 0.188 | 1.6 | 0.208 | 0.000 | | | | 2.1 | 0.156 | 0.030 | 0.1 | 0.815 | 0.001 | 0.0 | 1.000 | 0.000 |
| Somatomotor B: S2 2 | 3.7 | 0.007 | 0.249 | 1.1 | 0.337 | 0.000 | | | | 3.4 | 0.070 | 0.048 | 0.4 | 0.523 | 0.006 | 2.9 | 0.091 | 0.042 |
| Somatomotor B: Central 1 | 2.3 | 0.056 | 0.169 | 1.9 | 0.136 | 0.000 | | | | 1.6 | 0.208 | 0.024 | 0.0 | 0.960 | 0.000 | 0.4 | 0.534 | 0.006 |
| Dorsal Attention A: Temporal Occipital 1 | 3.7 | 0.007 | 0.249 | 2.4 | 0.074 | 0.000 | | | | 0.9 | 0.344 | 0.013 | 1.2 | 0.284 | 0.017 | 0.1 | 0.779 | 0.001 |
| Dorsal Attention A: Parietal Occipital 1 | 2.9 | 0.019 | 0.208 | 2.5 | 0.069 | 0.000 | | | | 1.0 | 0.327 | 0.014 | 0.7 | 0.398 | 0.011 | 0.0 | 0.856 | 0.000 |
| Dorsal Attention A: Superior Parietal Lobule 1 | 4.0 | 0.005 | 0.264 | 4.2 | 0.009 | 0.000 | 0.400 | 0.174 | 0.002 | 0.7 | 0.420 | 0.010 | 0.7 | 0.403 | 0.010 | 1.1 | 0.302 | 0.016 |
| Dorsal Attention B: Post Central 1 | 3.1 | 0.015 | 0.218 | 2.8 | 0.047 | 0.000 | 0.736 | 0.473 | 0.019 | 2.3 | 0.136 | 0.033 | 0.2 | 0.674 | 0.003 | 1.6 | 0.208 | 0.024 |
| Dorsal Attention B: Post Central 2 | 2.8 | 0.024 | 0.202 | 2.7 | 0.052 | 0.000 | | | | 0.4 | 0.553 | 0.005 | 0.4 | 0.521 | 0.006 | 1.1 | 0.295 | 0.016 |
| Dorsal Attention B: Frontal Eye Fields 1 | 3.2 | 0.013 | 0.224 | 2.5 | 0.068 | 0.000 | | | | 0.1 | 0.792 | 0.001 | 0.3 | 0.615 | 0.004 | 0.3 | 0.614 | 0.004 |
| Salience Ventral Attention A: Parietal Operculum 1 | 3.9 | 0.005 | 0.260 | 2.3 | 0.086 | 0.000 | | | | 2.2 | 0.144 | 0.032 | 0.0 | 0.943 | 0.000 | 0.9 | 0.334 | 0.014 |
| Salience Ventral Attention A: Insula: 1 | 3.6 | 0.008 | 0.243 | 1.8 | 0.157 | 0.000 | | | | 1.7 | 0.201 | 0.024 | 0.0 | 0.919 | 0.000 | 0.3 | 0.583 | 0.005 |
| Salience Ventral Attention A: Parietal Medial 1 | 3.4 | 0.010 | 0.231 | 3.3 | 0.024 | 0.000 | 0.687 | 0.446 | 0.015 | 0.8 | 0.382 | 0.011 | 0.6 | 0.425 | 0.010 | 1.8 | 0.182 | 0.026 |
| Salience Ventral Attention A: Frontal Medial 1 | 3.2 | 0.013 | 0.224 | 1.7 | 0.166 | 0.000 | | | | 0.6 | 0.455 | 0.008 | 0.4 | 0.526 | 0.006 | 0.0 | 0.943 | 0.000 |
| Salience Ventral Attention B: Inferior Parietal Lobule 1 | 4.2 | 0.004 | 0.272 | 1.7 | 0.168 | 0.000 | | | | 1.5 | 0.218 | 0.023 | 0.2 | 0.634 | 0.003 | 0.9 | 0.336 | 0.014 |
| Salience Ventral Attention B: Lateral Prefrontal Cortex 1 | 4.2 | 0.004 | 0.272 | 0.9 | 0.460 | 0.000 | | | | 2.9 | 0.096 | 0.041 | 0.0 | 0.861 | 0.000 | 2.0 | 0.161 | 0.029 |
| Salience Ventral Attention B: Medial Posterior Prefrontal 1 | 4.7 | 0.002 | 0.294 | 4.9 | 0.004 | 0.000 | 0.995 | 0.065 | 0.001 | 0.6 | 0.453 | 0.008 | 0.1 | 0.721 | 0.002 | 3.7 | 0.059 | 0.052 |
| LimbiC: B: Orbital Frontal Cortex 1 | 5.4 | 0.001 | 0.325 | 3.3 | 0.025 | 0.000 | 0.402 | 0.084 | 0.003 | 3.6 | 0.062 | 0.051 | 0.2 | 0.658 | 0.003 | 0.9 | 0.355 | 0.013 |
| LimbiC: A: Temporal Pole 1 | 8.9 | 0.000 | 0.444 | 6.4 | 0.001 | 0.000 | 0.049 | 0.001 | 0.000 | 2.0 | 0.162 | 0.029 | 0.3 | 0.564 | 0.005 | 1.2 | 0.285 | 0.017 |
| Control A: Intraparietal Sulcus 1 | 4.1 | 0.004 | 0.266 | 2.5 | 0.069 | 0.000 | | | | 1.8 | 0.187 | 0.026 | 1.1 | 0.309 | 0.015 | 0.0 | 0.913 | 0.000 |
| Control A: Lateral Prefrontal Cortex 1 | 4.8 | 0.002 | 0.302 | 2.7 | 0.051 | 0.000 | | | | 3.3 | 0.072 | 0.047 | 0.0 | 0.958 | 0.000 | 0.0 | 0.947 | 0.000 |
| Control A: Lateral Prefrontal Cortex 2 | 5.2 | 0.001 | 0.316 | 2.4 | 0.072 | 0.000 | | | | 0.6 | 0.445 | 0.009 | 0.4 | 0.544 | 0.006 | 1.8 | 0.184 | 0.026 |
| Control B: Temporal 1 | 6.2 | 0.000 | 0.357 | 3.0 | 0.035 | 0.000 | 0.288 | 0.071 | 0.004 | 3.2 | 0.079 | 0.045 | 0.0 | 0.932 | 0.000 | 0.9 | 0.359 | 0.013 |
| Control B: inferior parietal lobule 1 | 3.1 | 0.014 | 0.220 | 2.4 | 0.078 | 0.000 | | | | 0.8 | 0.364 | 0.012 | 0.8 | 0.388 | 0.011 | 0.0 | 0.833 | 0.001 |
| Control B: Lateral Prefrontal Cortexd 1 | 3.5 | 0.009 | 0.237 | 1.5 | 0.222 | 0.000 | | | | 1.0 | 0.326 | 0.014 | 0.1 | 0.758 | 0.001 | 0.4 | 0.543 | 0.006 |
| Control B: Lateral Prefrontal Cortexv 1 | 7.1 | 0.000 | 0.388 | 3.0 | 0.038 | 0.000 | 0.297 | 0.249 | 0.006 | 9.1 | 0.004 | 0.120 | 0.3 | 0.617 | 0.004 | 0.0 | 0.827 | 0.001 |
| Control C: Cingulate Posterior 1 | 4.2 | 0.004 | 0.273 | 4.5 | 0.006 | 0.000 | 0.882 | 0.819 | 0.008 | 0.6 | 0.425 | 0.010 | 0.9 | 0.355 | 0.013 | 0.0 | 0.835 | 0.001 |
| Control C: Precuneus 1 | 2.6 | 0.034 | 0.189 | 2.7 | 0.050 | 0.000 | | | | 0.2 | 0.632 | 0.003 | 1.1 | 0.295 | 0.016 | 0.0 | 0.917 | 0.000 |
| Default A: Inferior Parietal Lobule 1 | 3.2 | 0.014 | 0.220 | 2.4 | 0.078 | 0.000 | | | | 1.2 | 0.283 | 0.017 | 0.2 | 0.645 | 0.003 | 0.0 | 0.916 | 0.000 |
| Default A: Dorsal Prefrontal Cortex 1 | 4.6 | 0.002 | 0.292 | 3.4 | 0.024 | 0.000 | 0.573 | 0.082 | 0.003 | 0.4 | 0.548 | 0.005 | 0.2 | 0.629 | 0.004 | 0.2 | 0.666 | 0.003 |
| Default A: Precuneus Posterior Cingulate Cortex 1 | 2.4 | 0.047 | 0.175 | 2.6 | 0.057 | 0.000 | | | | 0.5 | 0.488 | 0.007 | 0.7 | 0.402 | 0.010 | 0.1 | 0.793 | 0.001 |
| Default A: Medial Prefrontal Cortex 1 | 5.0 | 0.002 | 0.307 | 2.7 | 0.050 | 0.000 | | | | 3.2 | 0.077 | 0.046 | 0.3 | 0.561 | 0.005 | 0.7 | 0.413 | 0.010 |
| Default B: Dorsal Prefrontal Cortex 1 | 4.6 | 0.003 | 0.290 | 1.4 | 0.243 | 0.000 | | | | 1.7 | 0.197 | 0.025 | 0.1 | 0.795 | 0.001 | 0.5 | 0.479 | 0.008 |
| Default B: Ventral Prefrontal Cortex 1 | 5.2 | 0.001 | 0.318 | 3.3 | 0.027 | 0.000 | 0.774 | 0.162 | 0.006 | 2.8 | 0.101 | 0.040 | 0.0 | 0.982 | 0.000 | 0.0 | 0.900 | 0.000 |
| Default B: Ventral Prefrontal Cortex 2 | 5.8 | 0.001 | 0.343 | 3.3 | 0.025 | 0.000 | 0.286 | 0.159 | 0.004 | 2.6 | 0.111 | 0.037 | 0.0 | 0.932 | 0.000 | 0.4 | 0.512 | 0.006 |
| Default C: Retro Superior Parietal Lobuleenial 1 | 2.2 | 0.062 | 0.164 | 1.2 | 0.317 | 0.000 | | | | 1.0 | 0.312 | 0.015 | 0.4 | 0.513 | 0.006 | 0.7 | 0.414 | 0.010 |
| Default C: Parahippocampal Cortex 1 | 3.3 | 0.012 | 0.228 | 2.9 | 0.040 | 0.000 | 0.276 | 0.043 | 0.005 | 0.4 | 0.536 | 0.006 | 0.3 | 0.609 | 0.004 | 0.7 | 0.412 | 0.010 |
| Temporal Parietal 1 | 4.7 | 0.002 | 0.295 | 3.8 | 0.014 | 0.000 | 0.944 | 0.099 | 0.003 | 0.7 | 0.423 | 0.010 | 0.2 | 0.627 | 0.004 | 0.0 | 0.924 | 0.000 |
| Temporal Parietal 2 | 3.9 | 0.005 | 0.257 | 2.5 | 0.068 | 0.000 | | | | 2.7 | 0.108 | 0.038 | 0.0 | 0.962 | 0.000 | 0.0 | 0.994 | 0.000 |
| Temporal Parietal 3 | 2.4 | 0.047 | 0.176 | 1.9 | 0.131 | 0.000 | | | | 1.3 | 0.251 | 0.020 | 0.3 | 0.600 | 0.004 | 0.4 | 0.546 | 0.005 |

YIS = younger insulin sensitive; YIR = younger insulin resistant; OIS = older insulin sensitive; OIR = older insulin resistant

Table S4. General linear models of the association between regional cerebral blood flow and age category, blood pressure, cortical thickness, resting heart rate, BMI, sex, years of education.

***Left Hemisphere***

| | Overall Model | | | Age Category | | | Systolic BP | | | Diastolic BP | | | Resting HR | | | BMI | | | Years of Education | | | Sex | | |
|---|---|---|---|---|---|---|---|---|---|---|---|---|---|---|---|---|---|---|---|---|---|---|---|---|
| | F | p-FDR | $\eta^2_p$ | F | p | $\eta^2_p$ | F | p | $\eta^2_p$ | F | p | $\eta^2_p$ | F | p | $\eta^2_p$ | F | p | $\eta^2_p$ | F | p | $\eta^2_p$ | F | p | $\eta^2_p$ |
| Visual Central: Extra Striate Cortex 1 | 4.8 | 0.000 | 0.417 | 0.6 | 0.461 | 0.010 | 0.0 | 0.999 | 0.000 | 2.3 | 0.131 | 0.042 | 0.2 | 0.679 | 0.003 | 0.4 | 0.537 | 0.007 | 5.9 | 0.018 | 0.099 | 14.2 | 0.000 | 0.208 |
| Visual Central: Extra Striate Cortex 2 | 6.2 | 0.000 | 0.479 | 4.1 | 0.047 | 0.071 | 0.1 | 0.763 | 0.002 | 2.2 | 0.142 | 0.040 | 0.1 | 0.726 | 0.002 | 1.4 | 0.244 | 0.025 | 1.1 | 0.304 | 0.020 | 14.6 | 0.000 | 0.213 |
| Visual Central: Striate Cortex 1 | 5.2 | 0.000 | 0.436 | 0.8 | 0.388 | 0.014 | 0.6 | 0.442 | 0.011 | 0.4 | 0.554 | 0.007 | 0.0 | 0.993 | 0.000 | 1.8 | 0.184 | 0.032 | 0.4 | 0.513 | 0.008 | 19.1 | 0.000 | 0.261 |
| Visual Central: Extra Striate Cortex 3 | 5.2 | 0.000 | 0.433 | 1.8 | 0.184 | 0.032 | 1.6 | 0.217 | 0.028 | 0.1 | 0.721 | 0.002 | 0.1 | 0.723 | 0.002 | 3.7 | 0.060 | 0.064 | 0.5 | 0.491 | 0.009 | 14.2 | 0.000 | 0.209 |
| Visual Peripheral: Extra Striate Inferior 1 | 3.5 | 0.003 | 0.339 | 0.3 | 0.603 | 0.005 | 0.0 | 0.858 | 0.001 | 0.1 | 0.762 | 0.002 | 0.1 | 0.718 | 0.002 | 1.7 | 0.198 | 0.030 | 2.6 | 0.115 | 0.045 | 14.6 | 0.000 | 0.212 |
| Visual Peripheral: Striate Cortex Calcarine 1 | 3.9 | 0.002 | 0.364 | 0.0 | 0.854 | 0.001 | 0.3 | 0.613 | 0.005 | 0.0 | 0.976 | 0.000 | 0.9 | 0.352 | 0.016 | 3.7 | 0.061 | 0.064 | 1.7 | 0.202 | 0.030 | 17.7 | 0.000 | 0.247 |
| Visual Peripheral: Extra Striate Cortex Sup 1 | 4.7 | 0.000 | 0.411 | 0.3 | 0.616 | 0.005 | 1.3 | 0.253 | 0.024 | 0.7 | 0.414 | 0.012 | 1.4 | 0.241 | 0.025 | 5.8 | 0.019 | 0.098 | 0.4 | 0.544 | 0.007 | 15.6 | 0.000 | 0.225 |
| Somatomotor A: 1 | 3.3 | 0.004 | 0.329 | 0.8 | 0.374 | 0.015 | 0.0 | 0.921 | 0.000 | 0.1 | 0.789 | 0.001 | 0.8 | 0.390 | 0.014 | 0.2 | 0.625 | 0.004 | 0.2 | 0.680 | 0.003 | 14.9 | 0.000 | 0.216 |
| Somatomotor A: 2 | 4.3 | 0.001 | 0.387 | 1.7 | 0.196 | 0.031 | 0.0 | 0.841 | 0.001 | 0.0 | 0.988 | 0.000 | 1.1 | 0.305 | 0.019 | 0.9 | 0.340 | 0.017 | 0.1 | 0.743 | 0.002 | 18.8 | 0.000 | 0.258 |
| Somatomotor B: Auditory 1 | 2.4 | 0.029 | 0.260 | 0.6 | 0.428 | 0.012 | 0.2 | 0.634 | 0.004 | 0.4 | 0.542 | 0.007 | 1.1 | 0.303 | 0.020 | 3.3 | 0.075 | 0.057 | 2.3 | 0.134 | 0.041 | 6.7 | 0.012 | 0.111 |
| Somatomotor B: S2 1 | 4.1 | 0.001 | 0.376 | 0.8 | 0.385 | 0.014 | 0.4 | 0.551 | 0.007 | 0.1 | 0.748 | 0.002 | 0.3 | 0.609 | 0.005 | 4.5 | 0.038 | 0.077 | 1.0 | 0.316 | 0.019 | 12.1 | 0.001 | 0.183 |
| Somatomotor B: S2 2 | 2.8 | 0.011 | 0.297 | 1.2 | 0.282 | 0.021 | 0.6 | 0.445 | 0.011 | 0.2 | 0.628 | 0.004 | 0.3 | 0.586 | 0.006 | 3.1 | 0.086 | 0.054 | 1.2 | 0.283 | 0.021 | 7.5 | 0.008 | 0.122 |
| Somatomotor B: Central 1 | 4.6 | 0.000 | 0.407 | 0.0 | 0.912 | 0.000 | 1.6 | 0.214 | 0.028 | 0.3 | 0.594 | 0.005 | 1.3 | 0.257 | 0.024 | 3.6 | 0.064 | 0.062 | 1.4 | 0.245 | 0.025 | 17.8 | 0.000 | 0.248 |
| Dorsal Attention A: Temporal Occipital 1 | 5.7 | 0.000 | 0.456 | 2.1 | 0.149 | 0.038 | 0.1 | 0.718 | 0.002 | 0.6 | 0.429 | 0.012 | 0.2 | 0.698 | 0.003 | 4.3 | 0.043 | 0.074 | 2.6 | 0.111 | 0.046 | 13.9 | 0.000 | 0.205 |
| Dorsal Attention A: Parietal Occipital 1 | 3.1 | 0.006 | 0.316 | 1.0 | 0.313 | 0.019 | 0.6 | 0.448 | 0.011 | 0.0 | 0.981 | 0.000 | 0.3 | 0.559 | 0.006 | 3.5 | 0.065 | 0.062 | 1.8 | 0.189 | 0.032 | 8.0 | 0.006 | 0.129 |
| Dorsal Attention A: Superior Parietal Lobule 1 | 8.3 | 0.000 | 0.553 | 6.3 | 0.015 | 0.105 | 1.4 | 0.239 | 0.026 | 0.0 | 0.833 | 0.001 | 0.4 | 0.509 | 0.008 | 2.3 | 0.137 | 0.040 | 0.5 | 0.498 | 0.009 | 26.4 | 0.000 | 0.328 |
| Dorsal Attention B: Post Central 1 | 3.3 | 0.005 | 0.328 | 0.9 | 0.335 | 0.017 | 0.7 | 0.410 | 0.013 | 0.7 | 0.404 | 0.013 | 0.6 | 0.459 | 0.010 | 1.6 | 0.216 | 0.028 | 0.5 | 0.470 | 0.010 | 12.3 | 0.001 | 0.185 |
| Dorsal Attention B: Post Central 2 | 3.7 | 0.002 | 0.357 | 1.3 | 0.264 | 0.023 | 0.4 | 0.550 | 0.007 | 0.0 | 0.902 | 0.000 | 0.8 | 0.381 | 0.014 | 3.0 | 0.091 | 0.052 | 0.2 | 0.700 | 0.003 | 15.0 | 0.000 | 0.217 |
| Dorsal Attention B: Post Central 3 | 5.5 | 0.000 | 0.451 | 5.4 | 0.025 | 0.090 | 0.8 | 0.379 | 0.014 | 0.6 | 0.448 | 0.011 | 1.7 | 0.199 | 0.030 | 0.9 | 0.352 | 0.016 | 0.0 | 0.882 | 0.000 | 18.4 | 0.000 | 0.254 |
| Dorsal Attention B: Frontal Eye Fields 1 | 4.5 | 0.001 | 0.401 | 1.5 | 0.223 | 0.027 | 0.0 | 0.865 | 0.001 | 0.3 | 0.606 | 0.005 | 0.3 | 0.601 | 0.005 | 0.5 | 0.478 | 0.009 | 0.8 | 0.386 | 0.014 | 13.5 | 0.001 | 0.200 |
| Salience Ventral Attention A: Parietal Operculum 1 | 5.4 | 0.000 | 0.445 | 3.7 | 0.059 | 0.065 | 0.2 | 0.682 | 0.003 | 0.4 | 0.526 | 0.007 | 0.0 | 0.985 | 0.000 | 2.3 | 0.138 | 0.040 | 1.0 | 0.319 | 0.018 | 15.4 | 0.000 | 0.222 |
| Salience Ventral Attention A: Insula: 1 | 5.6 | 0.000 | 0.454 | 4.0 | 0.050 | 0.069 | 0.2 | 0.627 | 0.004 | 0.0 | 0.831 | 0.001 | 1.0 | 0.314 | 0.019 | 1.9 | 0.177 | 0.034 | 2.9 | 0.094 | 0.051 | 16.6 | 0.000 | 0.235 |
| Salience Ventral Attention A: Insula: 2 | 5.1 | 0.000 | 0.430 | 1.7 | 0.200 | 0.030 | 1.3 | 0.256 | 0.024 | 1.2 | 0.286 | 0.021 | 0.5 | 0.481 | 0.009 | 4.5 | 0.039 | 0.077 | 0.8 | 0.373 | 0.015 | 12.4 | 0.001 | 0.187 |
| Salience Ventral Attention A: Parietal Medial 1 | 5.4 | 0.000 | 0.444 | 1.4 | 0.243 | 0.025 | 1.3 | 0.258 | 0.024 | 0.0 | 0.950 | 0.000 | 0.4 | 0.553 | 0.007 | 5.1 | 0.028 | 0.086 | 0.0 | 0.945 | 0.000 | 12.2 | 0.001 | 0.185 |
| Salience Ventral Attention A: Frontal Medial 1 | 3.4 | 0.004 | 0.335 | 1.0 | 0.319 | 0.018 | 0.9 | 0.360 | 0.016 | 1.1 | 0.294 | 0.020 | 0.1 | 0.736 | 0.002 | 0.9 | 0.360 | 0.016 | 0.2 | 0.658 | 0.004 | 12.5 | 0.001 | 0.188 |
| Salience Ventral Attention B: Lateral Prefrontal Cort | 5.7 | 0.000 | 0.456 | 3.0 | 0.091 | 0.052 | 0.9 | 0.339 | 0.017 | 0.2 | 0.649 | 0.004 | 0.5 | 0.493 | 0.009 | 0.7 | 0.405 | 0.013 | 0.5 | 0.495 | 0.009 | 20.7 | 0.000 | 0.277 |
| Salience Ventral Attention B: Medial Posterior Prefro | 5.9 | 0.000 | 0.468 | 6.2 | 0.016 | 0.103 | 0.0 | 0.870 | 0.000 | 0.0 | 0.961 | 0.000 | 0.1 | 0.711 | 0.003 | 2.0 | 0.160 | 0.036 | 0.2 | 0.662 | 0.004 | 16.3 | 0.000 | 0.232 |
| LimbiC: B: Orbital Frontal Cortex 1 | 5.1 | 0.000 | 0.429 | 5.0 | 0.029 | 0.085 | 2.8 | 0.102 | 0.049 | 0.2 | 0.632 | 0.004 | 0.0 | 0.904 | 0.000 | 3.7 | 0.059 | 0.064 | 0.1 | 0.769 | 0.002 | 1.1 | 0.296 | 0.020 |
| LimbiC: A: Temporal Pole 1 | 9.2 | 0.000 | 0.578 | 7.2 | 0.010 | 0.117 | 1.0 | 0.328 | 0.018 | 0.2 | 0.679 | 0.003 | 0.2 | 0.690 | 0.003 | 0.7 | 0.413 | 0.012 | 0.2 | 0.685 | 0.003 | 19.4 | 0.000 | 0.264 |
| LimbiC: A: Temporal Pole 2 | 6.5 | 0.000 | 0.489 | 11.2 | 0.002 | 0.172 | 0.0 | 0.853 | 0.001 | 1.7 | 0.197 | 0.031 | 0.0 | 0.970 | 0.000 | 2.6 | 0.113 | 0.046 | 1.6 | 0.215 | 0.028 | 10.4 | 0.002 | 0.161 |
| Control A: Intraparietal Sulcus 1 | 6.2 | 0.000 | 0.480 | 4.1 | 0.048 | 0.070 | 0.1 | 0.749 | 0.002 | 0.3 | 0.591 | 0.005 | 0.3 | 0.571 | 0.006 | 1.4 | 0.240 | 0.025 | 0.4 | 0.538 | 0.007 | 21.5 | 0.000 | 0.285 |
| Control A: Lateral Prefrontal Cortex 1 | 6.4 | 0.000 | 0.485 | 0.6 | 0.432 | 0.011 | 1.7 | 0.196 | 0.031 | 0.4 | 0.519 | 0.008 | 0.1 | 0.781 | 0.001 | 3.9 | 0.052 | 0.068 | 0.1 | 0.745 | 0.002 | 19.0 | 0.000 | 0.260 |
| Control A: Lateral Prefrontal Cortex 2 | 6.1 | 0.000 | 0.475 | 1.7 | 0.204 | 0.030 | 1.7 | 0.193 | 0.031 | 0.2 | 0.645 | 0.004 | 0.0 | 0.874 | 0.000 | 3.1 | 0.084 | 0.054 | 0.9 | 0.335 | 0.017 | 16.5 | 0.000 | 0.234 |
| Control B: Lateral Prefrontal Cortex 1 | 7.0 | 0.000 | 0.508 | 3.2 | 0.081 | 0.055 | 6.5 | 0.014 | 0.107 | 1.1 | 0.303 | 0.020 | 0.2 | 0.681 | 0.003 | 2.1 | 0.151 | 0.038 | 0.3 | 0.569 | 0.006 | 8.5 | 0.005 | 0.136 |
| Control C: Precuneus 1 | 5.3 | 0.000 | 0.442 | 2.3 | 0.135 | 0.041 | 0.7 | 0.413 | 0.012 | 0.0 | 0.878 | 0.000 | 0.4 | 0.515 | 0.008 | 3.3 | 0.073 | 0.058 | 0.7 | 0.404 | 0.013 | 18.4 | 0.000 | 0.254 |
| Control C: Precuneus 2 | 4.6 | 0.000 | 0.404 | 2.6 | 0.115 | 0.045 | 0.2 | 0.649 | 0.004 | 0.2 | 0.660 | 0.004 | 0.8 | 0.367 | 0.015 | 4.8 | 0.033 | 0.082 | 0.3 | 0.565 | 0.006 | 16.8 | 0.000 | 0.237 |
| Control C: Cingulate Posterior 1 | 6.3 | 0.000 | 0.481 | 0.9 | 0.342 | 0.017 | 0.7 | 0.418 | 0.012 | 0.0 | 0.983 | 0.000 | 0.0 | 0.981 | 0.000 | 5.3 | 0.025 | 0.089 | 0.1 | 0.738 | 0.002 | 18.2 | 0.000 | 0.252 |
| Default A: Dorsal Prefrontal Cortex 1 | 4.7 | 0.000 | 0.408 | 3.6 | 0.065 | 0.062 | 0.1 | 0.726 | 0.002 | 0.0 | 0.840 | 0.001 | 0.5 | 0.475 | 0.010 | 0.3 | 0.604 | 0.005 | 0.5 | 0.475 | 0.009 | 15.0 | 0.000 | 0.217 |
| Default A: Precuneus Posterior Cingulate Cortex1 | 4.3 | 0.001 | 0.391 | 0.3 | 0.589 | 0.005 | 0.7 | 0.417 | 0.012 | 0.1 | 0.704 | 0.003 | 0.4 | 0.513 | 0.008 | 6.8 | 0.012 | 0.112 | 0.0 | 0.835 | 0.001 | 14.1 | 0.000 | 0.207 |
| Default A: Medial Prefrontal Cortex 1 | 5.3 | 0.000 | 0.438 | 7.5 | 0.008 | 0.122 | 0.5 | 0.483 | 0.009 | 0.4 | 0.512 | 0.008 | 0.5 | 0.476 | 0.009 | 3.2 | 0.080 | 0.056 | 0.0 | 0.846 | 0.001 | 8.6 | 0.005 | 0.138 |
| Default B: Temp 1 | 5.7 | 0.000 | 0.456 | 1.2 | 0.274 | 0.022 | 0.0 | 0.989 | 0.000 | 0.3 | 0.578 | 0.006 | 0.1 | 0.802 | 0.001 | 1.5 | 0.223 | 0.027 | 0.8 | 0.368 | 0.015 | 16.6 | 0.000 | 0.235 |
| Default B: Temp 2 | 3.3 | 0.004 | 0.329 | 2.3 | 0.132 | 0.041 | 0.0 | 0.877 | 0.000 | 0.0 | 0.995 | 0.000 | 0.7 | 0.403 | 0.013 | 4.1 | 0.049 | 0.070 | 1.0 | 0.319 | 0.018 | 9.7 | 0.003 | 0.153 |
| Default B: Inferior Parietal Lobule 1 | 6.0 | 0.000 | 0.471 | 1.0 | 0.311 | 0.019 | 0.8 | 0.368 | 0.015 | 0.2 | 0.678 | 0.003 | 0.0 | 0.958 | 0.000 | 3.8 | 0.056 | 0.066 | 1.3 | 0.260 | 0.023 | 16.6 | 0.000 | 0.235 |
| Default B: Dorsal Prefrontal Cortex 1 | 4.6 | 0.000 | 0.405 | 4.1 | 0.048 | 0.070 | 0.1 | 0.716 | 0.002 | 0.1 | 0.709 | 0.003 | 0.3 | 0.599 | 0.005 | 0.7 | 0.413 | 0.012 | 2.0 | 0.159 | 0.036 | 9.8 | 0.003 | 0.154 |
| Default B: Lateral Prefrontal Cortex 1 | 3.6 | 0.002 | 0.351 | 1.9 | 0.176 | 0.034 | 0.5 | 0.486 | 0.009 | 0.0 | 0.958 | 0.000 | 0.1 | 0.769 | 0.002 | 1.1 | 0.293 | 0.020 | 1.8 | 0.181 | 0.033 | 11.2 | 0.001 | 0.172 |
| Default B: Ventral Prefrontal Cortex 1 | 5.0 | 0.000 | 0.427 | 5.6 | 0.021 | 0.094 | 0.1 | 0.732 | 0.002 | 0.2 | 0.689 | 0.003 | 0.2 | 0.654 | 0.004 | 1.7 | 0.202 | 0.030 | 0.9 | 0.339 | 0.017 | 16.7 | 0.000 | 0.236 |
| Default B: Ventral Prefrontal Cortex 2 | 8.5 | 0.000 | 0.559 | 3.6 | 0.064 | 0.062 | 3.1 | 0.083 | 0.055 | 0.6 | 0.430 | 0.012 | 0.0 | 0.996 | 0.000 | 5.8 | 0.020 | 0.096 | 0.7 | 0.401 | 0.013 | 11.9 | 0.001 | 0.181 |
| Default C: Retro Superior Parietal Lobuleenial 1 | 5.1 | 0.000 | 0.432 | 1.3 | 0.264 | 0.023 | 0.2 | 0.655 | 0.004 | 0.1 | 0.818 | 0.001 | 1.9 | 0.177 | 0.033 | 2.9 | 0.093 | 0.051 | 1.6 | 0.205 | 0.030 | 16.8 | 0.000 | 0.237 |
| Default C: Parahippocampal Cortex 1 | 3.7 | 0.002 | 0.353 | 3.9 | 0.055 | 0.067 | 0.0 | 0.887 | 0.000 | 0.0 | 0.827 | 0.001 | 0.4 | 0.528 | 0.007 | 3.2 | 0.079 | 0.056 | 2.9 | 0.092 | 0.052 | 7.8 | 0.007 | 0.126 |
| Temporal Parietal 1 | 2.9 | 0.011 | 0.299 | 0.9 | 0.352 | 0.016 | 0.0 | 0.852 | 0.001 | 0.0 | 0.901 | 0.000 | 0.1 | 0.766 | 0.002 | 5.2 | 0.027 | 0.087 | 1.7 | 0.203 | 0.030 | 8.3 | 0.006 | 0.133 |

Continued …

*… Table S4 continued*

***Right Hemisphere***

| | Overall Model | | | Age Category | | | Systolic BP | | | Diastolic BP | | | Resting HR | | | BMI | | | Years of Education | | | Sex | | |
|---|---|---|---|---|---|---|---|---|---|---|---|---|---|---|---|---|---|---|---|---|---|---|---|---|
| | F | p-FDR | $\eta^2_p$ | F | p | $\eta^2_p$ | F | p | $\eta^2_p$ | F | p | $\eta^2_p$ | F | p | $\eta^2_p$ | F | p | $\eta^2_p$ | F | p | $\eta^2_p$ | F | p | $\eta^2_p$ |
| Visual Central: Extra Striate Cortex 1 | 2.6 | 0.019 | 0.277 | 0.1 | 0.734 | 0.002 | 0.0 | 0.880 | 0.000 | 0.0 | 0.836 | 0.001 | 0.0 | 0.899 | 0.000 | 2.2 | 0.144 | 0.039 | 1.0 | 0.317 | 0.019 | 11.1 | 0.002 | 0.170 |
| Visual Central: Extra Striate Cortex 2 | 3.8 | 0.002 | 0.358 | 4.0 | 0.050 | 0.069 | 0.0 | 0.880 | 0.000 | 1.1 | 0.301 | 0.020 | 0.3 | 0.582 | 0.006 | 1.5 | 0.230 | 0.027 | 0.3 | 0.560 | 0.006 | 7.9 | 0.007 | 0.127 |
| Visual Central: Extra Striate Cortex 3 | 4.6 | 0.000 | 0.406 | 0.9 | 0.357 | 0.016 | 0.5 | 0.464 | 0.010 | 0.0 | 0.914 | 0.000 | 0.0 | 0.897 | 0.000 | 2.5 | 0.121 | 0.044 | 0.1 | 0.802 | 0.001 | 16.2 | 0.000 | 0.230 |
| Visual Peripheral: Striate Cortex Calcarine 1 | 2.3 | 0.035 | 0.253 | 0.9 | 0.354 | 0.016 | 0.0 | 0.900 | 0.000 | 0.0 | 0.940 | 0.000 | 1.2 | 0.282 | 0.021 | 1.7 | 0.193 | 0.031 | 0.4 | 0.517 | 0.008 | 9.8 | 0.003 | 0.153 |
| Visual Peripheral: Extra Striate Inferior 1 | 2.0 | 0.062 | 0.230 | 0.1 | 0.743 | 0.002 | 0.0 | 0.880 | 0.000 | 0.0 | 0.878 | 0.000 | 1.0 | 0.312 | 0.019 | 1.7 | 0.201 | 0.030 | 1.1 | 0.308 | 0.019 | 9.4 | 0.003 | 0.148 |
| Visual Peripheral: Extra Striate Superior 1 | 3.3 | 0.005 | 0.326 | 2.1 | 0.158 | 0.037 | 0.1 | 0.782 | 0.001 | 0.0 | 0.885 | 0.000 | 1.1 | 0.306 | 0.019 | 4.4 | 0.040 | 0.076 | 1.7 | 0.203 | 0.030 | 9.9 | 0.003 | 0.155 |
| Somatomotor A: 1 | 3.1 | 0.006 | 0.317 | 0.2 | 0.647 | 0.004 | 0.0 | 0.957 | 0.000 | 0.0 | 0.865 | 0.001 | 0.1 | 0.717 | 0.002 | 4.2 | 0.045 | 0.072 | 1.6 | 0.213 | 0.029 | 7.9 | 0.007 | 0.128 |
| Somatomotor A: 2 | 3.6 | 0.003 | 0.348 | 0.6 | 0.444 | 0.011 | 0.0 | 0.996 | 0.000 | 0.0 | 0.832 | 0.001 | 0.4 | 0.533 | 0.007 | 1.3 | 0.258 | 0.024 | 0.0 | 0.873 | 0.000 | 15.9 | 0.000 | 0.227 |
| Somatomotor A: 3 | 4.7 | 0.000 | 0.410 | 1.0 | 0.329 | 0.018 | 1.6 | 0.210 | 0.029 | 0.6 | 0.439 | 0.011 | 1.7 | 0.199 | 0.030 | 2.3 | 0.134 | 0.041 | 0.2 | 0.691 | 0.003 | 16.1 | 0.000 | 0.230 |
| Somatomotor A: 4 | 3.6 | 0.003 | 0.347 | 0.6 | 0.453 | 0.010 | 0.0 | 0.963 | 0.000 | 0.2 | 0.694 | 0.003 | 0.6 | 0.455 | 0.010 | 0.1 | 0.799 | 0.001 | 0.4 | 0.530 | 0.007 | 16.2 | 0.000 | 0.231 |
| Somatomotor B: Auditory 1 | 4.1 | 0.001 | 0.380 | 0.1 | 0.809 | 0.001 | 0.7 | 0.414 | 0.012 | 0.4 | 0.513 | 0.008 | 0.3 | 0.566 | 0.006 | 6.3 | 0.015 | 0.105 | 0.1 | 0.769 | 0.002 | 12.6 | 0.001 | 0.189 |
| Somatomotor B: S2 1 | 4.0 | 0.001 | 0.375 | 0.0 | 0.870 | 0.000 | 1.1 | 0.302 | 0.020 | 0.5 | 0.496 | 0.009 | 0.1 | 0.785 | 0.001 | 8.1 | 0.006 | 0.131 | 0.2 | 0.639 | 0.004 | 10.0 | 0.003 | 0.156 |
| Somatomotor B: S2 2 | 4.7 | 0.000 | 0.408 | 0.0 | 0.899 | 0.000 | 1.4 | 0.246 | 0.025 | 0.4 | 0.519 | 0.008 | 0.0 | 0.927 | 0.000 | 6.4 | 0.014 | 0.106 | 0.0 | 0.870 | 0.001 | 8.8 | 0.004 | 0.140 |
| Somatomotor B: Central 1 | 3.6 | 0.002 | 0.349 | 0.1 | 0.819 | 0.001 | 0.9 | 0.356 | 0.016 | 0.1 | 0.728 | 0.002 | 0.1 | 0.802 | 0.001 | 4.8 | 0.032 | 0.082 | 0.1 | 0.783 | 0.001 | 13.0 | 0.001 | 0.194 |
| Dorsal Attention A: Temporal Occipital 1 | 5.3 | 0.000 | 0.440 | 1.9 | 0.179 | 0.033 | 0.0 | 0.888 | 0.000 | 1.9 | 0.171 | 0.034 | 0.1 | 0.810 | 0.001 | 1.7 | 0.197 | 0.031 | 0.0 | 0.893 | 0.000 | 17.8 | 0.000 | 0.248 |
| Dorsal Attention A: Parietal Occipital 1 | 4.1 | 0.001 | 0.379 | 0.5 | 0.493 | 0.009 | 0.2 | 0.659 | 0.004 | 0.4 | 0.541 | 0.007 | 0.3 | 0.583 | 0.006 | 3.9 | 0.052 | 0.068 | 0.4 | 0.542 | 0.007 | 11.9 | 0.001 | 0.181 |
| Dorsal Attention A: Superior Parietal Lobule 1 | 5.5 | 0.000 | 0.450 | 4.5 | 0.038 | 0.077 | 0.0 | 0.950 | 0.000 | 0.3 | 0.604 | 0.005 | 0.2 | 0.678 | 0.003 | 2.1 | 0.157 | 0.037 | 0.2 | 0.697 | 0.003 | 22.3 | 0.000 | 0.292 |
| Dorsal Attention B: Post Central 1 | 4.2 | 0.001 | 0.385 | 0.9 | 0.356 | 0.016 | 2.3 | 0.133 | 0.041 | 0.3 | 0.568 | 0.006 | 0.3 | 0.594 | 0.005 | 6.4 | 0.014 | 0.106 | 0.0 | 0.996 | 0.000 | 11.6 | 0.001 | 0.176 |
| Dorsal Attention B: Post Central 2 | 4.2 | 0.001 | 0.385 | 3.9 | 0.053 | 0.067 | 0.0 | 0.864 | 0.001 | 0.0 | 0.998 | 0.000 | 1.4 | 0.240 | 0.025 | 0.7 | 0.397 | 0.013 | 0.5 | 0.465 | 0.010 | 16.5 | 0.000 | 0.234 |
| Dorsal Attention B: Frontal Eye Fields 1 | 4.8 | 0.000 | 0.418 | 4.2 | 0.045 | 0.072 | 0.1 | 0.811 | 0.001 | 0.0 | 0.983 | 0.000 | 0.2 | 0.678 | 0.003 | 0.7 | 0.413 | 0.012 | 0.2 | 0.668 | 0.003 | 17.5 | 0.000 | 0.245 |
| Salience Ventral Attention A: Parietal Operculum 1 | 4.4 | 0.001 | 0.395 | 0.0 | 0.980 | 0.000 | 1.3 | 0.260 | 0.023 | 0.0 | 0.857 | 0.001 | 0.1 | 0.727 | 0.002 | 7.6 | 0.008 | 0.123 | 0.1 | 0.761 | 0.002 | 5.7 | 0.020 | 0.096 |
| Salience Ventral Attention A: Insula: 1 | 5.5 | 0.000 | 0.448 | 0.0 | 0.919 | 0.000 | 0.9 | 0.344 | 0.017 | 0.5 | 0.478 | 0.009 | 0.3 | 0.617 | 0.005 | 6.6 | 0.013 | 0.109 | 0.0 | 0.887 | 0.000 | 12.9 | 0.001 | 0.193 |
| Salience Ventral Attention A: Parietal Medial 1 | 3.4 | 0.004 | 0.337 | 0.8 | 0.363 | 0.015 | 1.3 | 0.251 | 0.024 | 0.1 | 0.737 | 0.002 | 0.4 | 0.550 | 0.007 | 2.4 | 0.128 | 0.042 | 0.0 | 0.845 | 0.001 | 9.6 | 0.003 | 0.151 |
| Salience Ventral Attention A: Frontal Medial 1 | 5.4 | 0.000 | 0.445 | 0.8 | 0.385 | 0.014 | 0.0 | 0.893 | 0.000 | 0.5 | 0.490 | 0.009 | 0.0 | 0.867 | 0.001 | 0.8 | 0.381 | 0.014 | 0.1 | 0.769 | 0.002 | 20.2 | 0.000 | 0.272 |
| Salience Ventral Attention B: Inferior Parietal Lobule | 4.0 | 0.001 | 0.372 | 1.9 | 0.173 | 0.034 | 0.3 | 0.581 | 0.006 | 0.1 | 0.728 | 0.002 | 0.0 | 0.899 | 0.000 | 3.4 | 0.071 | 0.059 | 0.0 | 0.972 | 0.000 | 6.3 | 0.015 | 0.104 |
| Salience Ventral Attention B: Lateral Prefrontal Cort | 6.6 | 0.000 | 0.493 | 0.5 | 0.471 | 0.010 | 1.0 | 0.325 | 0.018 | 0.1 | 0.795 | 0.001 | 1.2 | 0.281 | 0.022 | 3.3 | 0.076 | 0.057 | 0.0 | 0.838 | 0.001 | 17.5 | 0.000 | 0.245 |
| Salience Ventral Attention B: Medial Posterior Prefro | 5.5 | 0.000 | 0.449 | 4.8 | 0.032 | 0.082 | 0.1 | 0.723 | 0.002 | 0.0 | 0.962 | 0.000 | 0.2 | 0.635 | 0.004 | 2.1 | 0.154 | 0.037 | 0.8 | 0.371 | 0.015 | 16.8 | 0.000 | 0.237 |
| LimbiC: B: Orbital Frontal Cortex 1 | 6.2 | 0.000 | 0.477 | 2.7 | 0.109 | 0.047 | 6.4 | 0.014 | 0.106 | 1.7 | 0.193 | 0.031 | 0.0 | 0.843 | 0.001 | 4.8 | 0.033 | 0.081 | 1.4 | 0.238 | 0.026 | 1.9 | 0.172 | 0.034 |
| LimbiC: A: Temporal Pole 1 | 9.1 | 0.000 | 0.573 | 7.9 | 0.007 | 0.128 | 0.4 | 0.511 | 0.008 | 0.5 | 0.470 | 0.010 | 0.1 | 0.801 | 0.001 | 1.1 | 0.304 | 0.020 | 1.7 | 0.202 | 0.030 | 13.3 | 0.001 | 0.197 |
| Control A: Intraparietal Sulcus 1 | 5.3 | 0.000 | 0.439 | 2.3 | 0.136 | 0.041 | 0.4 | 0.535 | 0.007 | 0.1 | 0.728 | 0.002 | 0.1 | 0.738 | 0.002 | 2.7 | 0.107 | 0.047 | 0.1 | 0.703 | 0.003 | 17.3 | 0.000 | 0.243 |
| Control A: Lateral Prefrontal Cortex 1 | 5.3 | 0.000 | 0.438 | 1.8 | 0.186 | 0.032 | 1.8 | 0.185 | 0.032 | 0.3 | 0.563 | 0.006 | 0.5 | 0.471 | 0.010 | 3.9 | 0.054 | 0.067 | 0.0 | 0.990 | 0.000 | 11.7 | 0.001 | 0.178 |
| Control A: Lateral Prefrontal Cortex 2 | 6.7 | 0.000 | 0.500 | 1.3 | 0.266 | 0.023 | 0.2 | 0.689 | 0.003 | 0.1 | 0.751 | 0.002 | 0.1 | 0.749 | 0.002 | 5.7 | 0.021 | 0.095 | 0.0 | 0.862 | 0.001 | 10.3 | 0.002 | 0.160 |
| Control B: Temporal 1 | 7.8 | 0.000 | 0.535 | 2.7 | 0.107 | 0.048 | 0.9 | 0.336 | 0.017 | 0.2 | 0.675 | 0.003 | 0.1 | 0.712 | 0.003 | 3.4 | 0.071 | 0.059 | 0.1 | 0.755 | 0.002 | 19.4 | 0.000 | 0.264 |
| Control B: inferior parietal lobule 1 | 4.1 | 0.001 | 0.377 | 1.4 | 0.249 | 0.024 | 0.1 | 0.751 | 0.002 | 0.4 | 0.536 | 0.007 | 0.0 | 0.873 | 0.000 | 2.9 | 0.097 | 0.050 | 0.0 | 0.831 | 0.001 | 12.6 | 0.001 | 0.190 |
| Control B: Lateral Prefrontal Cortexd 1 | 4.8 | 0.000 | 0.415 | 2.9 | 0.093 | 0.052 | 0.0 | 0.838 | 0.001 | 0.0 | 0.863 | 0.001 | 0.8 | 0.367 | 0.015 | 2.6 | 0.113 | 0.046 | 0.0 | 0.869 | 0.001 | 13.0 | 0.001 | 0.194 |
| Control B: Lateral Prefrontal Cortexv 1 | 7.7 | 0.000 | 0.532 | 0.7 | 0.401 | 0.013 | 6.4 | 0.014 | 0.107 | 0.5 | 0.497 | 0.009 | 0.0 | 0.964 | 0.000 | 3.2 | 0.078 | 0.056 | 1.3 | 0.260 | 0.023 | 13.3 | 0.001 | 0.198 |
| Control C: Cingulate Posterior 1 | 5.0 | 0.000 | 0.425 | 0.8 | 0.386 | 0.014 | 0.1 | 0.800 | 0.001 | 0.1 | 0.819 | 0.001 | 0.1 | 0.766 | 0.002 | 3.9 | 0.052 | 0.068 | 0.2 | 0.647 | 0.004 | 17.1 | 0.000 | 0.240 |
| Control C: Precuneus 1 | 3.1 | 0.007 | 0.314 | 1.5 | 0.225 | 0.027 | 0.1 | 0.707 | 0.003 | 0.0 | 0.984 | 0.000 | 0.8 | 0.368 | 0.015 | 3.8 | 0.057 | 0.066 | 1.2 | 0.278 | 0.022 | 8.9 | 0.004 | 0.141 |
| Default A: Inferior Parietal Lobule 1 | 5.0 | 0.000 | 0.427 | 0.2 | 0.688 | 0.003 | 0.5 | 0.470 | 0.010 | 0.0 | 0.889 | 0.000 | 0.3 | 0.614 | 0.005 | 6.4 | 0.014 | 0.106 | 0.0 | 0.933 | 0.000 | 13.3 | 0.001 | 0.198 |
| Default A: Dorsal Prefrontal Cortex 1 | 5.5 | 0.000 | 0.447 | 6.2 | 0.016 | 0.103 | 0.1 | 0.823 | 0.001 | 0.1 | 0.820 | 0.001 | 0.7 | 0.391 | 0.014 | 1.3 | 0.255 | 0.024 | 0.2 | 0.647 | 0.004 | 17.1 | 0.000 | 0.241 |
| Default A: Precuneus Posterior Cingulate Cortex 1 | 2.8 | 0.012 | 0.294 | 0.1 | 0.730 | 0.002 | 0.2 | 0.677 | 0.003 | 0.0 | 0.890 | 0.000 | 0.6 | 0.457 | 0.010 | 5.7 | 0.021 | 0.095 | 0.1 | 0.726 | 0.002 | 7.6 | 0.008 | 0.124 |
| Default A: Medial Prefrontal Cortex 1 | 5.6 | 0.000 | 0.454 | 1.2 | 0.281 | 0.022 | 2.0 | 0.165 | 0.035 | 1.2 | 0.272 | 0.022 | 0.3 | 0.557 | 0.006 | 2.8 | 0.101 | 0.049 | 0.0 | 0.832 | 0.001 | 14.0 | 0.000 | 0.206 |
| Default B: Dorsal Prefrontal Cortex 1 | 5.6 | 0.000 | 0.452 | 3.5 | 0.068 | 0.060 | 0.1 | 0.728 | 0.002 | 0.0 | 0.897 | 0.000 | 0.8 | 0.381 | 0.014 | 1.1 | 0.298 | 0.020 | 0.7 | 0.417 | 0.012 | 11.9 | 0.001 | 0.181 |
| Default B: Ventral Prefrontal Cortex 1 | 4.8 | 0.000 | 0.413 | 3.9 | 0.052 | 0.068 | 1.4 | 0.241 | 0.025 | 0.2 | 0.638 | 0.004 | 0.1 | 0.711 | 0.003 | 2.8 | 0.100 | 0.049 | 1.2 | 0.285 | 0.021 | 4.2 | 0.046 | 0.072 |
| Default B: Ventral Prefrontal Cortex 2 | 5.0 | 0.000 | 0.426 | 1.2 | 0.273 | 0.022 | 1.1 | 0.308 | 0.019 | 0.1 | 0.716 | 0.002 | 0.1 | 0.799 | 0.001 | 4.1 | 0.048 | 0.070 | 0.5 | 0.482 | 0.009 | 4.6 | 0.037 | 0.078 |
| Default C: Retro Superior Parietal Lobuleenial 1 | 2.5 | 0.021 | 0.273 | 1.0 | 0.315 | 0.019 | 0.1 | 0.801 | 0.001 | 0.0 | 0.834 | 0.001 | 0.7 | 0.394 | 0.013 | 3.1 | 0.084 | 0.054 | 0.4 | 0.524 | 0.008 | 7.4 | 0.009 | 0.121 |
| Default C: Parahippocampal Cortex 1 | 2.8 | 0.012 | 0.292 | 2.8 | 0.098 | 0.050 | 0.0 | 0.968 | 0.000 | 0.1 | 0.757 | 0.002 | 0.0 | 0.918 | 0.000 | 1.6 | 0.207 | 0.029 | 0.2 | 0.664 | 0.004 | 6.6 | 0.013 | 0.108 |
| Temporal Parietal 1 | 4.8 | 0.000 | 0.416 | 5.9 | 0.019 | 0.098 | 0.0 | 0.874 | 0.000 | 0.1 | 0.701 | 0.003 | 0.1 | 0.724 | 0.002 | 1.3 | 0.257 | 0.024 | 0.2 | 0.622 | 0.005 | 14.1 | 0.000 | 0.207 |
| Temporal Parietal 2 | 5.4 | 0.000 | 0.445 | 0.3 | 0.605 | 0.005 | 0.9 | 0.345 | 0.017 | 0.1 | 0.727 | 0.002 | 0.0 | 0.923 | 0.000 | 5.6 | 0.021 | 0.094 | 0.1 | 0.803 | 0.001 | 16.2 | 0.000 | 0.231 |
| Temporal Parietal 3 | 3.7 | 0.002 | 0.353 | 0.3 | 0.607 | 0.005 | 0.2 | 0.622 | 0.005 | 0.1 | 0.775 | 0.002 | 0.0 | 0.852 | 0.001 | 4.3 | 0.044 | 0.073 | 0.0 | 0.983 | 0.000 | 12.8 | 0.001 | 0.191 |

Table S5. General linear models of the association between cerebral blood flow among the four sub-groups based on age category and HOMA-IR levels, with blood pressure, cortical thickness, resting heart rate, BMI, sex, years of education as covariates. Post-hoc contrasts in the regions with significant group difference are shown, comparing the young insulin sensitive group to the other three groups.

***Left Hemisphere***

| | Overall | | | Systolic BP | | | Diastolic BP | | | Resting HR | | | BMI | | | Years of Education | | | Sex | | | Cortical Thickness | | | 4 Groups: Age Category and HOMA Median Split | | | Post-Hoc Contrasts | | |
|---|---|---|---|---|---|---|---|---|---|---|---|---|---|---|---|---|---|---|---|---|---|---|---|---|---|---|---|---|---|---|
| | F | p-FDR | $\eta^2_p$ | F | p | $\eta^2_p$ | F | p | $\eta^2_p$ | F | p | $\eta^2_p$ | F | p | $\eta^2_p$ | F | p | $\eta^2_p$ | F | p | $\eta^2_p$ | F | p | $\eta^2_p$ | F | p | $\eta^2_p$ | YIS: vs YIR | YIS: vs OIS | YIS: vs OIR |
| Visual Central: Extra Striate Cortex 1 | 5.1 | 0.000 | 0.495 | 0.1 | 0.710 | 0.003 | 1.2 | 0.285 | 0.022 | 0.3 | 0.601 | 0.005 | 3.2 | 0.080 | 0.058 | 4.7 | 0.035 | 0.083 | 18.8 | 0.000 | 0.266 | 3.4 | 0.072 | 0.061 | 2.9 | 0.045 | 0.142 | 0.031 | 0.777 | 0.141 |
| Visual Central: Extra Striate Cortex 2 | 5.6 | 0.000 | 0.518 | 0.5 | 0.468 | 0.010 | 2.5 | 0.123 | 0.045 | 0.0 | 0.837 | 0.001 | 2.8 | 0.101 | 0.051 | 0.2 | 0.621 | 0.005 | 16.0 | 0.000 | 0.235 | 0.9 | 0.359 | 0.016 | 2.8 | 0.047 | 0.141 | 0.047 | 0.949 | 0.792 |
| Visual Central: Striate Cortex 1 | 5.2 | 0.000 | 0.498 | 1.4 | 0.236 | 0.027 | 0.2 | 0.644 | 0.004 | 0.6 | 0.443 | 0.011 | 4.9 | 0.031 | 0.087 | 0.1 | 0.821 | 0.001 | 23.2 | 0.000 | 0.308 | 0.0 | 0.961 | 0.000 | 2.4 | 0.077 | | | | |
| Visual Central: Extra Striate Cortex 3 | 4.2 | 0.001 | 0.449 | 2.1 | 0.157 | 0.038 | 0.0 | 0.899 | 0.000 | 0.3 | 0.609 | 0.005 | 3.6 | 0.063 | 0.065 | 0.1 | 0.740 | 0.002 | 13.8 | 0.000 | 0.210 | 0.0 | 0.876 | 0.000 | 1.1 | 0.362 | | | | |
| Visual Peripheral: Extra Striate Inferior 1 | 3.5 | 0.002 | 0.405 | 0.3 | 0.558 | 0.007 | 0.0 | 0.926 | 0.000 | 1.3 | 0.268 | 0.024 | 4.7 | 0.036 | 0.082 | 1.4 | 0.239 | 0.027 | 17.7 | 0.000 | 0.254 | 0.5 | 0.487 | 0.009 | 2.0 | 0.125 | | | | |
| Visual Peripheral: Striate Cortex Calcarine 1 | 3.6 | 0.002 | 0.406 | 0.7 | 0.397 | 0.014 | 0.0 | 0.979 | 0.000 | 2.1 | 0.151 | 0.039 | 5.8 | 0.019 | 0.101 | 0.7 | 0.397 | 0.014 | 19.3 | 0.000 | 0.271 | 4.1 | 0.048 | 0.073 | 1.3 | 0.297 | | | | |
| Visual Peripheral: Extra Striate Cortex Sup 1 | 4.0 | 0.001 | 0.435 | 2.0 | 0.161 | 0.037 | 0.3 | 0.574 | 0.006 | 1.9 | 0.171 | 0.036 | 6.1 | 0.017 | 0.104 | 0.0 | 0.857 | 0.001 | 15.6 | 0.000 | 0.231 | 0.7 | 0.395 | 0.014 | 0.8 | 0.490 | | | | |
| Somatomotor A: 1 | 2.8 | 0.009 | 0.347 | 0.0 | 0.882 | 0.000 | 0.4 | 0.532 | 0.008 | 0.4 | 0.523 | 0.008 | 0.0 | 0.865 | 0.001 | 0.0 | 0.845 | 0.001 | 13.3 | 0.001 | 0.204 | 1.8 | 0.185 | 0.034 | 0.8 | 0.521 | | | | |
| Somatomotor A: 2 | 3.4 | 0.003 | 0.392 | 0.0 | 0.948 | 0.000 | 0.0 | 0.926 | 0.000 | 1.2 | 0.273 | 0.023 | 1.0 | 0.317 | 0.019 | 0.0 | 0.896 | 0.000 | 18.0 | 0.000 | 0.257 | 1.2 | 0.269 | 0.023 | 0.7 | 0.548 | | | | |
| Somatomotor B: Auditory 1 | 1.9 | 0.074 | 0.263 | 0.2 | 0.651 | 0.004 | 0.2 | 0.645 | 0.004 | 0.7 | 0.403 | 0.013 | 2.1 | 0.152 | 0.039 | 1.9 | 0.170 | 0.036 | 5.9 | 0.019 | 0.101 | 0.1 | 0.777 | 0.002 | 0.3 | 0.839 | | | | |
| Somatomotor B: S2 1 | 3.2 | 0.004 | 0.381 | 0.4 | 0.537 | 0.007 | 0.0 | 0.936 | 0.000 | 0.1 | 0.707 | 0.003 | 3.1 | 0.085 | 0.056 | 0.7 | 0.424 | 0.012 | 10.6 | 0.002 | 0.170 | 0.7 | 0.398 | 0.014 | 0.4 | 0.757 | | | | |
| Somatomotor B: S2 2 | 2.2 | 0.032 | 0.300 | 0.7 | 0.412 | 0.013 | 0.2 | 0.651 | 0.004 | 0.4 | 0.526 | 0.008 | 3.0 | 0.091 | 0.054 | 0.8 | 0.371 | 0.015 | 7.3 | 0.009 | 0.123 | 0.1 | 0.786 | 0.001 | 0.5 | 0.717 | | | | |
| Somatomotor B: Central 1 | 4.0 | 0.001 | 0.435 | 2.2 | 0.144 | 0.041 | 0.0 | 0.909 | 0.000 | 1.3 | 0.263 | 0.024 | 2.8 | 0.099 | 0.052 | 0.5 | 0.468 | 0.010 | 16.7 | 0.000 | 0.243 | 0.8 | 0.374 | 0.015 | 0.8 | 0.478 | | | | |
| Dorsal Attention A: Temporal Occipital 1 | 5.2 | 0.000 | 0.501 | 0.4 | 0.555 | 0.007 | 0.2 | 0.638 | 0.004 | 1.2 | 0.285 | 0.022 | 8.0 | 0.007 | 0.133 | 2.0 | 0.163 | 0.037 | 16.9 | 0.000 | 0.245 | 0.4 | 0.523 | 0.008 | 2.3 | 0.088 | | | | |
| Dorsal Attention A: Parietal Occipital 1 | 2.5 | 0.016 | 0.327 | 0.8 | 0.363 | 0.016 | 0.0 | 0.898 | 0.000 | 0.5 | 0.491 | 0.009 | 3.3 | 0.075 | 0.060 | 1.0 | 0.316 | 0.019 | 7.7 | 0.008 | 0.129 | 0.1 | 0.701 | 0.003 | 0.6 | 0.611 | | | | |
| Dorsal Attention A: Superior Parietal Lobule 1 | 6.5 | 0.000 | 0.556 | 1.6 | 0.218 | 0.029 | 0.0 | 0.969 | 0.000 | 0.4 | 0.539 | 0.007 | 1.7 | 0.198 | 0.032 | 0.2 | 0.627 | 0.005 | 24.4 | 0.000 | 0.319 | 1.1 | 0.298 | 0.021 | 2.2 | 0.103 | | | | |
| Dorsal Attention B: Post Central 1 | 2.7 | 0.012 | 0.338 | 1.0 | 0.315 | 0.019 | 0.4 | 0.519 | 0.008 | 0.7 | 0.410 | 0.013 | 1.6 | 0.206 | 0.031 | 0.2 | 0.678 | 0.003 | 11.7 | 0.001 | 0.184 | 0.0 | 0.870 | 0.001 | 0.6 | 0.629 | | | | |
| Dorsal Attention B: Post Central 2 | 3.0 | 0.005 | 0.368 | 0.6 | 0.453 | 0.011 | 0.0 | 0.950 | 0.000 | 0.9 | 0.350 | 0.017 | 2.7 | 0.104 | 0.050 | 0.4 | 0.534 | 0.007 | 14.2 | 0.000 | 0.214 | 0.6 | 0.456 | 0.011 | 0.7 | 0.553 | | | | |
| Dorsal Attention B: Post Central 3 | 4.3 | 0.001 | 0.451 | 0.8 | 0.384 | 0.015 | 0.5 | 0.475 | 0.010 | 1.5 | 0.219 | 0.029 | 0.8 | 0.383 | 0.015 | 0.0 | 0.911 | 0.000 | 17.1 | 0.000 | 0.248 | 1.7 | 0.195 | 0.032 | 1.7 | 0.173 | | | | |
| Dorsal Attention B: Frontal Eye Fields 1 | 3.9 | 0.001 | 0.426 | 0.0 | 0.936 | 0.000 | 0.2 | 0.683 | 0.003 | 1.0 | 0.334 | 0.018 | 1.5 | 0.223 | 0.028 | 0.3 | 0.563 | 0.006 | 14.8 | 0.000 | 0.222 | 2.8 | 0.102 | 0.051 | 1.3 | 0.287 | | | | |
| Salience Ventral Attention A: Parietal Operculum 1 | 4.5 | 0.000 | 0.463 | 0.4 | 0.535 | 0.007 | 0.8 | 0.362 | 0.016 | 0.0 | 0.990 | 0.000 | 1.7 | 0.195 | 0.032 | 0.4 | 0.547 | 0.007 | 14.3 | 0.000 | 0.216 | 0.0 | 0.957 | 0.000 | 1.8 | 0.160 | | | | |
| Salience Ventral Attention A: Insula: 1 | 4.4 | 0.001 | 0.458 | 0.3 | 0.600 | 0.005 | 0.0 | 0.975 | 0.000 | 1.0 | 0.314 | 0.019 | 1.8 | 0.188 | 0.033 | 3.0 | 0.091 | 0.054 | 16.2 | 0.000 | 0.237 | 3.9 | 0.053 | 0.070 | 1.4 | 0.249 | | | | |
| Salience Ventral Attention A: Insula: 2 | 4.0 | 0.001 | 0.436 | 1.5 | 0.230 | 0.028 | 1.1 | 0.293 | 0.021 | 0.8 | 0.378 | 0.015 | 4.8 | 0.034 | 0.084 | 0.6 | 0.460 | 0.011 | 12.3 | 0.001 | 0.191 | 0.4 | 0.537 | 0.007 | 0.7 | 0.542 | | | | |
| Salience Ventral Attention A: Parietal Medial 1 | 5.0 | 0.000 | 0.490 | 2.3 | 0.134 | 0.043 | 0.2 | 0.630 | 0.005 | 0.6 | 0.448 | 0.011 | 5.3 | 0.026 | 0.092 | 0.3 | 0.565 | 0.006 | 12.8 | 0.001 | 0.198 | 0.5 | 0.502 | 0.009 | 2.1 | 0.115 | | | | |
| Salience Ventral Attention A: Frontal Medial 1 | 2.9 | 0.007 | 0.357 | 0.4 | 0.520 | 0.008 | 1.2 | 0.279 | 0.023 | 0.0 | 0.979 | 0.000 | 1.5 | 0.226 | 0.028 | 0.0 | 0.910 | 0.000 | 12.9 | 0.001 | 0.199 | 1.9 | 0.170 | 0.036 | 0.9 | 0.432 | | | | |
| Salience Ventral Attention B: Lateral Prefrontal Corte | 4.6 | 0.000 | 0.472 | 1.4 | 0.245 | 0.026 | 0.1 | 0.702 | 0.003 | 1.0 | 0.328 | 0.018 | 1.3 | 0.258 | 0.025 | 0.1 | 0.713 | 0.003 | 21.0 | 0.000 | 0.288 | 0.0 | 0.997 | 0.000 | 1.5 | 0.222 | | | | |
| Salience Ventral Attention B: Medial Posterior Prefro | 4.9 | 0.000 | 0.486 | 0.0 | 0.936 | 0.000 | 0.1 | 0.726 | 0.002 | 0.2 | 0.648 | 0.004 | 1.8 | 0.187 | 0.033 | 0.0 | 0.946 | 0.000 | 15.1 | 0.000 | 0.225 | 0.0 | 0.959 | 0.000 | 2.7 | 0.056 | | | | |
| LimbiC: B: Orbital Frontal Cortex 1 | 4.1 | 0.001 | 0.443 | 2.8 | 0.100 | 0.051 | 0.0 | 0.898 | 0.000 | 0.0 | 0.927 | 0.000 | 2.0 | 0.159 | 0.038 | 0.2 | 0.645 | 0.004 | 0.8 | 0.379 | 0.015 | 2.0 | 0.168 | 0.036 | 2.1 | 0.114 | | | | |
| LimbiC: A: Temporal Pole 1 | 7.1 | 0.000 | 0.578 | 0.9 | 0.345 | 0.017 | 0.2 | 0.660 | 0.004 | 0.1 | 0.734 | 0.002 | 0.5 | 0.497 | 0.009 | 0.2 | 0.679 | 0.003 | 17.9 | 0.000 | 0.256 | 4.5 | 0.038 | 0.080 | 2.3 | 0.087 | | | | |
| LimbiC: A: Temporal Pole 2 | 5.3 | 0.000 | 0.505 | 0.0 | 0.931 | 0.000 | 1.0 | 0.329 | 0.018 | 0.1 | 0.721 | 0.002 | 4.0 | 0.052 | 0.071 | 1.3 | 0.253 | 0.025 | 11.5 | 0.001 | 0.182 | 0.0 | 0.876 | 0.000 | 4.3 | 0.009 | 0.198 | 0.414 | 0.042 | 0.160 |
| Control A: Intraparietal Sulcus 1 | 5.0 | 0.000 | 0.489 | 0.2 | 0.633 | 0.004 | 0.6 | 0.460 | 0.011 | 0.4 | 0.535 | 0.007 | 1.3 | 0.267 | 0.024 | 0.1 | 0.732 | 0.002 | 20.1 | 0.000 | 0.279 | 0.1 | 0.790 | 0.001 | 1.7 | 0.186 | | | | |
| Control A: Lateral Prefrontal Cortex 1 | 5.2 | 0.000 | 0.501 | 2.4 | 0.126 | 0.045 | 0.2 | 0.650 | 0.004 | 0.3 | 0.616 | 0.005 | 4.2 | 0.046 | 0.074 | 0.0 | 0.968 | 0.000 | 17.8 | 0.000 | 0.255 | 0.6 | 0.436 | 0.012 | 0.8 | 0.520 | | | | |
| Control A: Lateral Prefrontal Cortex 2 | 5.0 | 0.000 | 0.492 | 2.4 | 0.129 | 0.044 | 0.1 | 0.768 | 0.002 | 0.2 | 0.659 | 0.004 | 3.7 | 0.061 | 0.066 | 0.4 | 0.555 | 0.007 | 16.5 | 0.000 | 0.241 | 0.1 | 0.783 | 0.001 | 1.2 | 0.335 | | | | |
| Control B: Lateral Prefrontal Cortex 1 | 6.7 | 0.000 | 0.562 | 8.6 | 0.005 | 0.141 | 0.1 | 0.735 | 0.002 | 0.0 | 0.841 | 0.001 | 0.8 | 0.380 | 0.015 | 1.1 | 0.296 | 0.021 | 7.5 | 0.009 | 0.125 | 0.8 | 0.366 | 0.016 | 3.3 | 0.028 | 0.159 | 0.528 | 0.951 | 0.071 |
| Control C: Precuneus 1 | 4.5 | 0.000 | 0.465 | 1.2 | 0.285 | 0.022 | 0.0 | 0.866 | 0.001 | 0.6 | 0.455 | 0.011 | 3.2 | 0.079 | 0.058 | 0.2 | 0.682 | 0.003 | 17.9 | 0.000 | 0.256 | 0.1 | 0.757 | 0.002 | 1.5 | 0.219 | | | | |
| Control C: Precuneus 2 | 3.6 | 0.002 | 0.407 | 0.3 | 0.616 | 0.005 | 0.1 | 0.769 | 0.002 | 0.7 | 0.418 | 0.013 | 3.6 | 0.064 | 0.064 | 0.2 | 0.667 | 0.004 | 15.1 | 0.000 | 0.225 | 0.9 | 0.347 | 0.017 | 0.9 | 0.444 | | | | |
| Control C: Cingulate Posterior 1 | 5.9 | 0.000 | 0.532 | 1.4 | 0.241 | 0.026 | 0.4 | 0.525 | 0.008 | 0.0 | 0.976 | 0.000 | 4.1 | 0.047 | 0.074 | 0.9 | 0.349 | 0.017 | 17.1 | 0.000 | 0.248 | 0.2 | 0.639 | 0.004 | 2.2 | 0.097 | | | | |
| Default A: Dorsal Prefrontal Cortex 1 | 3.9 | 0.001 | 0.431 | 0.4 | 0.547 | 0.007 | 0.0 | 0.877 | 0.000 | 1.1 | 0.289 | 0.022 | 0.9 | 0.359 | 0.016 | 0.1 | 0.711 | 0.003 | 15.6 | 0.000 | 0.231 | 0.7 | 0.395 | 0.014 | 1.9 | 0.146 | | | | |
| Default A: Precuneus Posterior Cingulate Cortex1 | 3.7 | 0.001 | 0.416 | 1.2 | 0.273 | 0.023 | 0.0 | 0.937 | 0.000 | 0.6 | 0.434 | 0.012 | 6.0 | 0.018 | 0.103 | 0.1 | 0.793 | 0.001 | 13.5 | 0.001 | 0.206 | 0.0 | 0.947 | 0.000 | 0.9 | 0.472 | | | | |
| Default A: Medial Prefrontal Cortex 1 | 4.8 | 0.000 | 0.481 | 0.9 | 0.352 | 0.017 | 0.0 | 0.942 | 0.000 | 0.3 | 0.619 | 0.005 | 1.7 | 0.201 | 0.031 | 0.0 | 0.828 | 0.001 | 7.1 | 0.010 | 0.120 | 0.3 | 0.558 | 0.007 | 4.0 | 0.012 | 0.189 | 0.627 | 0.350 | 0.014 |
| Default B: Temp 1 | 4.5 | 0.000 | 0.463 | 0.0 | 0.897 | 0.000 | 0.1 | 0.736 | 0.002 | 0.0 | 0.853 | 0.001 | 1.3 | 0.262 | 0.024 | 1.1 | 0.294 | 0.021 | 16.3 | 0.000 | 0.239 | 4.8 | 0.033 | 0.084 | 0.6 | 0.604 | | | | |
| Default B: Temp 2 | 2.6 | 0.015 | 0.330 | 0.0 | 0.900 | 0.000 | 0.0 | 0.992 | 0.000 | 0.6 | 0.456 | 0.011 | 3.1 | 0.083 | 0.057 | 0.9 | 0.334 | 0.018 | 9.0 | 0.004 | 0.147 | 0.0 | 0.852 | 0.001 | 0.8 | 0.525 | | | | |
| Default B: Inferior Parietal Lobule 1 | 5.0 | 0.000 | 0.491 | 1.3 | 0.253 | 0.025 | 0.5 | 0.480 | 0.010 | 0.0 | 0.927 | 0.000 | 3.0 | 0.089 | 0.054 | 0.5 | 0.500 | 0.009 | 15.4 | 0.000 | 0.229 | 0.0 | 0.870 | 0.001 | 1.0 | 0.389 | | | | |
| Default B: Dorsal Prefrontal Cortex 1 | 3.8 | 0.001 | 0.423 | 0.3 | 0.571 | 0.006 | 0.1 | 0.713 | 0.003 | 0.8 | 0.387 | 0.014 | 1.4 | 0.240 | 0.026 | 1.2 | 0.269 | 0.023 | 10.5 | 0.002 | 0.168 | 0.3 | 0.607 | 0.005 | 1.9 | 0.142 | | | | |
| Default B: Lateral Prefrontal Cortex 1 | 3.1 | 0.005 | 0.371 | 1.0 | 0.327 | 0.019 | 0.0 | 0.981 | 0.000 | 0.4 | 0.524 | 0.008 | 1.7 | 0.192 | 0.033 | 0.9 | 0.351 | 0.017 | 11.4 | 0.001 | 0.180 | 0.1 | 0.724 | 0.002 | 1.2 | 0.323 | | | | |
| Default B: Ventral Prefrontal Cortex 1 | 4.1 | 0.001 | 0.438 | 0.0 | 0.897 | 0.000 | 0.1 | 0.717 | 0.003 | 0.5 | 0.478 | 0.010 | 2.2 | 0.140 | 0.041 | 0.5 | 0.497 | 0.009 | 16.6 | 0.000 | 0.242 | 0.2 | 0.625 | 0.005 | 2.2 | 0.101 | | | | |
| Default B: Ventral Prefrontal Cortex 2 | 7.3 | 0.000 | 0.583 | 4.5 | 0.038 | 0.080 | 0.2 | 0.684 | 0.003 | 0.1 | 0.802 | 0.001 | 5.6 | 0.022 | 0.097 | 1.8 | 0.184 | 0.034 | 11.6 | 0.001 | 0.183 | 0.0 | 0.847 | 0.001 | 2.2 | 0.096 | | | | |
| Default C: Retro Superior Parietal Lobuleenial 1 | 4.4 | 0.001 | 0.457 | 0.5 | 0.470 | 0.010 | 0.1 | 0.782 | 0.001 | 3.0 | 0.089 | 0.054 | 4.3 | 0.043 | 0.076 | 0.7 | 0.401 | 0.014 | 17.7 | 0.000 | 0.254 | 1.8 | 0.183 | 0.034 | 1.2 | 0.305 | | | | |
| Default C: Parahippocampal Cortex 1 | 3.9 | 0.001 | 0.426 | 0.0 | 0.886 | 0.000 | 0.0 | 0.840 | 0.001 | 2.1 | 0.156 | 0.038 | 7.6 | 0.008 | 0.128 | 2.4 | 0.130 | 0.043 | 11.0 | 0.002 | 0.175 | 0.8 | 0.370 | 0.015 | 3.6 | 0.019 | 0.172 | 0.052 | 0.496 | 0.654 |
| Temporal Parietal 1 | 2.3 | 0.028 | 0.305 | 0.1 | 0.748 | 0.002 | 0.0 | 0.897 | 0.000 | 0.2 | 0.622 | 0.005 | 5.2 | 0.026 | 0.092 | 1.1 | 0.305 | 0.020 | 8.2 | 0.006 | 0.136 | 0.0 | 0.968 | 0.000 | 0.5 | 0.713 | | | | |

*Continued …*

*… Table S5 Continued*
***Right Hemisphere***

| | Overall | | | Systolic BP | | | Diastolic BP | | | Resting HR | | | BMI | | | Years of Education | | | Sex | | | Cortical Thickness | | | 4 Groups: Age Category and HOMA Median Split | | | Post-Hoc Contrasts | | |
|---|---|---|---|---|---|---|---|---|---|---|---|---|---|---|---|---|---|---|---|---|---|---|---|---|---|---|---|---|---|---|
| | F | p-FDR | $\eta^2_p$ | F | p | $\eta^2_p$ | F | p | $\eta^2_p$ | F | p | $\eta^2_p$ | F | p | $\eta^2_p$ | F | p | $\eta^2_p$ | F | p | $\eta^2_p$ | F | p | $\eta^2_p$ | F | p | $\eta^2_p$ | YIS: vs YIR | YIS: vs OIS | YIS: vs OIR |
| Visual Central: Extra Striate Cortex 1 | 2.3 | 0.029 | 0.304 | 0.0 | 0.937 | 0.000 | 0.0 | 0.935 | 0.000 | 0.3 | 0.560 | 0.007 | 3.6 | 0.062 | 0.065 | 0.6 | 0.452 | 0.011 | 11.9 | 0.001 | 0.187 | 1.8 | 0.182 | 0.034 | 0.7 | 0.545 | | | | |
| Visual Central: Extra Striate Cortex 2 | 3.6 | 0.002 | 0.409 | 0.1 | 0.788 | 0.001 | 2.0 | 0.167 | 0.036 | 0.1 | 0.817 | 0.001 | 1.9 | 0.169 | 0.036 | 0.0 | 0.988 | 0.000 | 8.0 | 0.007 | 0.134 | 0.7 | 0.393 | 0.014 | 2.9 | 0.044 | 0.143 | 0.080 | 0.793 | 0.470 |
| Visual Central: Extra Striate Cortex 3 | 4.2 | 0.001 | 0.444 | 1.1 | 0.301 | 0.021 | 0.3 | 0.614 | 0.005 | 0.0 | 0.962 | 0.000 | 2.4 | 0.124 | 0.045 | 0.1 | 0.791 | 0.001 | 16.0 | 0.000 | 0.235 | 0.1 | 0.781 | 0.002 | 1.5 | 0.232 | | | | |
| Visual Peripheral: Striate Cortex Calcarine 1 | 2.1 | 0.039 | 0.291 | 0.0 | 0.929 | 0.000 | 0.0 | 0.947 | 0.000 | 2.4 | 0.130 | 0.044 | 3.2 | 0.079 | 0.058 | 0.1 | 0.747 | 0.002 | 11.2 | 0.002 | 0.178 | 1.1 | 0.292 | 0.021 | 1.2 | 0.312 | | | | |
| Visual Peripheral: Extra Striate Inferior 1 | 1.7 | 0.108 | 0.245 | 0.1 | 0.727 | 0.002 | 0.0 | 0.904 | 0.000 | 1.5 | 0.220 | 0.029 | 2.2 | 0.144 | 0.041 | 0.5 | 0.471 | 0.010 | 9.5 | 0.003 | 0.155 | 0.1 | 0.775 | 0.002 | 0.4 | 0.761 | | | | |
| Visual Peripheral: Extra Striate Superior 1 | 3.0 | 0.005 | 0.369 | 0.5 | 0.500 | 0.009 | 0.0 | 0.905 | 0.000 | 1.8 | 0.191 | 0.033 | 5.0 | 0.029 | 0.088 | 0.5 | 0.462 | 0.010 | 10.0 | 0.003 | 0.162 | 1.2 | 0.286 | 0.022 | 1.9 | 0.143 | | | | |
| Somatomotor A: 1 | 2.6 | 0.014 | 0.333 | 0.0 | 0.904 | 0.000 | 0.2 | 0.669 | 0.004 | 0.1 | 0.742 | 0.002 | 3.2 | 0.081 | 0.057 | 0.8 | 0.368 | 0.016 | 7.1 | 0.010 | 0.120 | 4.5 | 0.038 | 0.080 | 0.5 | 0.690 | | | | |
| Somatomotor A: 2 | 2.8 | 0.008 | 0.353 | 0.0 | 0.943 | 0.000 | 0.1 | 0.709 | 0.003 | 0.3 | 0.592 | 0.006 | 0.9 | 0.351 | 0.017 | 0.1 | 0.770 | 0.002 | 14.4 | 0.000 | 0.217 | 2.6 | 0.111 | 0.048 | 0.3 | 0.799 | | | | |
| Somatomotor A: 3 | 3.9 | 0.001 | 0.427 | 2.1 | 0.149 | 0.040 | 0.5 | 0.500 | 0.009 | 2.5 | 0.122 | 0.045 | 3.2 | 0.081 | 0.057 | 0.4 | 0.520 | 0.008 | 16.5 | 0.000 | 0.241 | 0.2 | 0.669 | 0.004 | 0.8 | 0.475 | | | | |
| Somatomotor A: 4 | 3.0 | 0.005 | 0.369 | 0.1 | 0.795 | 0.001 | 0.3 | 0.593 | 0.006 | 1.0 | 0.317 | 0.019 | 0.3 | 0.594 | 0.005 | 0.1 | 0.778 | 0.002 | 16.5 | 0.000 | 0.241 | 1.9 | 0.177 | 0.035 | 0.8 | 0.500 | | | | |
| Somatomotor B: Auditory 1 | 3.3 | 0.003 | 0.390 | 0.9 | 0.334 | 0.018 | 0.2 | 0.646 | 0.004 | 0.5 | 0.501 | 0.009 | 5.7 | 0.021 | 0.099 | 0.0 | 0.975 | 0.000 | 12.0 | 0.001 | 0.188 | 3.3 | 0.076 | 0.059 | 0.3 | 0.826 | | | | |
| Somatomotor B: S2 1 | 3.2 | 0.004 | 0.379 | 1.2 | 0.274 | 0.023 | 0.4 | 0.543 | 0.007 | 0.2 | 0.679 | 0.003 | 7.5 | 0.009 | 0.126 | 0.1 | 0.769 | 0.002 | 9.6 | 0.003 | 0.156 | 2.0 | 0.167 | 0.036 | 0.1 | 0.945 | | | | |
| Somatomotor B: S2 2 | 3.6 | 0.002 | 0.411 | 1.5 | 0.228 | 0.028 | 0.3 | 0.573 | 0.006 | 0.0 | 0.854 | 0.001 | 5.6 | 0.021 | 0.098 | 0.1 | 0.788 | 0.001 | 8.4 | 0.005 | 0.139 | 5.1 | 0.028 | 0.089 | 0.1 | 0.975 | | | | |
| Somatomotor B: Central 1 | 2.9 | 0.007 | 0.360 | 1.1 | 0.303 | 0.020 | 0.0 | 0.942 | 0.000 | 0.1 | 0.793 | 0.001 | 3.9 | 0.054 | 0.069 | 0.3 | 0.616 | 0.005 | 12.0 | 0.001 | 0.188 | 1.0 | 0.327 | 0.018 | 0.3 | 0.805 | | | | |
| Dorsal Attention A: Temporal Occipital 1 | 4.4 | 0.001 | 0.456 | 0.0 | 0.902 | 0.000 | 1.6 | 0.211 | 0.030 | 0.0 | 0.860 | 0.001 | 2.7 | 0.104 | 0.050 | 0.0 | 0.903 | 0.000 | 18.3 | 0.000 | 0.261 | 1.0 | 0.320 | 0.019 | 1.1 | 0.339 | | | | |
| Dorsal Attention A: Parietal Occipital 1 | 3.6 | 0.002 | 0.412 | 0.6 | 0.443 | 0.011 | 0.7 | 0.391 | 0.014 | 0.1 | 0.803 | 0.001 | 4.2 | 0.045 | 0.075 | 0.0 | 0.901 | 0.000 | 12.1 | 0.001 | 0.188 | 0.1 | 0.743 | 0.002 | 1.1 | 0.348 | | | | |
| Dorsal Attention A: Superior Parietal Lobule 1 | 4.5 | 0.000 | 0.465 | 0.0 | 0.866 | 0.001 | 0.7 | 0.399 | 0.014 | 0.1 | 0.786 | 0.001 | 1.2 | 0.288 | 0.022 | 0.0 | 0.896 | 0.000 | 20.0 | 0.000 | 0.278 | 0.3 | 0.558 | 0.007 | 2.0 | 0.129 | | | | |
| Dorsal Attention B: Post Central 1 | 3.6 | 0.002 | 0.408 | 3.1 | 0.085 | 0.056 | 0.1 | 0.775 | 0.002 | 0.5 | 0.495 | 0.009 | 6.0 | 0.018 | 0.104 | 0.1 | 0.705 | 0.003 | 11.2 | 0.002 | 0.177 | 1.6 | 0.216 | 0.029 | 0.9 | 0.430 | | | | |
| Dorsal Attention B: Post Central 2 | 3.3 | 0.003 | 0.392 | 0.1 | 0.753 | 0.002 | 0.0 | 0.909 | 0.000 | 1.6 | 0.216 | 0.029 | 0.8 | 0.372 | 0.015 | 0.2 | 0.626 | 0.005 | 15.8 | 0.000 | 0.233 | 0.3 | 0.601 | 0.005 | 1.5 | 0.235 | | | | |
| Dorsal Attention B: Frontal Eye Fields 1 | 4.0 | 0.001 | 0.437 | 0.2 | 0.646 | 0.004 | 0.0 | 0.928 | 0.000 | 0.5 | 0.475 | 0.010 | 1.2 | 0.270 | 0.023 | 0.5 | 0.466 | 0.010 | 17.9 | 0.000 | 0.256 | 0.0 | 0.878 | 0.000 | 2.0 | 0.125 | | | | |
| Salience Ventral Attention A: Parietal Operculum 1 | 3.4 | 0.002 | 0.398 | 1.1 | 0.297 | 0.021 | 0.0 | 0.953 | 0.000 | 0.2 | 0.622 | 0.005 | 5.0 | 0.029 | 0.088 | 0.1 | 0.788 | 0.001 | 4.9 | 0.032 | 0.086 | 3.8 | 0.056 | 0.068 | 0.1 | 0.954 | | | | |
| Salience Ventral Attention A: Insula: 1 | 4.4 | 0.000 | 0.456 | 1.2 | 0.281 | 0.022 | 0.4 | 0.548 | 0.007 | 0.5 | 0.489 | 0.009 | 6.5 | 0.013 | 0.112 | 0.0 | 0.936 | 0.000 | 12.6 | 0.001 | 0.195 | 5.0 | 0.030 | 0.087 | 0.3 | 0.858 | | | | |
| Salience Ventral Attention A: Parietal Medial 1 | 3.3 | 0.003 | 0.388 | 2.3 | 0.133 | 0.043 | 0.0 | 0.914 | 0.000 | 0.7 | 0.417 | 0.013 | 2.6 | 0.112 | 0.048 | 0.1 | 0.702 | 0.003 | 9.7 | 0.003 | 0.158 | 0.1 | 0.804 | 0.001 | 1.7 | 0.172 | | | | |
| Salience Ventral Attention A: Frontal Medial 1 | 4.8 | 0.000 | 0.481 | 0.2 | 0.625 | 0.005 | 0.6 | 0.457 | 0.011 | 0.5 | 0.501 | 0.009 | 2.0 | 0.168 | 0.036 | 0.0 | 0.886 | 0.000 | 21.9 | 0.000 | 0.297 | 2.1 | 0.155 | 0.039 | 1.5 | 0.233 | | | | |
| Salience Ventral Attention B: Inferior Parietal Lobule | 3.1 | 0.005 | 0.375 | 0.2 | 0.645 | 0.004 | 0.1 | 0.702 | 0.003 | 0.1 | 0.787 | 0.001 | 2.1 | 0.153 | 0.039 | 0.0 | 0.920 | 0.000 | 5.7 | 0.021 | 0.098 | 0.3 | 0.610 | 0.005 | 0.7 | 0.552 | | | | |
| Salience Ventral Attention B: Lateral Prefrontal Cort | 5.4 | 0.000 | 0.511 | 1.4 | 0.243 | 0.026 | 0.1 | 0.763 | 0.002 | 2.1 | 0.153 | 0.039 | 4.6 | 0.037 | 0.081 | 0.0 | 0.965 | 0.000 | 18.6 | 0.000 | 0.264 | 2.9 | 0.095 | 0.053 | 0.8 | 0.503 | | | | |
| Salience Ventral Attention B: Medial Posterior Prefro | 4.7 | 0.000 | 0.473 | 0.4 | 0.518 | 0.008 | 0.0 | 0.881 | 0.000 | 0.6 | 0.441 | 0.011 | 2.7 | 0.108 | 0.049 | 0.2 | 0.655 | 0.004 | 17.0 | 0.000 | 0.246 | 0.0 | 0.868 | 0.001 | 2.4 | 0.079 | | | | |
| LimbiC: B: Orbital Frontal Cortex 1 | 5.3 | 0.000 | 0.504 | 7.2 | 0.010 | 0.121 | 0.6 | 0.430 | 0.012 | 0.0 | 0.968 | 0.000 | 2.9 | 0.095 | 0.053 | 2.3 | 0.139 | 0.042 | 1.4 | 0.247 | 0.026 | 1.8 | 0.185 | 0.034 | 1.8 | 0.151 | | | | |
| LimbiC: A: Temporal Pole 1 | 7.1 | 0.000 | 0.578 | 0.2 | 0.660 | 0.004 | 0.5 | 0.482 | 0.010 | 0.0 | 0.984 | 0.000 | 0.4 | 0.510 | 0.008 | 1.2 | 0.271 | 0.023 | 12.2 | 0.001 | 0.190 | 6.1 | 0.017 | 0.104 | 2.8 | 0.050 | | | | |
| Control A: Intraparietal Sulcus 1 | 4.3 | 0.001 | 0.453 | 0.7 | 0.419 | 0.013 | 0.3 | 0.560 | 0.007 | 0.2 | 0.679 | 0.003 | 2.4 | 0.131 | 0.043 | 0.0 | 0.962 | 0.000 | 16.5 | 0.000 | 0.241 | 0.0 | 0.855 | 0.001 | 1.2 | 0.321 | | | | |
| Control A: Lateral Prefrontal Cortex 1 | 4.1 | 0.001 | 0.442 | 2.0 | 0.167 | 0.036 | 0.2 | 0.685 | 0.003 | 0.5 | 0.500 | 0.009 | 3.0 | 0.090 | 0.054 | 0.0 | 0.883 | 0.000 | 10.8 | 0.002 | 0.172 | 0.0 | 0.824 | 0.001 | 0.7 | 0.556 | | | | |
| Control A: Lateral Prefrontal Cortex 2 | 5.2 | 0.000 | 0.501 | 0.2 | 0.646 | 0.004 | 0.1 | 0.730 | 0.002 | 0.1 | 0.821 | 0.001 | 4.9 | 0.032 | 0.086 | 0.0 | 0.947 | 0.000 | 9.7 | 0.003 | 0.157 | 5.4 | 0.024 | 0.094 | 0.5 | 0.718 | | | | |
| Control B: Temporal 1 | 6.0 | 0.000 | 0.535 | 0.9 | 0.357 | 0.016 | 0.1 | 0.716 | 0.003 | 0.1 | 0.759 | 0.002 | 2.9 | 0.096 | 0.052 | 0.1 | 0.762 | 0.002 | 18.3 | 0.000 | 0.260 | 1.4 | 0.247 | 0.026 | 0.9 | 0.463 | | | | |
| Control B: inferior parietal lobule 1 | 3.3 | 0.003 | 0.385 | 0.2 | 0.686 | 0.003 | 0.6 | 0.424 | 0.012 | 0.0 | 0.842 | 0.001 | 1.8 | 0.181 | 0.034 | 0.0 | 0.987 | 0.000 | 11.4 | 0.001 | 0.179 | 0.0 | 0.904 | 0.000 | 0.7 | 0.577 | | | | |
| Control B: Lateral Prefrontal Cortexd 1 | 3.9 | 0.001 | 0.426 | 0.1 | 0.703 | 0.003 | 0.0 | 0.915 | 0.000 | 1.2 | 0.269 | 0.023 | 3.0 | 0.087 | 0.055 | 0.0 | 0.947 | 0.000 | 13.1 | 0.001 | 0.201 | 0.8 | 0.362 | 0.016 | 1.3 | 0.287 | | | | |
| Control B: Lateral Prefrontal Cortexv 1 | 6.6 | 0.000 | 0.558 | 7.4 | 0.009 | 0.125 | 0.0 | 0.874 | 0.000 | 0.0 | 0.934 | 0.000 | 2.2 | 0.143 | 0.041 | 1.9 | 0.169 | 0.036 | 12.4 | 0.001 | 0.192 | 0.0 | 0.997 | 0.000 | 1.3 | 0.300 | | | | |
| Control C: Cingulate Posterior 1 | 4.8 | 0.000 | 0.482 | 0.4 | 0.520 | 0.008 | 0.6 | 0.427 | 0.012 | 0.1 | 0.740 | 0.002 | 3.2 | 0.081 | 0.057 | 0.0 | 0.863 | 0.001 | 16.1 | 0.000 | 0.237 | 2.8 | 0.101 | 0.051 | 2.2 | 0.104 | | | | |
| Control C: Precuneus 1 | 2.8 | 0.009 | 0.350 | 0.5 | 0.489 | 0.009 | 0.1 | 0.741 | 0.002 | 1.0 | 0.321 | 0.019 | 3.5 | 0.068 | 0.063 | 0.4 | 0.528 | 0.008 | 8.3 | 0.006 | 0.137 | 0.0 | 0.867 | 0.001 | 1.5 | 0.235 | | | | |
| Default A: Inferior Parietal Lobule 1 | 4.0 | 0.001 | 0.432 | 0.7 | 0.416 | 0.013 | 0.1 | 0.747 | 0.002 | 0.2 | 0.653 | 0.004 | 4.8 | 0.032 | 0.085 | 0.0 | 0.908 | 0.000 | 12.2 | 0.001 | 0.190 | 0.5 | 0.465 | 0.010 | 0.2 | 0.873 | | | | |
| Default A: Dorsal Prefrontal Cortex 1 | 4.4 | 0.001 | 0.457 | 0.0 | 0.986 | 0.000 | 0.1 | 0.735 | 0.002 | 1.0 | 0.327 | 0.018 | 1.4 | 0.247 | 0.026 | 0.0 | 0.855 | 0.001 | 16.6 | 0.000 | 0.242 | 0.1 | 0.771 | 0.002 | 2.4 | 0.082 | | | | |
| Default A: Precuneus Posterior Cingulate Cortex 1 | 2.6 | 0.013 | 0.336 | 0.6 | 0.426 | 0.012 | 0.0 | 0.903 | 0.000 | 1.0 | 0.311 | 0.020 | 6.0 | 0.018 | 0.103 | 0.0 | 0.872 | 0.001 | 7.3 | 0.009 | 0.123 | 0.6 | 0.450 | 0.011 | 1.1 | 0.342 | | | | |
| Default A: Medial Prefrontal Cortex 1 | 4.9 | 0.000 | 0.484 | 3.0 | 0.091 | 0.054 | 0.4 | 0.517 | 0.008 | 0.4 | 0.512 | 0.008 | 2.4 | 0.125 | 0.045 | 0.4 | 0.516 | 0.008 | 13.5 | 0.001 | 0.206 | 0.5 | 0.482 | 0.010 | 1.4 | 0.251 | | | | |
| Default B: Dorsal Prefrontal Cortex 1 | 5.3 | 0.000 | 0.503 | 0.3 | 0.596 | 0.005 | 0.1 | 0.722 | 0.002 | 2.5 | 0.118 | 0.046 | 3.8 | 0.058 | 0.068 | 0.3 | 0.590 | 0.006 | 15.1 | 0.000 | 0.225 | 2.1 | 0.153 | 0.039 | 3.0 | 0.039 | 0.147 | 0.064 | 0.553 | 0.806 |
| Default B: Ventral Prefrontal Cortex 1 | 4.2 | 0.001 | 0.444 | 2.5 | 0.123 | 0.045 | 0.0 | 0.849 | 0.001 | 0.3 | 0.584 | 0.006 | 2.6 | 0.111 | 0.048 | 2.1 | 0.158 | 0.038 | 3.8 | 0.057 | 0.068 | 0.1 | 0.701 | 0.003 | 2.3 | 0.087 | | | | |
| Default B: Ventral Prefrontal Cortex 2 | 4.2 | 0.001 | 0.445 | 1.4 | 0.248 | 0.026 | 0.0 | 0.987 | 0.000 | 0.1 | 0.745 | 0.002 | 2.6 | 0.112 | 0.048 | 0.9 | 0.343 | 0.017 | 3.8 | 0.056 | 0.069 | 2.3 | 0.139 | 0.042 | 1.0 | 0.397 | | | | |
| Default C: Retro Superior Parietal Lobuleenial 1 | 2.2 | 0.035 | 0.295 | 0.2 | 0.635 | 0.004 | 0.1 | 0.769 | 0.002 | 1.4 | 0.249 | 0.025 | 3.8 | 0.058 | 0.067 | 0.1 | 0.765 | 0.002 | 7.9 | 0.007 | 0.132 | 0.1 | 0.760 | 0.002 | 0.9 | 0.456 | | | | |
| Default C: Parahippocampal Cortex 1 | 2.3 | 0.025 | 0.309 | 0.1 | 0.800 | 0.001 | 0.1 | 0.768 | 0.002 | 0.2 | 0.657 | 0.004 | 2.4 | 0.131 | 0.043 | 0.0 | 0.850 | 0.001 | 7.0 | 0.011 | 0.118 | 0.4 | 0.538 | 0.007 | 1.4 | 0.261 | | | | |
| Temporal Parietal 1 | 3.8 | 0.001 | 0.422 | 0.0 | 1.000 | 0.000 | 0.1 | 0.704 | 0.003 | 0.3 | 0.595 | 0.005 | 1.6 | 0.212 | 0.030 | 0.1 | 0.762 | 0.002 | 13.9 | 0.000 | 0.211 | 0.1 | 0.814 | 0.001 | 2.1 | 0.114 | | | | |
| Temporal Parietal 2 | 4.3 | 0.001 | 0.452 | 1.2 | 0.287 | 0.022 | 0.0 | 0.839 | 0.001 | 0.0 | 0.958 | 0.000 | 5.1 | 0.027 | 0.090 | 0.0 | 0.986 | 0.000 | 15.5 | 0.000 | 0.229 | 0.7 | 0.395 | 0.014 | 0.3 | 0.817 | | | | |
| Temporal Parietal 3 | 3.0 | 0.006 | 0.365 | 0.5 | 0.505 | 0.009 | 0.2 | 0.648 | 0.004 | 0.0 | 0.969 | 0.000 | 3.9 | 0.054 | 0.069 | 0.1 | 0.797 | 0.001 | 12.2 | 0.001 | 0.190 | 0.0 | 0.896 | 0.000 | 0.4 | 0.736 | | | | |

YIS = younger insulin sensitive; YIR = younger insulin resistant; OIS = older insulin sensitive; OIR = older insulin resistant

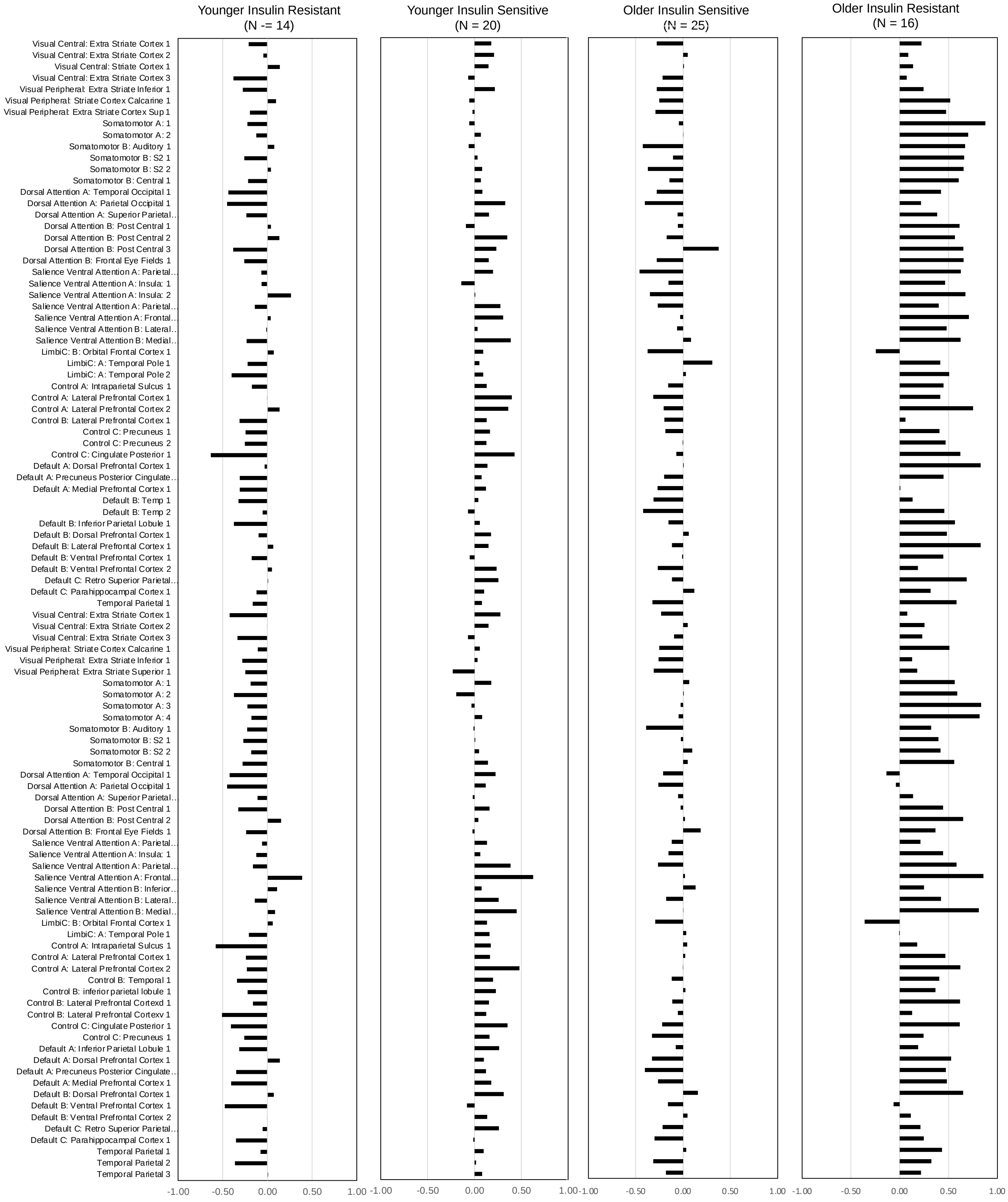

Figure S1. Partial correlation of regional CBF and $CMR_{GLC}$ controlling cortical thickness and systolic and diastolic blood pressure for four groups of participants based on age category and HOMA-IR levels: younger insulin sensitive; younger insulin resistant; older insulin sensitive; and older insulin resistant. The data for each group are also plotted on the brain surface in Figure 4 in main document and the GLMs are in Table S3.

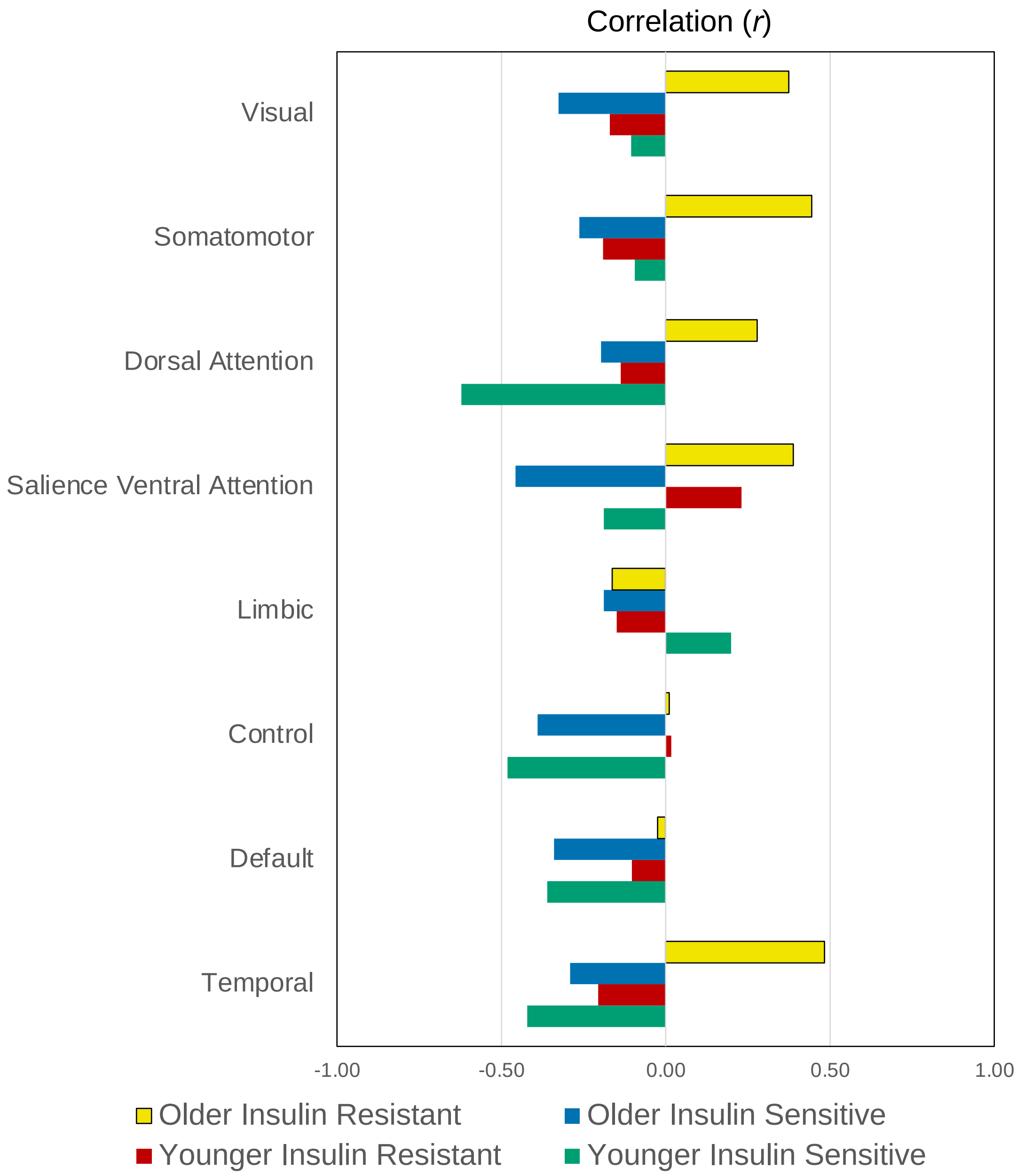

Figure S2. Partial correlation of network cerebral blood flow (CBF) and glucose metabolism ($CMR_{GLC}$) controlling for cortical thickness blood pressure, resting heart rate, BMI, sex and years of education. Four groups based on age category and HOMA-IR levels: younger insulin sensitive; younger insulin resistant; older insulin sensitive; and older insulin resistant.

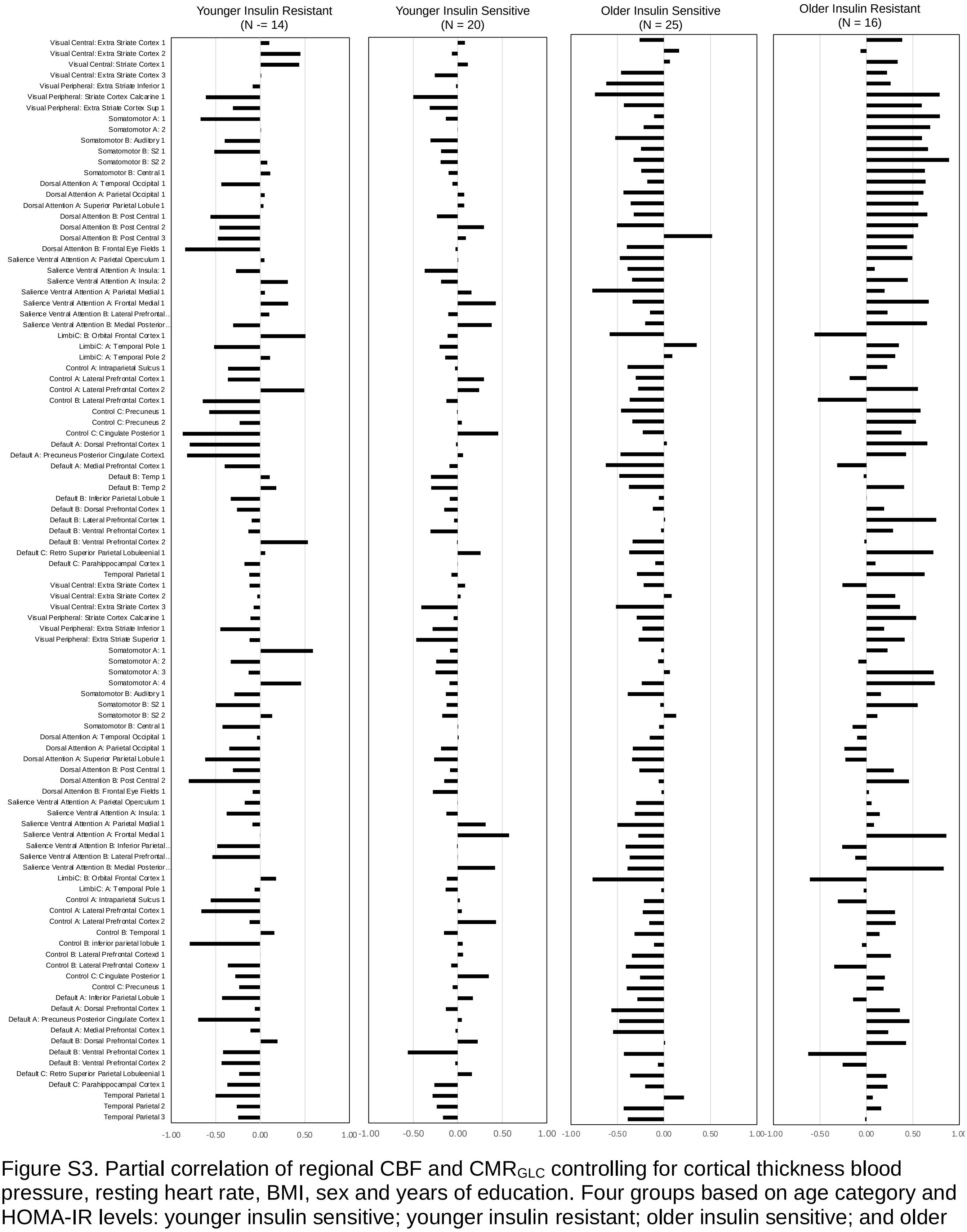

Figure S3. Partial correlation of regional CBF and $CMR_{GLC}$ controlling for cortical thickness blood pressure, resting heart rate, BMI, sex and years of education. Four groups based on age category and HOMA-IR levels: younger insulin sensitive; younger insulin resistant; older insulin sensitive; and older insulin resistant
